# Partial repair causes permanent defects in papillary structure and function after reversal of urinary obstruction

**DOI:** 10.1101/2024.09.11.612436

**Authors:** Thitinee Vanichapol, Alex Gonzalez, Rachel Delgado, Maya Brewer, Kelly A. Clouthier, Anna Menshikh, William E. Snyder, Teebro Rahman, Veronika Sander, Haichun Yang, Alan Davidson, Mark de Caestecker

## Abstract

Urinary obstruction causes injury to the renal papilla and leads to defects in the ability to concentrate urine which predisposes to progressive kidney injury. However, the regenerative capacity of the papilla after reversal of obstruction is poorly understood. To address this, we developed a mouse model of reversible urinary obstruction which is characterized by extensive papillary injury, followed by a robust regeneration response and complete histological recovery over a 3- month period. However, these mice have a pronounced defect in urinary concentrating capacity. We now show that this is due to permanent changes in the composition, organization, and transcriptional signatures of epithelial, endothelial, and interstitial cell lineages in the papilla. There are persistent inflammatory responses that are also seen in patients with renal stone disease but are associated with cell-specific adaptive responses to the increasingly hypoxic environment of the papilla after reversal of obstruction. Taken together, our analysis of a new model of reversible urinary obstruction reveals that partial repair leads to permanent changes in the structure and function of all of the major cellular compartments in the papilla that include both shared and distinct responses to different types of renal papillary injury in humans and mice.

**Summary:** Partial repair after reversal of urinary obstruction leads to permanent changes in structure and function of all major cellular compartments in the renal papilla

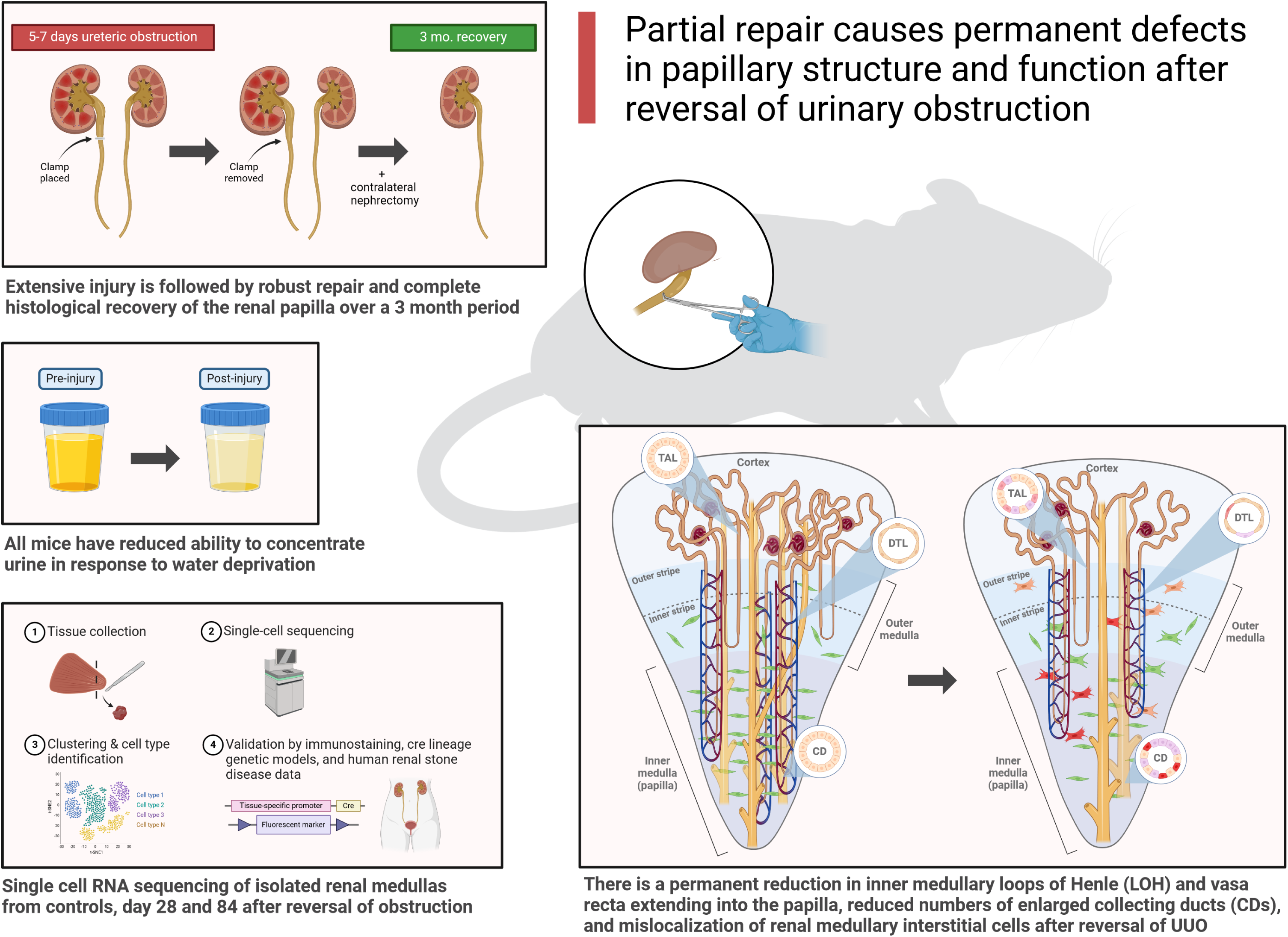

## Introduction

The ability to concentrate urine in response to water deprivation depends on the ability to generate axial osmotic gradients in the renal papilla.^1^ This in turn is dependent on maintaining the precise anatomic integrity of the papilla, with correct positioning with correct positioning of thin limbs of the loops of henle (LOH), collecting ducts (CDs), and specialized capillary networks, the vasa recta, which supply blood to the papilla, and play a central role in counter current ion and water exchange that is required to generate these axial osmotic gradients. ^1,2^ While there is an extensive literature on normal structure and function of the renal papilla, much less is known about how it is affected by injury and disease.

Urinary obstruction is a common cause of renal papillary dysfunction.^3^ It is often asymptomatic and diagnosed late after prolonged obstruction.^4^ Treatment requires reversal of the obstruction, but patients are at increased risk of a progressive decline in renal function despite early treatment.^5,6^ In addition to this decline in renal function, determined by a reduction in glomerular filtration rates (GFR), these patients have long-term defects in their ability to concentrate urine.^7^ Split renal function studies in dogs with >2 weeks of unilateral ureteric obstruction (UUO) have shown that the affected kidney is unable to concentrate urine up to 18 months after reversal.^8^ This indicates that the effects of obstruction on urinary concentrating capacity may be permanent. Since failure to concentrate urine predisposes patients to dehydration and recurrent acute kidney injury (AKI), this may contribute to progressive decline in renal function in these patients.

The renal medulla extends from the corticomedullary junction into the outer medulla, and transitions abruptly from the inner stripe of the outer medulla (ISOM) into the papilla, or inner medulla, which is characterized by the presence of specialized inner medullary CDs, TLs, and vasa recta (see summary Fig. 8).^9^ Urine drains directly from the deep inner medullary CDs into the renal pelvis. The papilla is particularly sensitive to damage caused by urinary tract obstruction, in part due to the increase in pressure in the renal pelvis,^10^ but also because of a profound reduction in inner medullary blood flow. ^10,11^ As the papilla already has low tissue oxygenation,^12^ this renders it particularly susceptible to ischemic damage after urinary obstruction, which when severe can cause papillary necrosis and radiological evidence of papillary flattening.^13,14^ This is recapitulated in mice with irreversible UUO, in which papillary compression begins after 3 days of obstruction.^15^ In rats, 2 weeks of partial ureteric obstruction causes a marked shortening of renal papillae associated with a reduction in the number of long LOH and inner medullary CDs. ^16^ However, no studies have evaluated whether these anatomical changes to the renal papilla are reversible, and if so, whether there are structural defects resulting from incomplete or failed repair, and whether these are sufficient to prevent long-term recovery of inner medullary function.

The pathophysiological changes that occur in rodents are likely to reflect similar changes in humans since, other than size and the lobulated architecture of the human kidney, the cellular composition and organization of the renal medulla is highly conserved between rodents and humans.^17^ Irreversible UUO models have been widely used, but do not recapitulate the clinical situation in which patients undergo reversal of obstruction soon after diagnosis. In addition, they do not allow assessment of regenerative responses after reversal. Reversal of ureteric obstruction in rats lasting less than 24 hours shows that there is a persistent defect in the ability to concentrate the urine up to 60 days after reversal associated with downregulation of water channels in inner medullary LOH and CDs. ^18,19^ However, these studies do not recapitulate the effect of prolonged urinary tract obstruction in which there is structural damage to the papilla. Reversible UUO models have been developed that could study this, but these are more technically challenging and have not been widely adopted ^20^ A variety of approaches have been used, including sequential placement of clamps along the ureter; ^20,21^ UUO followed by bladder re-implantation of the ureter;^22–25^ and the use non-damaging clamps and tubing to obstruct the ureter.^26–28^ Long-term outcome studies in rats and mice have shown improved renal function and reduced renal cortical fibrosis after reversal of obstruction, but only partial recovery if the obstruction lasts longer than 2 days.^20–23,26,28–33^ However, aside from the short-term rat studies described above,^18,19^ there have been no long-term studies on the effects of reversing more prolonged ureteric obstruction on renal papillary structure and function.

Highly dimensional single cell profiling which began with the advent of single cell RNA sequencing (scRNA-Seq) technologies,^34^ has led to a number of reports describing changes that occur to the dominant cell populations collected from kidneys before and after AKI.^35^ This has led to the observation that some proximal tubular epithelial cells (PTECs), which undergo de-differentiation and proliferation after injury, do not fully repair and become fixed in a permanent, senescent, pro-inflammatory state that is thought to drive local inflammation, fibrosis, and chronic kidney disease (CKD).^36,37^ These so called “maladaptive” or “failed repair” PTECs (FR-PTECs) have been identified using different experimental models, including mice with ischemia reperfusion induced AKI (IRI-AKI) and irreversible UUO, ^36–39^ as well as human AKI and CKD. ^40–42^ More recently, genome-wide, single cell ATAC-Seq studies have shown that there is widespread epigenetic reprogramming of both FR-PTECs and PTECs that have undergone adaptive repair after injury, which is thought to maintain these cells in permanent maladaptive and dedifferentiated states.^40,43,44^ However, there has been remarkably little published data using single cell profiling to evaluate the fate of renal papillary cell populations in these disease states. One of the challenges is that that renal medullary cells are not well represented in whole kidney preparations used for many of these scRNA-Seq studies.^45^ For this reason, it is unclear whether the same paradigm of maladaptive repair is operative in the renal medulla. This is important because the papilla has a uniquely hypoxic and hyperosmolar environments that inner medullary cell populations need to adapt to are not found elsewhere in the kidney,^46^ so it is likely that there will be distinct requirements for cell survival and recovery after reversal of prolonged urinary obstruction.

Here we describe a new mouse model of reversible UUO in which we evaluated renal function up to three months after reversal of prolonged UUO by removing the contralateral kidney. We show that these mice have a permanent reduction in GFR associated with a marked and irreversible defect in urinary concentrating capacity. Using scRNA-Seq of isolated renal medullas with orthogonal validation by cell lineage and quantitative immunostaining studies, we show that despite robust regenerative responses that restore overall dimensions of the renal papilla after reversal of UUO, there are permanent changes to all of the major cellular compartments that would be expected to have a profound effects on renal medullary function. This includes changes to the numbers and organization of inner medullary LOH, CDs, and vasa recta, as well as reduction and mislocalization of renal medullary interstitial cells,^47,48^ which are known to regulate urinary concentrating capacity in the papilla.^49,50^ We also show that persistent proinflammatory responses seen in FR-PTECs after AKI, are also seen in inner medullary LOH and CD cells after reversal of UUO, and that these responses are conserved in renal papillary CDs from patients with renal stone disease. These results indicate that there is a common, long-term chronic injury response to different types of renal papillary injury in both humans and mice.

## Results

### Long-term decrease in renal function and urinary concentrating capacity after reversal of UUO

To explore the effects of prolonged ureteric obstruction on the kidney, a light vascular clamp was placed on the proximal left ureter of male BALB/c mice and removed after different time periods to induce reversible UUO. A contralateral nephrectomy (Nx) was performed 10 days later to evaluate renal function (Fig 1A). After 7 days of UUO, 30-50% of mice died within 2 days of the Nx, and 80% died after 8 days of UUO. Amongst those surviving 7 days of UUO, BUN peaked at maximum of ∼50-60mg/dl 1 day after Nx, gradually returning to baseline values (Fig. 1B/C, S. Fig. 1). There was a reduction in transdermal GFR (tGFR) at 28 days that partially improved but was still significantly reduced after 84 days (Fig. 1D/E). This improvement over time may in part be due to compensatory hypertrophy of surviving nephrons. However, there was also a marked reduction in urinary concentrating capacity at 28 days that was still present 84 days after reversal of UUO (Fig. 1F/G). This suggests that there is a persistent defect in renal medullary function that is independent of the reduction in tGFR seen in mice after reversal of UUO. Survivable reversible UUO clamp times were different in different mouse strains (5- and 6- days of UUO in mice on two of the different genetic backgrounds used in these studies), but all showed a similar reduction in tGFR and marked decrease in urinary concentrating capacity, 84 days after reversal (see S. Fig. 7/15).

**Figure 1.**
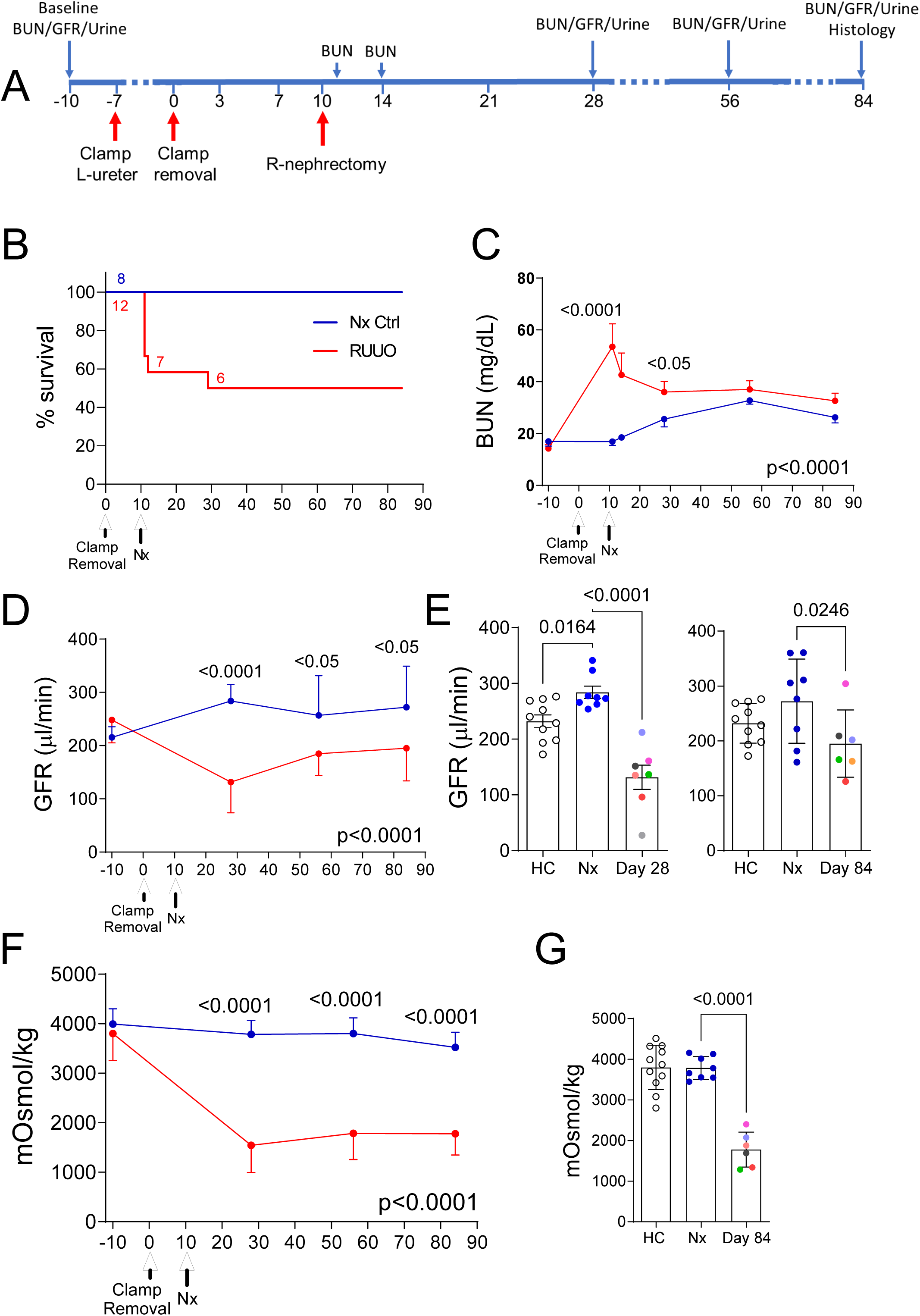
Long-term decrease in renal function and urinary concentrating capacity after reversible UUO. Male BALB/c mice underwent R-UUO and contralateral nephrectomy, or nephrectomy alone (Nx). A, Study design; B, Survival, numbers of mice indicated at each time point; C, BUN time course; D, tGFR time course; E, tGFR at day 28 and 84 after R-UUO; F, Urinary osmolality after 18hr water restriction; G, Urinary osmolality after 18hr water restriction at 84 days; C/D/F, data shown mean +/-SD. 2-way ANOVA for between group comparisons. E/G, individual data points shown with mean +/-SEM. R-UUO datapoints are color coded to show the relationship between tGFR and urine osmolality in individual mice. 1-way ANOVA Nx vs. HC and R-UUO. If p<0.05, q values shown for between group comparisons corrected for repeat testing.

### Early renal papillary injury, repair, and remodeling after reversal of UUO

Renal histology over the first 28 days after reversal of UUO showed tubular damage and increased peritubular cellularity throughout the renal medulla (Fig. 2A). At early time points after reversal, the papilla was shortened. From days 0 to 3, more than 50% of mice had evidence of papillary necrosis with focal areas in the distal inner medulla with complete loss of nuclei and empty tubular basement membrane structures indicating large, denuded areas of tubular epithelium. Over this time period, papillary urothelium, which is normally a single layer of cells on the outside of the papilla, increased from 1 to 3 or more cell layers in thickness. By day 7 there was evidence of tubular repair, with repopulation of denuded tubular basement membranes with epithelium that gradually increased over the next 3 weeks so that most of the tubular basement membranes structures had been repopulated with epithelium by 28 days. This active process of repair was associated with increased expression of the cell cycle and regenerative markers Ki67 and Sox9, which peaked 7 days after reversal of UUO (Fig. 2B-E).

**Figure 2.**
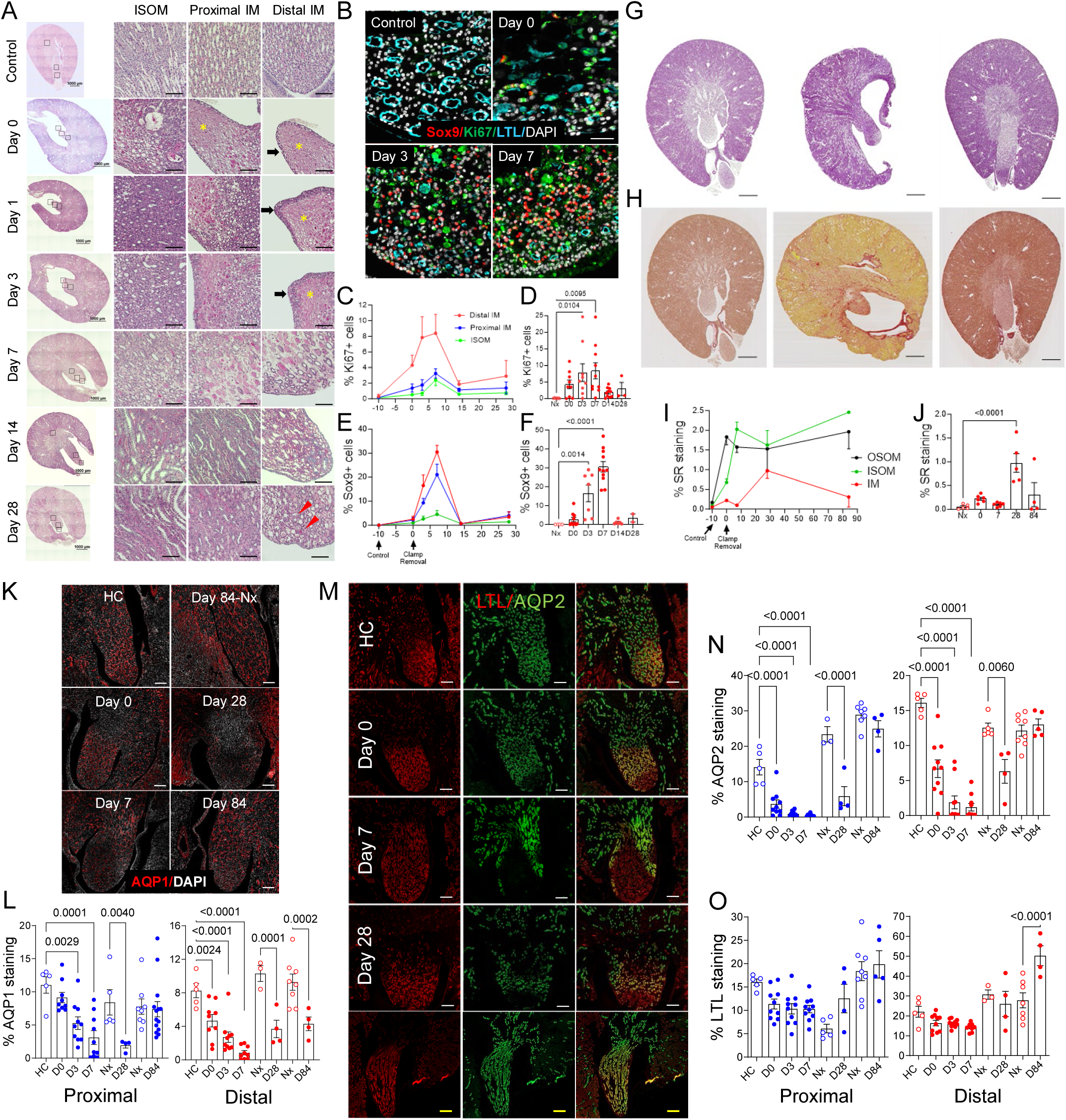
Long-term changes to renal papillary epithelial cells despite robust repair and remodeling after reversible UUO. Kidneys collected at different time points after R-UUO. A, Periodic acid-Schiff (PAS) staining, whole kidneys scale bars=1mm, inner stripe of the outer medulla (ISOM), proximal and distal inner medulla (IM), scale bars=100uM. *indicates papillary necrosis; black arrows show thickening of papillary urothelium; red arrow heads indicate expansion of interstitial myofibroblast-like cells; B, Ki67, Sox9 and LTL in the distal IM at different time points after R-UUO. Scale bars=50uM; C/E, Time course of Ki67+ and Sox9+ cells (% of total nuclei). D/F, Ki67+ and Sox9+ cells in the distal IM. G/H, PAS and Sirius red staining in Nx control, 28 and 84 days after R-UUO. Scale bars=1mM; I/J, Quantification of Sirius red staining. Time course in the OSOM, the ISOM, and the IM (I), and in the IM only (J). K/L, *Persistent loss of AQP1 expression in IM after R-UUO.* K, AQP1 staining of the IM in healthy controls (HC), Nx, and at different time points after R-UUO. Scale bars=200uM. L, Quantification of AQP1 staining in the proximal and distal IM time course. Healthy controls (HC), day 28 and 84 Nx shown. M-O, *Transient loss of AQP2 with increased LTL staining of IM CDs after R-UUO.* AQP2 and LTL staining of IM CDs in HCs, and at different time points after R-UUO. While scale bars=200uM, yellow scale bars=400uM; N/O, Quantification of AQP2 and LTL staining as the % of the area in the proximal and distal IM over time after R-UUO. Individual data points shown. Results expressed as means +/- SEM. 1-way ANOVA, HC or Nx vs. R-UUO time points. If p<0.05, q values are shown for between group comparisons corrected for repeat testing.

At day 28 after reversal of UUO, most of the inner medullary epithelium had regenerated but the interstitial space was disorganized and expanded with the appearance of myofibroblast-like interstitial cells (Fig. 2A). There was also tubular injury and disorganized repair in the proximal inner medulla, and in the inner stripe of the outer medulla (ISOM), which lies just above the papilla. However, by day 84 the papilla had recovered its full length, and on PAS-stained sections, the ISOM and inner medulla were indistinguishable from controls (Fig. 2G, S. Fig. 2). Staining for collagen deposition with Sirius red showed that while there was an early increase in fibrosis in the outer medulla that persisted 84 days after reversal, collagen deposition in the inner medulla only increased after 7 days, was maximal at 28 days, and decreased by day 84 (Fig. 2H-J, S. Fig. 3). These findings suggest that despite robust early repair after reversal of UUO, there must be defects in the fine structure and function of the inner medulla that are not apparent on standard histological staining in order to account for the long-term defects in urinary concentrating capacity after reversal of UUO.

### Permanent loss of AQP1 and transient loss of AQP2 expression in the renal papilla

To evaluate this further, we examined markers of differentiated inner medullary LOH and CD epithelia by immunostaining. We used AQP1 antibodies, which stain descending thin limb (DTL) LOH and descending vasa recta (DVR) in the papilla. We found a marked reduction in AQP1 staining throughout the papilla in the first 7 days after reversal (Fig. 2K/L). AQP1 staining was still decreased throughout the inner medulla by day 28 and was decreased in the distal but not the proximal inner medulla 84 days after reversal of UUO. This persistent reduction in AQP1 in the distal inner medulla could result from a reduction in AQP1 expression in dedifferentiated long LOHs, a reduction in numbers of long LOHs extending into the inner medulla, and/or a reduction AQP1 positive DVRs. To evaluate inner medullary CDs, we first assessed AQP2 immunostaining, which labels differentiated principal cell CDs, and Lotus Tetroglobinous Lectin (LTL), which in addition to staining PTECs, provides a marker of inner medullary, but not cortical or outer medullary, CDs (S. Figure 4), ^51^ as well as V-ATPase positive intercalated cells (IC) in cortical and outer medullary CDs (S. Figure 5). We found an initial reduction in AQP2 staining in proximal and distal inner medulla that decreased for 7 days after reversal of UUO (Fig. 2M/N). Reduced AQP2 staining persisted at 28 days but was indistinguishable from controls after 84 days. In contrast to AQP2, there was a slight, but non-significant, reduction in LTL staining throughout the inner medulla, but no reduction by 28 days, and by day 84, the surface area staining with LTL in the distal inner medulla was actually increased (Fig. 3M/O). Restoration of AQP2 staining suggests that CD epithelium has regenerated and differentiated, while expansion in LTL staining 84 days after reversal of UUO could be due to an increase in density of CD epithelium, an increase in the surface area of existing CD epithelium, and/or non-specific staining of other structures in the papilla after reversal of prolonged UUO.

**Figure 3.**
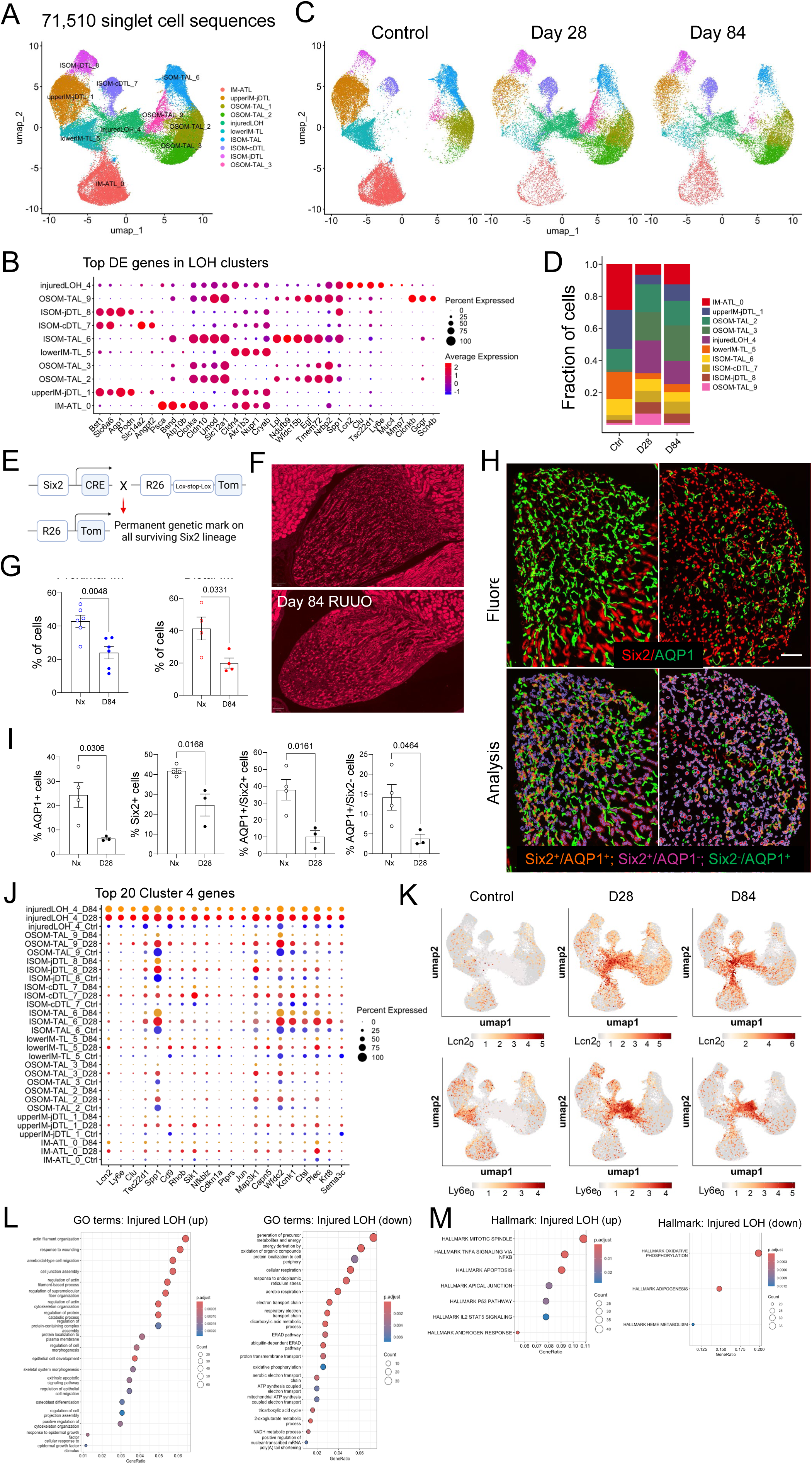
Permanent reduction in inner medullary loops of Henle is associated with expansion of injured cells. A/C, Re-clustering of LOH cells identified 10 cell populations, including injured LOH cells (cluster 4), and 3 clusters of IM LOH cells in the combined data (A), and at different time points after R-UUO (C); B, Top DEGs in the LOH cell populations; D, Fraction of LOH cells at different time points after R-UUO; E, Genetics showing Six2 Cre-dependent labeling with tdTomato of all IM LOH cells; F, Fluorescence images showing Six2 lineages in controls and 84 days after R-UUO, scale bars=200um; G, Quantification of Six2 lineage cells in the proximal and distal IM; H, Images and overlay of AQP1 and Six2 lineage staining of the distal IM of controls and 28 days after R-UUO; I, Quantification of AQP1+ and Six2 lineage cells in the distal IM. (% of cells). Individual data points, means +/- SEM. T- test, p values shown. J, Clustering of the top 20 cluster 4 DEGs in other LOH cell populations after R- UUO; K, UMAPs showing expression distribution of selected injury markers Lcn2 and Ly6e in controls, day 28 and 84 after R-UUO; L/M, Gene set enrichment analysis of the top 20 injured cluster 4 DEGs, showing upregulated and downregulated GO terms (L), and Hallmark terms (M).

### Permanent reduction in inner medullary CD and LOH populations and expansion of de-differentiated injured cells

To address these questions, and to determine whether reversal of UUO results in changes to other papillary cell populations, we performed scRNA-Seq on isolated renal medullas dissected from multiple male BALB/c mouse kidneys at 28 (n=12) and 84 days (n=6) after reversal of UUO and compared these with healthy controls (n=12). We identified 181,255 singlet cell sequences, including 7 major cell types (S. Fig. 6A/B). Immunostaining of isolated medullas that were dissected in parallel showed that while the papilla was intact in all samples as determined by LTL staining of inner medullary CDs, there was variable staining ISOM cells, as determined by the characteristically strong Uromodulin (UMOD) staining of densely packed TALs in the ISOM, and only occasional staining of LTL positive PTECs in the outer stripe of the outer medulla (OSOM), which lies between the ISOM and the renal cortex, in some samples (S. Fig. 6C). For this reason, while it is possible to use this scRNA-Seq data to interpret changes in numbers of all of the inner medullary cell populations, changes in numbers of TAL and PTECs in the outer medulla have to be interpreted with caution.

We used UMAP plots to visualize the cell populations in these samples and used published anchor genes, some of which provide information about the spatial localization of clusters in the inner vs. outer medulla (see Supplemental Methods 1), ^45,52,53^ to assign identities to each cluster. This enabled us to identify 25 cell clusters, including inner and outer medullary CD principal cells (referred to as CD cells here), LOH, endothelial cells, fibroblasts, and immune and intercalated cell (IC) clusters (S. Fig.6D/E, see S. Table 2 for differentially expressed genes (DEGs) in these “Big UMAP” clusters). There was a long-term reduction in the proportion of all inner medullary CD (“deep IM CD” and “IM CD”), LOH (“DTL”, “ATL” (ascending thin limb) and “IM TL” (thin limb), and endothelial cells (“Endo VR/papilla”) after reversal of UUO (S. Fig. 6F/G). We were unable to distinguish inner and outer medullary fibroblast clusters on this initial analysis, but there was a reduction in renal medullary fibroblasts associated with expansion of myofibroblasts at day 84. There was also an expansion in populations of less differentiated CD and LOH clusters with features of injured cell populations. This included a reduction in expression of classical ATL, DTL, TAL, and CD markers, and increased expression of injury markers such as *Lcn2, Spp1, Ccn2, Clu,* and *Igfbp2* (S. Fig. 6E/H). These injured and partially de-differentiated LOH and CD clusters are reminiscent of the maladaptive FR-PTECs described after IRI-AKI^36–39^, irreversible-UUO, ^38^ and in patients with CKD, ^41^ and sepsis-associated AKI.^42^ Since we had limited data on the spatial organization of these renal medullary cell populations after reversal of UUO, we performed a more focused analysis of papillary LOH, endothelial, fibroblasts, and CD clusters.

### Permanent reduction of inner medullary loops of Henle

Re-clustering of LOH cells and interrogation of their DEGs with anchor genes to assign identities, ^52^ identified 3 populations of papillary thin limb cells, the “IM ATL cluster 0”, “upper IM jDTL cluster 1” (which represent DTL cells from juxtamedullary nephrons), and “lower IM TL cluster 5” cells (Fig. 3A/B, see S. Table 2 for LOH cluster DEGs). There was a marked reduction in all 3 papillary thin limb cell populations that persisted 84 days after reversal of UUO (Fig. 3C/D). To validate these findings, we performed lineage analysis using *Six2 Cre* mice crossed with *R26R LSL tdTomato* mice to genetically label all nephronic epithelium, but not CD, including papillary LOH cells, with the tdTomato fluorophore (Fig. 3E).^54^ After optimizing the UUO times to allow > 50% survival after Nx (5 days), these mice showed a similar long-term reduction in tGFR and urinary concentrating capacity to that seen in mice used for our scRNA-Seq studies (S. Fig. 7). We then quantified tdTomato expression to assess the density of papillary structures irrespective of their state of injury and differentiation. There was a generalized reduction in thin limb cells throughout the papilla 84 days after reversal (Fig. 3F/G). Since we have shown that reversible UUO results in a persistent reduction in AQP1 staining in the papilla (see Fig. 2K/L), we took advantage of Six2 lineage staining to determine whether this reduction in AQP1 also resulted from a reduction in Six2^-^, AQP1^+^ DVRs. Overlaying the Six2 lineage marker with AQP1 showed that the reduction in both AQP1^+^ and Six2^+^ cells in the distal inner medulla (Fig. 3H/I) was associated with fewer Six2^-^,AQP1^+^ DVRs throughout the inner medulla 28 days after reversal (Fig. 3H/I, S. Fig. 8). These data indicate there is a generalized reduction in LOH thin limb and descending vasa recta throughout the papilla after reversal of prolonged UUO.

### Injured LOH cells are derived from all of the LOH cell clusters and have features of failed repair cells

There was also an expansion of “injured LOH cluster 4” cells after reversal of UUO. These cells express the injury markers including *Lcn2, Spp1, Clu,* and *Mmp7*, and lower levels of segment-specific LOH markers (Fig. 3B, see S. Table 2 for DEGs in the “Injured LOH cluster 4”), consistent with these cells being partially dedifferentiated cells with persistent cellular injury. Clustering of the top 20 DEGs in cluster 4 cells in the different LOH sub-clusters in controls and day 28 and 84 after reversal of UUO shows that while some, such as *Sik1, Wfdc2, Kcnk1, Ctsl, Plec,* and *Krt8*, are also increased in the other LOH cell populations, the majority are either only upregulated in “injured LOH cells” or are expressed in other LOH cell clusters but not increased with injury (Fig. 3J/K). More detailed analysis indicates that there are four sub-clusters of “injured LOH cluster 4” cells (S. Fig. 9A). Unbiased clustering of differentially expressed genes (DEGs) shows that subclusters 4-1 and 4-3 are more closely related to “DTL, ATL, and IM TL clusters 0, 1, 5, 6, and 7”, while subclusters 4-0 and 4-2 are more closely related to “OSOM and ISOM TAL clusters 2, 3, 6 and 9” (S. Fig 9B, see Table 2 for sub-cluster DEGs). This can be visualized by evaluating gene expression profiles within the LOH clusters at different time points after reversal of UUO (S. Fig. 9C).

For example, *Aqp1*, which is expressed by “DTL clusters 1 and 8” in controls, is expressed in the “injured LOH cluster 4-3”, while *Psca*, which is expressed by “IM-ATL cluster 0”, is expressed in “LOH cluster 4-0” after reversal of UUO (S. Fig. 9C). Likewise, the pan-TAL markers, *Slc12a1*, is expressed in “injured LOH 4-0 and 4-2” clusters after reversal of UUO. This indicates that “injured LOH cluster 4” cells are derived from all of the LOH cell types in the inner and outer medulla.

Gene set enrichment analysis (GSEA) of the top 20 upregulated DEGs in “injured LOH cluster 4” cells (Fig. 3J), shows enrichment for Gene Ontology (GO) terms for processes involved in cell migration and wound healing, and downregulation of pathways involved in respiration, metabolite precursors, oxidative reactions (Fig. 3L, S. Table 3). This suggests that these cells are metabolically quiescent but are attempting to undergo repair. GSEA for Hallmark terms shows enrichment for apoptosis and TNFα via Nfκb signaling, and a downregulation in genes involved in oxidative phosphorylation (Fig. 3M, S. Table 3), which are also characteristic of FR-PTECs after AKI.^36,37^ There was also highly significant overlap between FR-PTEC DEGs after irreversible UUO,^38^ and “injured LOH” DEGs after reversal of UUO (S. Fig. 10A-E, for DEGs in FR-PTECs used for these comparisons see S. Table 4). ^38^ These findings indicate that similar pathways of cell migration, wound healing, and inflammation are also activated in FR-PTECs after irreversible UUO.

### Minor long-term increase in inflammatory macrophages in the papilla

There is an expansion of inflammatory macrophages around failed repair cells after IRI-AKI.^36^ However, our scRNA-Seq data showed only a minor increase in renal medullary inflammatory cells 28 and 84 days after reversal of UUO (see S. Fig. 6D/F). While the numbers of singlet cell sequences was relatively low (3266 cells), we identified three populations: “T cells”, and two macrophage clusters (“Mac 0” and “Mac 1”), with a decrease in the number of “T cells”, and an increase the “Mac 1” cluster after reversal of UUO (S. Fig 11A-D, see S. Table 2 for DEGs in the immune cell clusters). GSEA of DEGs upregulated in these cells showed that GO term pathways are activated for cell migration, adaptive immunity, and phagocytic responses in “Mac 0” and “Mac 1” clusters. Hallmark terms show inflammatory responses, particularly in the “Mac 1” cluster, with upregulation of genes in the TNFα via NfκB, IL2/Stat, and inflammatory response categories (S. Fig. 11E/F). However, more detailed analysis using datasets for pro-inflammatory (M1 activated) and anti-inflammatory (M2 activated) macrophage gene sets, which have been used to characterize renal macrophage populations after AKI,^55–57^ showed no evidence of proinflammatory M1 type, or anti-inflammatory M2 type responses (S. Fig. 11G). Because immune cells maybe underrepresented in scRNA-Seq studies, we also evaluated F4/80 antibody staining, a marker of renal macrophages.

While there was an early increase in F4/80 staining throughout the kidney, F4/80 staining decreased over time (S. Fig. 11H-J) but was still increased in the cortex, outer medulla, and in the proximal but not distal inner medulla 84 days after reversal of UUO (S. Fig 11J). These data indicate that despite activation of inflammatory signaling in injured LOH cells, there is only a minor long-term increase in inflammatory macrophages in the inner medulla after reversal of UUO.

### Permanent reduction in vasa recta throughout the renal papilla

Endothelial cell re-clustering identified 9 clusters, including “papillary AVR and DVR clusters 1 and 6”, other “AVR and DVR clusters 0 and 2”, and “capillary angiogenic clusters 3 and 7” (Fig. 4A/B, see S. Table 2 for DEGs in the endothelial cell clusters). “Papillary AVR and DVR” clusters decreased 28 and 84 days after reversal, while both capillary angiogenic clusters increased (Fig. 4C/D). GSEA of the top 20 DEGs upregulated in these clusters showed that “papillary AVR and DVR clusters 1 and 6” were enriched for GO terms involved in cell migration, and “papillary AVR cluster 1” showed evidence of an ER stress response (Fig. 4E). GO terms for angiogenesis and cell migration were also increased in “capillary angiogenesis clusters 3 and 7” (Fig. 4F), suggesting that these cells may be attempting to repair after reversal of UUO. However, quantification of endothelial cell staining with CD31 antibodies showed that there was a reduction in the density of endothelial cells throughout the inner medulla 84 days after reversal of UUO (Fig. 4G/H). These findings are consistent with our data showing a reduction in Six2 lineage negative, AQP1^+^ DVR cells throughout the inner medulla 28 days after reversal (see Fig. 4H/I, S. Fig. 9), and with our scRNA-Seq data showing a persistent reduction in “papillary AVR and DVR” cells after reversal of UUO. These data indicate that despite attempts at endothelial cell repair, there is a permanent reduction in the density of inner medullary peritubular capillaries after reversal of prolonged UUO. This is notable since peritubular capillary rarefaction may give rise to worsening hypoxia, particularly in the distal inner medulla, which may exacerbate cellular injury after reversal of UUO.

**Figure 4.**
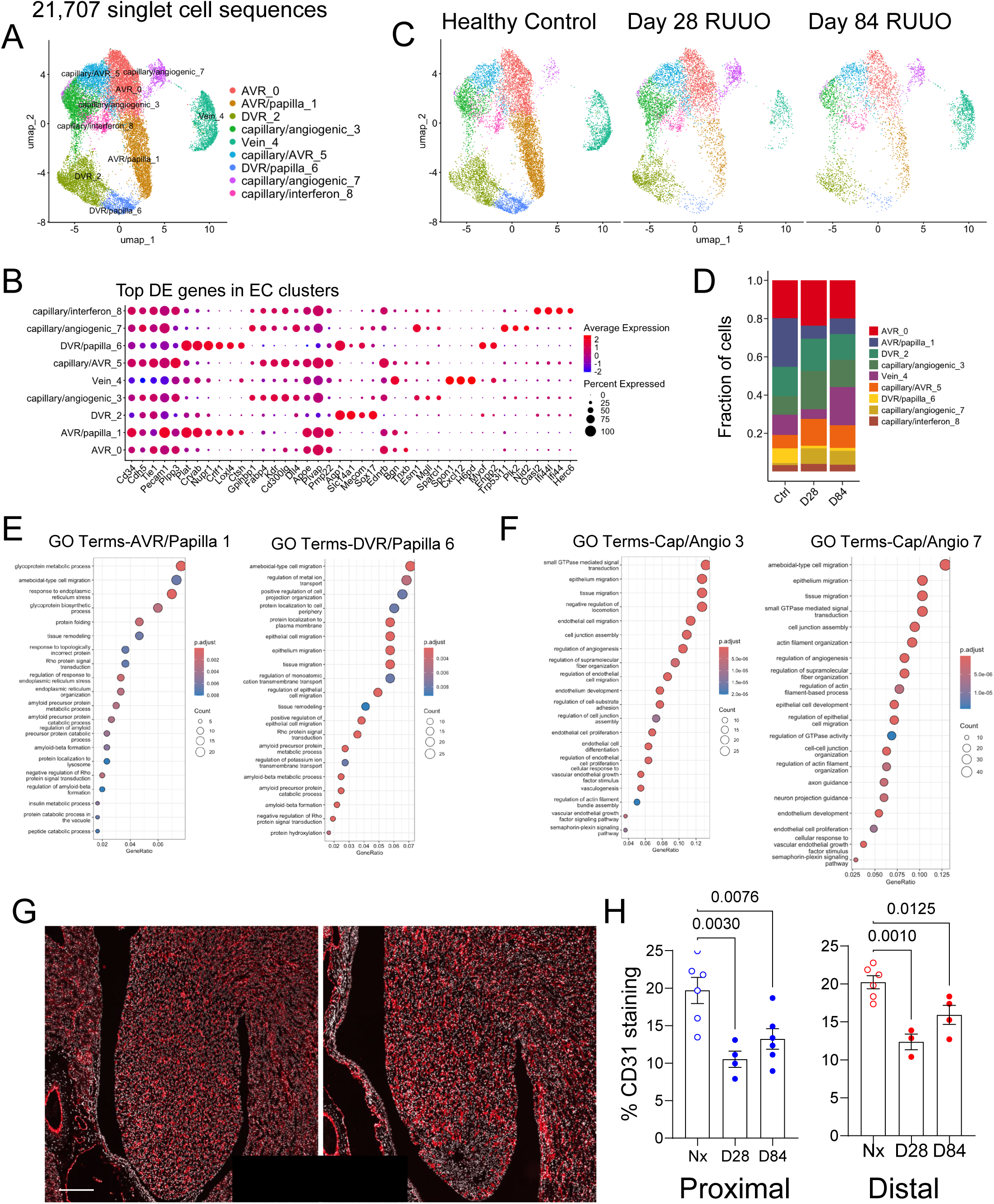
Reduced density of vasa recta throughout the renal papilla. A, Re-clustering of renal medullary endothelial cells (ECs) identified 9 cell populations, including papillary AVR and DVR clusters localized in the IM, and two capillary angiogenic clusters in the combined data (A), and at different time points after R-UUO (C); B, Top DEGs in the EC cell populations; D, Fraction of EC populations at different time points after R-UUO; E/F, Gene set enrichment analysis show upregulated GO terms in the top 20 injured DEGs from papillary AVR and DVR clusters 1 and 6 (E), and capillary angiogenic clusters 3 and 7 (F). G, CD31 antibody staining of ECs in the IM 84 days after R0UUO, and in Day 84 Nx controls. Scale bars=200um. H, Quantification of CD31 staining in proximal and distal IM after R-UUO. Data represented as mean+/- SEM with individual data points show. 1-way ANOVA, Nx vs. day 28 and 84 after R-UUO. If p<0.05, q values are shown for between group comparisons corrected for repeat testing.

### Temporal and spatially restricted expansion of myofibroblasts in the renal medulla

Re-clustering of medullary fibroblasts identified 8 subclusters: including 4 “fibroblast clusters 2, 6, 3, and 1”, “pericytes”, “Fib/MyoF cluster 4”, and “myofibroblast cluster 0” (Fig. 5A/B, see S. Table 2 for DEGs in these clusters). There was a marked increase in “myofibroblast cluster 0”, with a lesser increase in “Fib/MyoF cluster 4” 28 and 84 days after reversal of UUO (Fig. 5C/D). This was associated with a decrease in “fibroblast clusters 2 and 6”, a lesser reduction in “fibroblast cluster 1”, but no change in “Fib/interferon cluster 7”, “fibroblast cluster 3”, or “pericytes” (Fig. 5C/D). Immunostaining with an α-smooth muscle actin antibody (α-SMA, the gene product of *Acta2*) to identify myofibroblasts showed a marked increase in α- SMA^+^ cells the cortex and outer medulla at early time points. In contrast, expansion of α-SMA^+^ cells in the inner medulla only peaked at 28 days (Fig. 5E/F). By day 84 ISOM and inner medullary α-SMA staining was markedly reduced. These findings indicate that there is temporal and spatially restricted expansion of α-SMA^+^ myofibroblasts after reversal of UUO. At a transcriptional level, low levels of *Acta2* mRNA are expressed by “myofibroblast cluster 0” cells, which are characterized by high expression of *Fibronectin 1 (Fn1),* and collagens *Col1a2, Col1a1*, and *Col3a1* (Fig. 5B, S. Fig. 12A).

**Figure 5.**
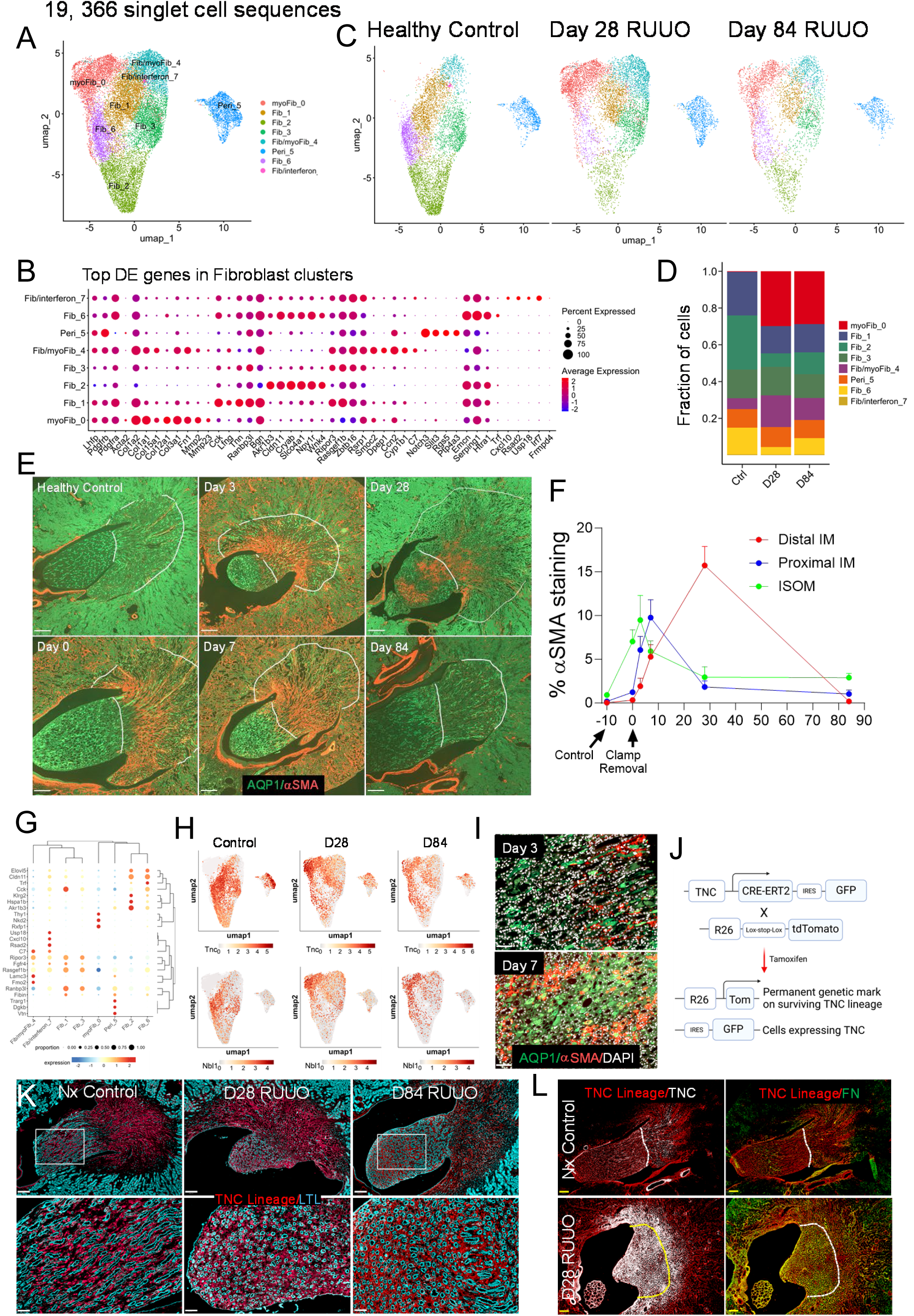
Temporal and spatially restricted expansion of myofibroblasts are derived from renal medullary interstitial cells. A, Re-clustering of fibroblasts identified 8 clusters, including 4 fibroblast, a pericyte, and two myofibroblast clusters in the combined data (A), and at different time points after R-UUO (C); B, Top DEGs in fibroblast clusters; D, Fraction of fibroblast populations after R-UUO; E, α-SMA and AQP1 staining of myofibroblasts, PTECs and DTLs in healthy controls (HCs), and at different time points after R- UUO. White lines demarcate the ISOM and IM boundaries. Scale bars=250um. F, Quantification of α- SMA staining in the proximal, distal IM, and ISOM after R-UUO. 4 HCs, 9 day 0, 8 day 3, 12 day 7, 4 day 28, and 14 day 84 mice. Means +/- SEM. G, Clustering of the top 5 DEGs in fibroblast clusters; H, UMAPs showing distribution RMIC markers, TNC and Dan/Nbl1; I, α-SMA staining in the distal IM day 3 and 7 after R-UUO. Scale bars=50um; J, TNC CreERT2 IRES GFP genetics used to evaluate RMIC lineages; K/L, TNC lineage (tdTomato) and LTL staining to label PTECs and IM CDs (K), white scale bars=200um, yellow scale bars=50um, GFP to detect cells currently expressing TNC and Fibronectin (FN) (L), scale bars=250um, dotted lines indicate the IM/ISOM boundaries.

### Myofibroblasts are largely derived from renal medullary interstitial cells after reversal of UUO

Unlike LOH and EC populations, the anchor genes used to identify fibroblast clusters did not provide information about the spatial localization of these cells within the renal medulla. To investigate this, we performed unbiased clustering of the top five DEGs in the fibroblast clusters (Fig. 5G). These data show that “myofibroblast cluster 0” is more closely related to “fibroblast clusters 2, 6”, and “pericytes”, while “Fib-MyoF and Fib-interferon clusters 4 and 7” are more closely related to “fibroblast clusters 1 and 3”. *Smoc2*, which is expressed by interstitial fibroblasts in the outer medulla, ^58^ is expressed by “Fib-MyoF cluster 4” in controls and extends to “fibroblast clusters 1 and 3”, and “myofibroblast cluster 0”, after reversal of UUO (S. Fig. 12A). These findings are consistent with data showing that Smoc^+^ myofibroblasts are increased in the outer medulla of mice after severe IRI-AKI,^59^ and suggests that “fibroblast clusters 1, 3” and “Fib-MyoF cluster 4” cells are likely to be localized in the outer medulla and give rise to outer medullary myofibroblasts. In contrast, *CryAB, Akr1b3,* and *Lmo7*, are expressed in control “fibroblast 2 and 6 clusters” and are restricted to “fibroblast cluster 2” after reversal of UUO (S. Fig. 12A). *Akr1b3* regulates osmolyte synthesis, and *Lmo7* is upregulated in hyperosmolar states, and both are increased in the inner medulla, ^60,61^ while *CryAB* is a molecular chaperone that is increased in all inner medullary cell types (S. Fig. 12B).^45^ This suggests that “fibroblast clusters 2 and 6” are likely to be restricted to the inner medulla. Since pericytes give rise to most of the cortical myofibroblasts expansion that occurs after IRI-AKI and irreversible UUO, ^62,63^ and since cortical expansion of myofibroblasts is associated with a reduction in pericytes,^64^ our observation that there was no change in the proportion of pericytes despite the increase in myofibroblasts after reversible UUO, suggests that pericytes make only a relatively small contribution the expansion of myofibroblasts in the renal medulla. However, “fibroblast clusters 2 and 6”, and to a lesser extent, “fibroblast cluster 1”, decrease after reversible UUO, suggesting that these cell populations contribute to renal medullary myofibroblasts expansion after reversal of UUO.

Renal medullary interstitial cells (RMICs) are specialized fibroblasts that are arrayed in columns along the length inner medullary CDs.^47,48^ *Tenascin C (Tnc)* and *Dan/Nbl1* are selectively expressed by RMICs,^65^ and are expressed in “fibroblast clusters 1, 2, 3, and 6” in control kidneys (Fig. 5H). In addition, α-SMA staining at early time points after reversible UUO identified localized areas of staining showing the characteristic ladder-like appearance of RMICs in the papilla in some areas, while in other areas there was a distinct pattern of α-SMA staining around areas of papillary necrosis (Fig. 5I).

These findings suggest that at least some of the inner medullary myofibroblasts maybe derived from RMICs, and that “fibroblast clusters 1, 2, 3, and 6” include RMICs. To test this hypothesis, we used a tamoxifen inducible, *Tenascin C Cre- ERT2-IRES-GFP* knock-in mouse ^65^ to evaluate expression of *Tnc* using the *Gfp* reporter, and the fate of RMICs by crossing *Tnc Cre-ERT2* mice with *R26R LSL tdTomato* reporter mice and treating with tamoxifen to activate CreERT2 (Fig. 5J). These mice selectively target RMICs, but not other fibroblast or pericyte populations in the kidney.^49,50,65,66^ After optimizing ureteric clamp times to allow for greater than 50% survival after Nx (6 days), these mice showed no reduction in tGFR, but a marked reduction in urinary concentrating capacity 28 days after reversible UUO (S. Fig. 13). Mice were treated with tamoxifen 6 weeks before undergoing reversible UUO to induce tdTomato expression. Co-staining with LTL and tdTomato antibodies showed that RMIC lineage cells extended throughout the papilla and into the ISOM just below the OSOM/ISOM junction (Fig. 5K). *Tnc* lineage cells had a similar distribution 28 days after reversal but had a more disorganized appearance in the inner medulla, where the typical laddering of RMICs between tubules was no longer apparent. By Day 84, there was only patchy re-organization of RMIC lineage cells in the inner medulla. Co-staining with GFP antibodies showed a more limited distribution of Tnc expression within the RMIC lineage domain both in controls and 28 days after reversal of UUO. Fibronectin (FN) staining, which is limited to a subset of RMIC lineage cells in control mice, was expressed by the majority of RMIC lineage cells in the inner medulla and ISOM 28 days after reversal of UUO (Fig. 5L, S. Fig. 14). α-SMA staining showed a similar distribution to FN, and like FN, a few inner medullary α-SMA^+^ cells were *Tnc* lineage negative. FN^+^ and α-SMA^+^ myofibroblasts in the renal cortex were also *Tnc* lineage negative. These findings indicate that: 1) most myofibroblasts in the ISOM and inner medulla are derived from RMICs; 2) there are at least two populations of RMICs, both of which express RMIC markers and contribute to the *Tnc* lineage: “fibroblast clusters 1 and 3” which also express *Smoc2* and are most likely localized in the ISOM, and “fibroblast clusters 2 and 6”, which express *CryAB, Akr1B,* and *Lmo7*, and are most likely localized in the inner medulla; and 3) the organization of RMICs in the papilla is disrupted as the cells differentiate into myofibroblasts and remains abnormal 84 days after reversal of UUO.

### Reduced numbers of enlarged collecting ducts in the papilla

Re-clustering of CD principal cells in the renal medulla identified two CD populations that are spatially restricted to the papilla, “IM CD cluster 2” and “deep IM CD cluster 0” (Fig. 6A/B, see S. Table 2 for DEGs in the CD clusters). “Injured CD cluster 5” was only present in reversible UUO samples, and there were two outer medullary “OM CD clusters 1 and 4”, and a population of papillary “urothelial cells in cluster 3” that are contiguous with “deep IM CD cluster 0” (Fig. 6A/B). Focusing on the two inner medullary CD clusters, there was a reduction in “CD clusters 0 and 2” that persisted 84 days after reversal and was associated with an increase in “urothelial cluster 3” cells (Fig. 6C/D). These findings were unexpected as we had shown that Aqp2 expression was restored, and LTL staining increased in the distal inner medulla 84 days after reversal of UUO (see Fig. 2M-O). To explore this further, we performed lineage analysis using *HoxB7 Cre* knock-in mice to genetically label all CD cells, but not nephronic epithelium including inner medullary LOH, by crossing these mice with *R26R LSL tdTomato* mice (Fig. 6E/F).^67^ A ureteric clamp time of 6 days gave a similar survival and reduction in tGFR and urinary concentrating capacity 84 days to that seen in mice used for scRNA-Seq studies after reversible UUO (S. Fig 15, see Fig 1). Consistent with our data using LTL staining to quantify the surface area of inner medullary CD cells, quantification of the *HoxB7* lineage nuclei showed that the proportion of CD lineage cells occupying the distal but not the proximal inner medulla was increased 84 days after reversal of UUO (Fig. 6G). However, this was also associated with a reduction in the total number of cells in the distal inner medulla, potentially because of the observed reduction in non-CD LOH and endothelial cell populations (Fig. 6G). On this basis, the relative increase in CD nuclei in the distal inner medulla could be because CDs occupy a larger space than the other inner medullary cell populations. More detailed analysis showed that this was linked to a reduction in the number of CD tubules throughout the inner medulla (Fig. 6H), but there was also an increase in CD tubule and luminal surface areas (Fig. 6I/J).

**Figure 6.**
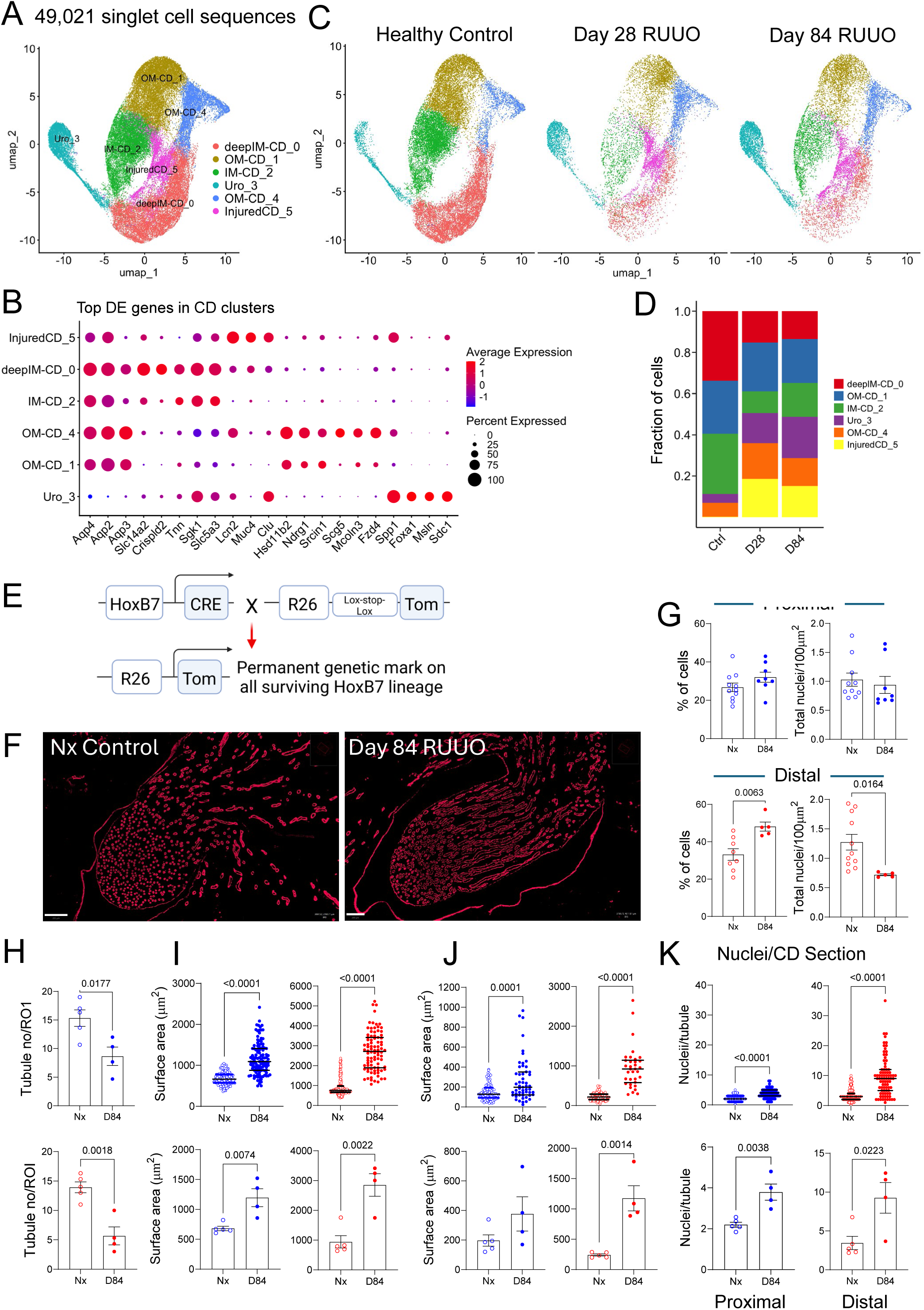
Reduced numbers of enlarged collecting ducts in the papilla after reversible UUO. A/C, Re-clustering of CD principal cells identified 6 cell populations, including deep IM and IM CD clusters 0 and 2, injured CD cluster 5, and papillary urothelium cluster 3 in the combined data (A), and at different time points after R-UUO (C); B, Top DEGs in the CD cell populations; D, Fraction of CD cells at different time points after R-UUO; E, Genetics showing HoxB7-dependent labeling with tdTomato of all renal medullary CD cells; F, HoxB7 lineage in controls and 84 days after R-UUO, scale bars=200um; G, Quantification of HoxB7 lineage cells and total nuclei in proximal and distal IM; H-K, Quantification of CD numbers (H), surface areas (I), CD luminal areas (J), and nuclei per CD section (K) using HoxB7 Cre tdTomato and AQP2 staining to delineate CD tubular cells. Individual data points, means +/- SEM. T-test, p values shown.

Consistent with our data showing that there was an increase in the percentage of *HoxB7* lineage cells in the distal inner medulla, this was associated with a corresponding increase in the numbers of nuclei detected per CD section (Fig. 6K). These data indicate that after an early period of active repair, there is a permanent reduction in the number of inner medullary CDs associated with compensatory increase in the size of the surviving CDs.

### Widespread and persistent expression of injury markers in collecting ducts throughout the inner medulla

“Injured CD cluster 5” cells express lower levels of CD markers found in other CD clusters and have increased expression of injury markers including *Lcn2, Spp1, Clu, Mmp7*, and *Ccn1* (Fig. 6B, S. Fig. 16A/B). Like “LOH injury cluster 4” cells, this suggests these cells are relatively dedifferentiated with evidence of persistent injury. Expression of CD markers, *Aqp2, 3*, and *4*, which are expressed in all of the CD clusters, and *Slc14a2*, which is dominantly expressed in “deep IM CD cluster 0”, are also increased in the “injured CD cluster 5” after reversal of UUO. This suggests that “injured CD cluster 5” cells are derived from both inner and outer medullary CD cell populations (S. Fig. 16A). To further characterize this population cells, we evaluated expression of the top 20 upregulated DEGs in “injured CD cluster 5” after reversal of UUO (Fig. 7A). A few of these DEGs, such as *Pdgf* and *Dcdc2a* are only expressed in “injured cluster 5” cells, while others, such as *Clu* and *Mmp7*, are also expressed in “urothelium”, or other more differentiated CD clusters, including *Lcn2, Spp1, Ccn2, Fcgbp, Grem2, Ly6e,* and *Gdf15*, after reversal of UUO (Fig. 7A/B, S. Fig 16B). We performed immunostaining to evaluate the spatial distribution of two of these injury markers, Spp1, and N-Gal, the gene product of *Lcn2*, in *HoxB7* Cre lineage mice. Spp1 was detected in TALs (which also express high levels of NaKATPase), and interspersed HoxB7 lineage cells (mostly likely intercalated cells) in the ISOM and proximal inner medulla in uninjured mice (Fig. 7C/D). However, Spp1 was also increased in papillary urothelium, TALs, and in the distal inner medullary *HoxB7* lineage CDs 84 days after reversal of UUO (Fig. 7D). There was also increased Spp1 staining of intercalated *HoxB7* lineage cells in the ISOM and proximal inner medulla after reversal of UUO than in control mice. N-Gal staining was absent in control mice and was markedly increased after reversal of UUO in papillary urothelium, AQP1^+^ DTLs, and *HoxB7* lineage CD cells in the inner medulla (Fig. 6E). These findings provide orthogonal validation of our scRNA-Seq data and indicate that there is a permanent and widespread spatial expression of injury markers, particularly in the distal inner medulla, after reversal of prolonged UUO.

**Figure 7.**
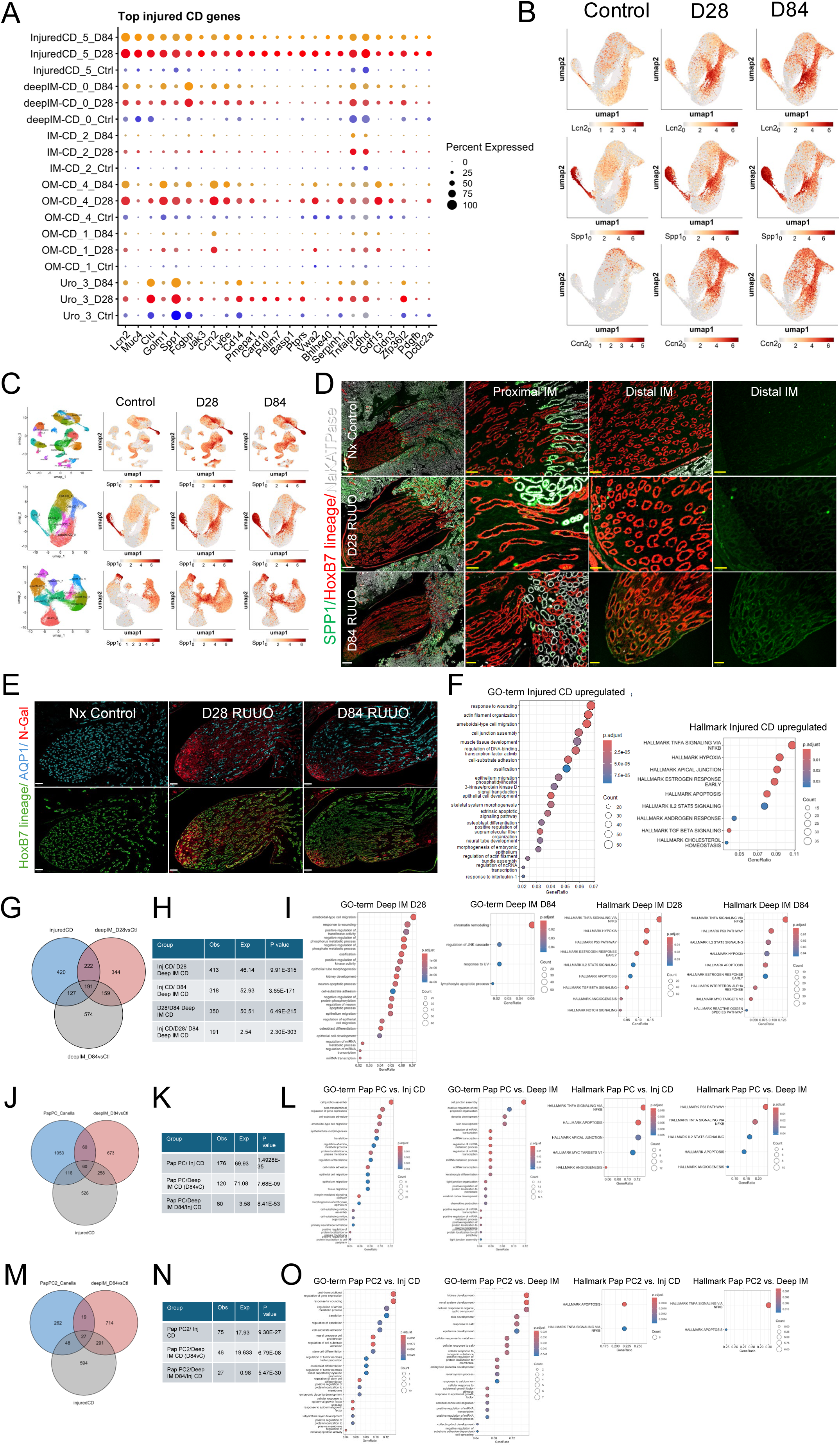
Injured inner medullary collecting ducts are hypoxic and share common pathways of migration, apoptosis, and inflammation that are activated in renal papillas of patients with renal stone disease. A, Clustering of the top 25 DEGs in injured CD cluster 5 cells in other LOH cell populations in controls, day 28 and 84 after R-UUO; B/C, UMAPs showing changes in the distribution of injury markers Lcn2, Spp1, and Ccn2 in CD clusters (B), and Spp1 in all of the renal medullary single cell clusters (C); D, Spp1 staining of the renal medulla of Nx controls, day 28 and 84 after R-UUO, along with NaKATPase (highest expression in TAL) and the HoxB7 lineage CDs. White scale bars=250um, yellow scale bars=50um; E, N-Gal/Lcn2 staining of the renal medulla of Nx controls, day 28 and 84 after R-UUO, along with AQP1 and the HoxB7 lineage CDs. Scale bars=100um; F, GSEA showing upregulated GO and Hallmark terms in DEGs from injured CD cluster 5 cells; G/H, Venn diagram illustrating the overlap between DEGs from injured CD cluster 5 cells, and Deep IM CDs ctrl vs. day 28, and ctrl. vs. day 84 R-UUO (G), and statistical significance between multi-set intersections from the Venn Diagrams (H); I, GSEA showing upregulated GO and Hallmark terms in DEGs from Deep IM CDs after R-UUO; J/K/M/N, Venn diagrams illustrating the overlap between DEGs from PapPC1 (J) and PapPC2 (M) populations from patients with renal stone disease, with injured CD cluster 5, and Deep IM CDs after R-UUO, and statistical significance between multi-set intersections from the respective Venn Diagrams (K/N); L,O, GSEA showing upregulated GO and Hallmark terms in DEGs in PapPC1 (L) and PapPC1 from renal stone disease patients (O) vs. injured CD cluster 5 cells after R-UUO.

### Injured inner medullary collecting ducts are hypoxic and have features of failed repair proximal tubular cells

Like injured LOH cells, GSEA of DEGs in injured CD cells showed enrichment for processes involved in cell migration, wound healing, apoptosis, and TNFα via Nfκb signaling (Fig. 7F, S. Table 2 for DEGs in the CD clusters, S. Table 3). This suggests overlap between injured CD and LOH clusters after reversal of UUO, and the previously identified FR-PTECs after IRI-AKI and irreversible UUO (S. Fig. 10).^38^ However, unlike “injured LOH cluster 4” and FR-PTECs, “injured CD cluster 5” cells also have a gene signature for hypoxic responses (Fig 7F, S. Fig. 10C/D, S. Table 3). This includes upregulation in *Ldha, Slc2a1/Glut1, Pfkl,* and *Pfkp*, which enhance anerobic glycolysis, and may represent a compensatory response to increasing hypoxia.

Since many of the DEGs in injured CD cells are also upregulated in “deep IM” and “IM CD” clusters 0 and 2 after reversal of UUO (Fig. 7A/B), we asked whether more differentiated CDs showed similar responses. There was substantial overlap in DEGs between day 28 and 84 “deep IM CD cluster 0”, “injured CD cluster 5”, and FR-PTECs after irreversible UUO (Fig. 7G/H, S. Fig. 17A-C). Cell migration, wound healing, apoptosis, inflammation, and hypoxic responses are also increased in “deep IM CD cluster 0” cells after reversal of UUO (Fig. 7I, S. Table 5). Hypoxia and inflammatory signatures were preserved in the “IM CD cluster 2”, but there was less overlap with injured CDs 84 days after reversal of UUO (S. Fig. 17D/E, S. Table 5). These findings contrast with inner medullary LOH clusters. For example, “lower IM TL cluster 5” did not show wound healing, cell migration or hypoxic responses, but did show evidence of activated TNFα signaling 28 days after reversal of UUO (S. Fig. 17F/G, S. Table 6). “IM ATL cluster 0” cells also showed evidence of wound healing responses at day 28 and 84 and activated TNFα signaling at day 28 but not 84 days after reversal of UUO, but there was also no evidence of a hypoxic response (S. Fig. 17H/I, S. Table 7). These findings indicate that there is widespread and persistent CD injury in the inner medulla after reversal of UUO, but that unlike FR-PTECs and injured LOH cells, CD injury is characterized by a hypoxic response that probably reflects an adaptive response to increased papillary hypoxia.

### Long-term effects on cellular interactions in the renal papilla

To determine how changes in CD, LOH, and fibroblasts affect cellular interactions after reversal of UUO, we evaluated ligand-receptor pairing between cell clusters identified from our scRNA-Seq dataset using CellChat (S. Fig. 18, see S.Table 8 for complete information on ligands and receptor pairs in controls and after reversal of UUO).^68^ Focusing on ligand-receptor pairing in the inner medulla, the most striking finding is the *de novo* activation of the Spp1 pathway in injured “LOH cluster 4” and “injured CD cluster 5” cells. Spp1, signals via integrins and activates proinflammatory and profibrotic responses after AKI.^69,70^. Other pathways, including canonical Wnt signaling, show autocrine activity in control “deep IM CDs” which express *Wnt7b* and *9b*, but the same ligands are secreted by injured LOH and CD clusters after reversal of UUO. Likewise, the *PdgfA* ligand is expressed by “IM ATL cluster 0 cells”, and the Pdgf receptor*, PdgfRA*, is expressed by inner medullary “fibroblast clusters 2 and 6” in controls, but at 28 days, *PdgfA* ligand is expressed by “injured LOH cluster 4”, and its cognate receptors, *PdgfRA* and *PdgfRB*, are dominantly expressed by “myofibroblast cluster 0” cells. By day 84, dominant *PdgfA* ligand expression is restored in “IM ATL” cells, and while “myofibroblasts cluster 0” cells remain the dominant responding cells expressing Pdgf receptors. Conversely, the non-cannonical Wnt ligand, *Wnt5A*, which is normally secreted by “fibroblast 1 and 2 clusters” with “deep IM CD” responder cells, switches to “myofibroblast cluster 0” and “injured LOH cluster 4” cells after reversal of UUO. Activation of these pathways could play a role in maintaining the inflammatory milieu of the inner medulla, and in promoting remodeling in the papilla that occurs between 28 and 84 days after reversal of UUO. However, in addition, long-term alterations in these cellular interactions could play a role in disrupting normal tissue homeostasis and perpetuating the cellular disorganization that occurs in the renal papilla after reversal of UUO.

### Common pathways that are activated in renal papillas of patients with renal stone disease

To determine whether similar responses are activated in patients with renal medullary disease, we evaluated a published scRNA-Seq analysis of renal papillary biopsies from patients undergoing ureteroscopy for recurrent renal stone disease.^71^ The authors compared these biopsies with normal samples obtained from patients undergoing nephrectomy for other reasons, or deceased donor kidneys that were not suitable to transplantation. They identified two renal papillary principal cell populations (“PapPC1” and “PapPC2”), both with features of CD PCs, but with “PapPC2” showing features of chronic injury, including expression of *Lnc2, Spp1,* and *Mmp7*. Like Spp1 and N-Gal expression in the distal CD and LOH after reversal of UUO (see Fig. 7D/E), these injury markers showed a widespread increase in expression in renal stone disease patients. There was a significant overlap between DEGs from renal stone disease vs. normal human “PapPC1” and “PapPC2” cells, with “injured CD cluster 5”and “deep IM CD cluster 0” cells 84 days after reversal of UUO (Fig. 8J/K/M/N, see S. Table 3 for DEGs in “PapPC1” and “PapPC2” cells ^71^). For example, *Mmp7* is increased in PapPCs from patients with renal stone disease,^71^ and in injured CD and LOH clusters after reversible UUO. GSEA shows enrichment for pathways involved in wound healing, cell migration, apoptosis, and TNFα signaling in diseased “PapPC1” and “PapPC2” cells that are also increased in “injured CD cluster 5”, and “deep IM CD” cells after reversal of UUO (Fig. 7L/O, S. Tables 9/10). However, unlike reversible UUO, there was no activation of hypoxic pathways in the human renal stone disease patients. These data show that common pro-inflammatory, cell migration, and wound healing responses are activated in human PapPC cells from patients with renal stone disease patients, and in mouse inner medullary CD cells after reversal of UUO. However, the enhanced hypoxic response we saw in inner medullary CDs after reversible UUO, does not occur in patients with renal stone disease.

**Figure 8.**
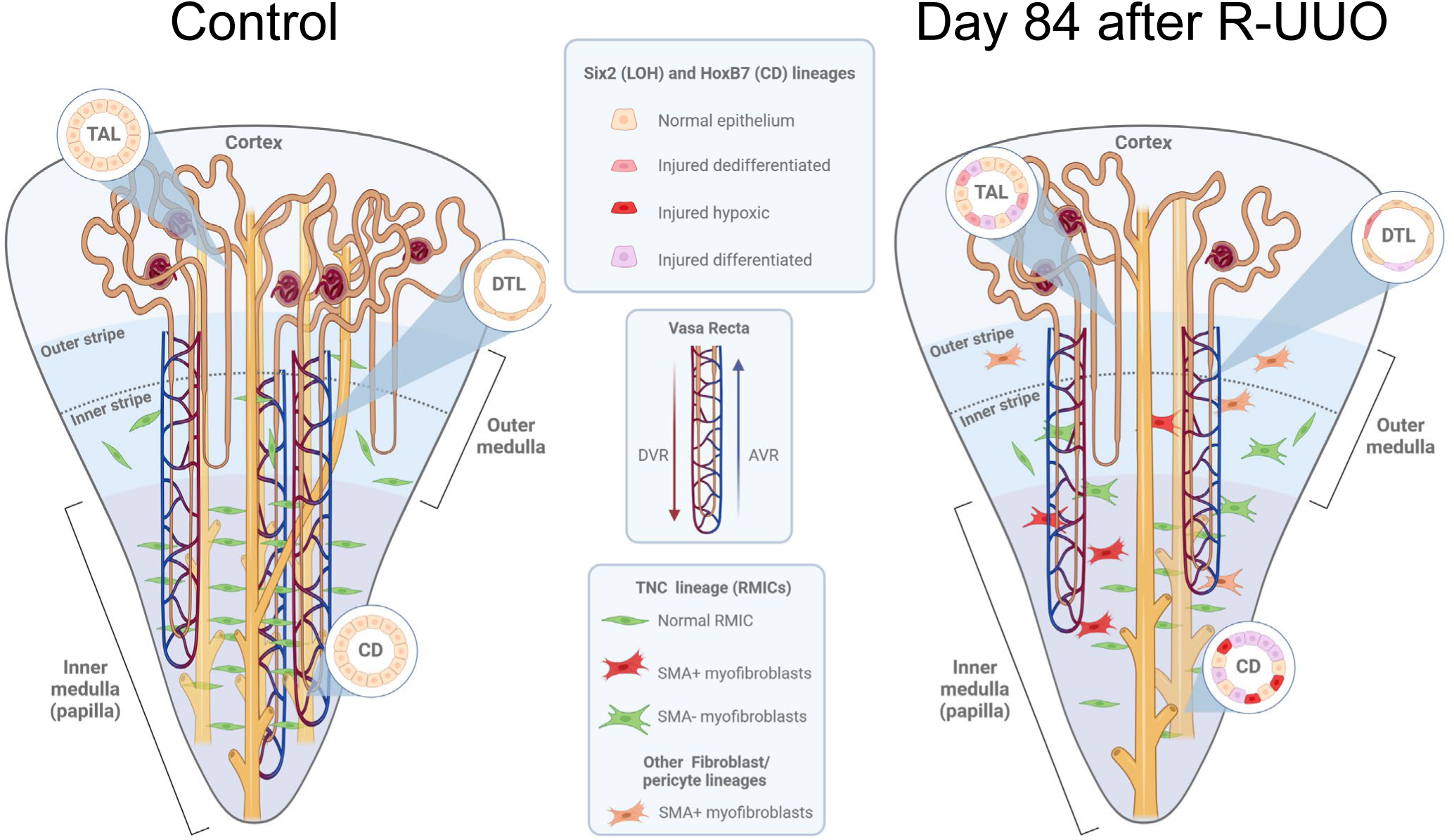
Permanent defects in the organization and cells in the renal papilla after reversal of ureteric obstruction. Summary of the experiments and data showing that there is a permanent reduction in number of inner medullary loops of Henle (LOH) and vasa recta extending into the papilla, reduced numbers of enlarged collecting ducts (CDs), and depletion and mislocalization of renal medullary interstitial cells (RMICs) after reversal of UUO (R-UUO). scRNA-Seq was used to evaluated all renal medullary cell populations, Six2, HoxB7, and TNC Cre mice were used to label and evaluate LOH, CD, and RMIC lineages. DTL, descending thin limb, DVR, descending vasa recta, AVR, ascending vasa recta.

## Discussion

We describe a robust new model of reversible UUO in which we were able to evaluate renal function up to three months after reversal of prolonged UUO by removing the contralateral kidney. Our data show that despite reversal of the obstruction, there is a permanent reduction in GFR associated with a disproportionately large reduction in urinary concentrating capacity, indicating that there is a permanent defect in renal papillary function. These findings are consistent with clinical observations that in addition to the well-documented polyuria that occurs in the first few days after reversal of urinary obstruction, ^72,73^ some patients have a long-term defect in their ability to maximally concentrate their urine.^7^ These experimental findings are of clinical significance since: 1) the cellular composition and organization of the renal medulla is highly conserved between rodents and humans suggesting that the pathophysiological changes that occur in rodents after reversal of obstruction are likely to reflect similar changes that occur in humans; 2) the high penetrance of this phenotype in our experimental model suggests that this effect is likely to be more common than generally thought; and 3) the failure to concentrate urine predisposes susceptible patients to dehydration and recurrent AKI, and may contribute to documented progressive decline in GFR that has been documented in these individuals.^5,6^

Our studies show that while there is widespread cellular damage throughout the inner medulla along with shortening of the renal papilla at early time points after reversal of UUO, there is a robust regenerative response so that by 84 days mice have normal sized papillas that are indistinguishable from controls using standard histological staining. At least some of this regeneration appears to be driven by proliferation of Sox9^+^ and Ki67^+^ dedifferentiated tubular cells that repopulate denuded basement membranes. However, our scRNA-Seq and quantitative immunostaining analyses revealed that repair is incomplete, and there are permanent changes to all of the major cellular compartments in the inner medulla after reversal of UUO (summarized in Fig. 8). This includes a permanent reduction in long loops of Henle, vasa recta, and a reduction in the number of inner medullary CDs with a concomitant increase in their cross-sectional diameters. The enlargement of surviving or regenerated CDs is unique to this epithelial compartment. We speculate this may occur as a compensatory response to increasing urine flow through fewer CDs after reversal of UUO. In addition to changes in inner medullary epithelia and capillaries, there is a permanent reduction and mislocalization of renal medullary interstitial cells in the inner medulla, a specialized population of renal fibroblasts that regulate urinary concentrating capacity by regulating AQP2 levels and activity in CDs.^49,50^ Moreover, analysis of ligand-receptor pairing indicates that there is long-term disruption of some of the normal cellular interactions in the renal medulla, including Pdgf and non-cannonical Wnt signaling, which regulate both epithelial and fibroblast stability and function, and could also play a role in mediating tissue remodeling that occurs after reversal of UUO. On this basis, the breakdown in organization of the major cellular compartments in the inner medulla would be expected not only to disrupt the counter current multiplier system that is required to generate osmotic gradients in the renal medulla,^1,9^ but could also play a role in destabilizing cellular interactions and perpetuating the cellular disorganization that occurs in the papilla after reversal of UUO.

In addition to these changes in the organization and number of inner medullary epithelial cell populations, there is widespread activation of pathways involved in regulating cell migration and inflammation in injured papillary LOH and CD cells after R-UUO. These responses are similar to the responses that have been shown to be activated in maladaptive or failed repair PTECs, which are thought to drive local inflammation, fibrosis, and CKD progression after AKI.^36,37^ Consistent with these findings, analysis of ligand-receptor pairing in the papilla showed that there is a persistent, *de novo* activation of the Spp1 pathway in injured LOH and CD cells after reversal of UUO. Since Spp1 signals via integrins and has been shown to activate proinflammatory and profibrotic responses after AKI, ^69,70^ this response could play a role in promoting the persistent pro-inflammatory state of inner medullary epithelial cell populations. Moreover, the same cell migration and pro-inflammatory pathways that are activated in the renal papilla after reversible UUO, are activated in renal papillary CDs from patients with renal stone disease.^71^ These findings suggest that these pathways represent shared epithelial responses to different types of cortical and medullary injury in both humans and mice, and that findings using our mouse model may provide insights into the pathophysiology of papillary dysfunction in patients after reversal of urinary obstruction. As an example of this, *Mmp7*, which is expressed by injured CDs after reversal of UUO, is also upregulated in papillary CDs from patients with renal stone disease. ^71^ In addition, urinary MMP7 levels correlate with disease activity in patients with renal stone disease,^71^ and increased levels of urinary MMP7 is predictive of renal outcomes in children after reversal of uretero-pelvic junction obstruction.^74^ However, there are also notable differences in the cellular responses to different types of injury. For example, while we show that there is increased expression of genes involved in regulating compensatory responses to worsening hypoxia in inner medullary CDs after reversal of UUO, including a number of genes involved in promoting anaerobic glycolysis as an alternative energy source, this is not seen in failed repair PTECs after irreversible UUO, injured LOH cells after reversal of UUO, or in papillary CDs from patients with renal stone disease. This most likely results from a compensatory response to hypoxia resulting from the reduction in vasa recta capillaries supplying oxygenated blood to the inner medulla that occurs after reversal of prolonged UUO. This may explain why inner medullary CDs, but not long LOH cells which do not have this adaptive response to hypoxia, are able to survive in the hostile hypoxic environment of the papilla after reversal of UUO. Since this hypoxic response is not seen in CDs from renal stone disease patients, these data also suggest that capillary rarefaction may exert additional hypoxic stress on inner medullary CDs after reversal of prolonged urinary obstruction.

In conclusion, our data indicate that despite a robust regenerative response at early time points after reversal of UUO, there are permanent changes to all of the major cellular compartments of the renal papilla that account for the permanent defect in urinary concentrating capacity that occurs after reversal of prolonged ureteric obstruction. These findings have additional implications for the management of patients with urinary obstruction because: 1) the histological appearance of renal papillas are essentially normal 3 months after reversal of urinary obstruction in our model, standard histological assessment of renal papillas in these patients is unlikely to detect major differences in the cellular organization of the renal papilla that occurs after reversal of obstruction; and 2) while focused molecular studies in patients after reversal of urinary obstruction are therefore needed to establish common responses that can be leveraged to develop new therapeutic approaches to prevent long-term renal papillary dysfunction, some of the molecular changes seen in this mouse model are also seen in renal papillas obtained from patients with renal stone disease, another cause of renal papillary injury, suggesting that this model is likely to reflect similar changes occurring in humans.

## Methods

### Sex as a biological variable

Our studies only evaluated male mice. It is currently unknown whether these findings are relevant to female mice.

### Reversible unilateral ureteral obstruction (R-UUO)

R-UUO was performed in 12–13-wk male mice. For this, a small vascular clamp was applied to the left ureter, the kidney and clamped ureter gently pushed back into the retroperitoneal space. After a variable interval of time depending on the mouse strain (between 5 and 7 days), the left kidney exteriorized, and the vascular clamp carefully removed. For long-term studies, a contralateral nephrectomy (Nx) was performed 10 days after R-UUO. Nx controls were performed at the same age/time point as the R-UUO studies. After completion of the studies, the left kidney was initially checked for obstruction, and if uncertain, methylene blue injected into the renal pelvis to determine whether the left ureter was patent. Since external evidence of hydronephrosis resolves within ∼24 hrs of reversal of obstruction, if there was evidence of persistent obstruction, data was discarded (S. Methods 1 includes detailed information about tissue harvesting, surgical technique and renal function tests).

### Immunofluorescence staining

FFF and FFPE sections were prepared, washed, blocked, and primary and secondary antibodies applied, as described (see S. Table 1 for the list of antibodies, fluorophores, and conditions).^55^ Digital images were scanned using Zeiss AxioScan Z1 slide scanner (Carl Zeiss Microscopy GmbH, Oberkochen, Germany, 10X), and downloaded into QuPath for analysis (version 0.5.0). For quantification, regions of interest (ROIs, cortex, OSOM, ISOM, proximal 1/3 and distal 1/3 of the IM were demarcated using the QuPath annotation tool. If sections through the IM were less than 1mm in length, we would only quantify the area of interest in the proximal IM. To quantify collecting duct surface areas, digital images of HoxB7 lineage-labelled sections with IM lengths of ≥1mm were used. To quantify tubular surface areas, we evaluated circular-shaped CDs defined as having major/minor axis ratios of >0.8, as described.^75^ We quantified the surface area of each tubule and its lumen using the freehand selection tool in Fiji, 15 and 30 tubules in each area of interest, and the number of nuclei counted in each tubule cross section (S. Methods 1 includes detailed information on staining and image quantification, and S. Methods 2 provides a detailed immunofluorescence staining protocol).

### Single cell RNA sequencing (scRNA seq)

Papillae were dissected at two times points after injury (d28, d84 after R-UUO) and from healthy controls. Three to four papillae from the same time point were placed in a cryogenic vial and flash frozen with liquid nitrogen, then stored at -80C. Each frozen tube was fixed and dissociated into single cells using the 10x Genomics Chromium Next GEM Single Cell Fixed RNA Sample Preparation Kit #1000414. Dissociated cells from each of the pooled samples were individually hybridized with bar coded genome-wide probe sets from 10X Genomics Flex kits according to the manufacturer’s instructions. After probe hybridization and construction of libraries, samples were sequenced using an Illumina Novoseq 6000 PE150 sequencer (S. Methods 3 for a detailed sample preparation protocol).

### Bioinformatics pipeline and analysis of transcriptome data

After quality control measures were applied, dimensionality reduction was performed using Principal Component Analysis (PCA) and visualized with uniform manifold projection (UMAP). Cell population identities were identified using published anchor genes for renal medullary loop of Henle (LOH), collecting duct (CD), fibroblast, endothelial cell (EC), and immune cell subclusters. ^45,52,53^ Differential gene expression analysis between clusters and timepoints was carried out using the FindAllMarkers function and the FindMarkers function, respectively. GO and Hallmark pathway gene set enrichment analysis (GSEA) was performed and visualized by ClusterProlifer, as well as published FR-PTEC DEG gene sets from published data. ^38^ The subset function was used to extract clusters of each cell type on which PC analysis, re-clustering, and differential gene expression analysis were carried out.

The injured LOH cluster 4 was subclustered using the FindSubCluster function. Cell-cell communication analysis was performed using CellChat. ^68^ Data were visualized using Seurat, dittoSeq, and ShinyCell R packages (see S. Methods 1 for additional information about the quality control and analytical approaches used to evaluate the scRNA-Seq data).^76^

### Statistical analysis

were performed using GraphPad Prism 10.1.2 software (San Diego, CA), and results expressed as means +/- SEM, with individual datapoints shown where indicated. Two-tailed, unpaired T-Tests were used to compare two groups, 1-way ANOVA to compare between 3 or more groups, and 2-way ANOVA to compare groups over time. If p values were <0.05 for between group analyses, multiple between group comparisons were performed controlling for false discovery rates (FDR) of p<0.05 using the two-step method of Benjamini, Krieger and Yekutieli. Q values are indicated after correction for multiple between group comparisons. Statistically significance between multi-set intersections from the scRNA-Seq subcluster Venn Diagrams was performed using the SuperExactTest function in R.^77^

## Study approval

All mouse experiments were approved by the Vanderbilt Institutional Animal Care and Use Committee.

## Data availability

All relevant data are found in this article and in the supporting data file. Single cell RNA sequencing data was deposited with the NCBIs Gene Expression Omnibus repository.

## Author Contributions

T.V. performed the scRNA-Seq data analysis and prepared the scRNA-Seq figures and supplemental datasets with help from A.G. R.D. performed mouse genotyping, R-UUO surgery and renal function assays with help from M.B., K.A.C., and A.M. M.B., A.M., W.E.S., T.R., and V.S. performed immunostaining and quantitative images analyses, R.D. quantified Sirius red staining. H.Y. provided renal pathology advice on R-UUO samples. With help from A.J.D, M.D.C. conceptualized and planned studies, interpreted all of the data, finalized and prepared figures, and wrote the manuscript.

## Acknowledgements

The Vanderbilt Technologies for Advanced Genomics (VANTAGE) performed 10X Genomics bar coded probe hybridization, library preparation, and sequencing for the scRNA-Seq studies, the Vanderbilt Translational Pathology Shared Resource (TPSR) processed FFPE tissue blocks, and Dr. Jeff Spraggins from Vanderbilt University allowed us to use their Zeiss AxioScan Z1 slide scanner. Andy McMahon from USC Keck School of Medicine kindly provided HoxB7 Cre transgenic and Six2 eGFP-Cre BAC transgenic mice. Chuan-Ming Hao Fudan University gave permission for us to use Tenascin C Cre-ERT2-IRES-GFP mice provided by Agnes Fogo at VUMC. In addition, the University of Alabama at Birmingham-University of California-San Diego O’Brien Core Center for Acute Kidney Injury Research performed serum creatinine assays. This Research was supported by the National Institute of Diabetes and Digestive and Kidney Diseases (NIDDK) under Award Numbers: RO1DK128823 and U54 DK120058-01 (MDC), and UC2DK126122 (MDC and AJD).

---

**S. Figure 1.**
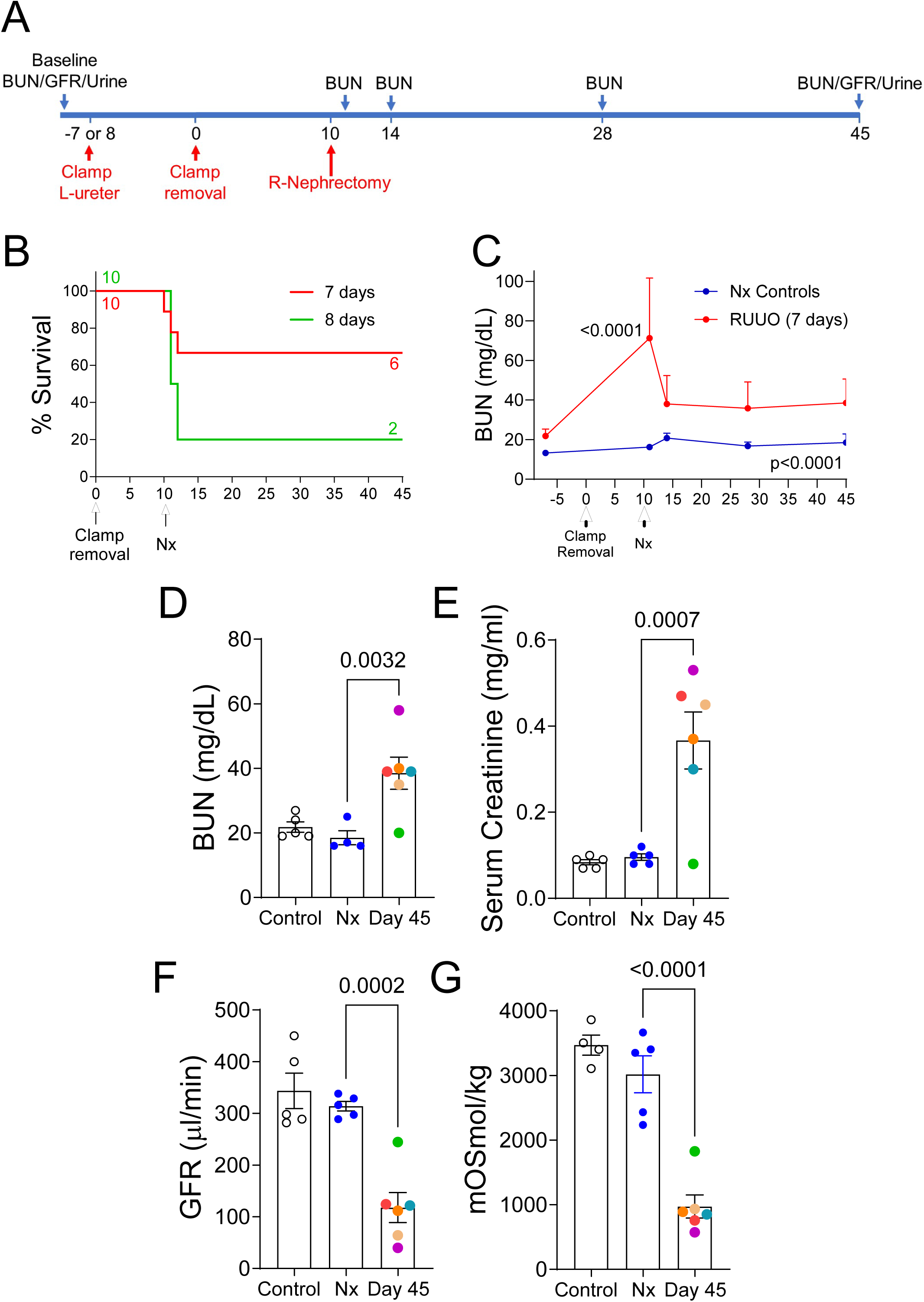
Optimizing ureteric clamp times in male BALB/c mice to achieve long-term survival after R-UUO. Male BALB/c mice underwent 7 or 8-day R-UUO and contralateral nephrectomy, or nephrectomy alone (Nx). A, Study design; B, Survival, numbers of mice indicated; C, BUN time course after 7-day R-UUO; D-G, BUN, serum creatinine, tGFR, and urine osmolality after 18hrs water restriction, in healthy controls (HC), Nx, and day 45 after 7-day R-UUO. C, BUN mean +/-SD. 2-way ANOVA. D-G, individual data points shown with mean +/-SEM. R-UUO datapoints are color coded to show the relationship between BUN, creatinine, tGFR, and urine osmolality in individual mice. 1-way ANOVA Nx vs. HC and R-UUO. If p<0.05, q values shown for between group comparisons corrected for repeat testing.

**S. Figure 2.**
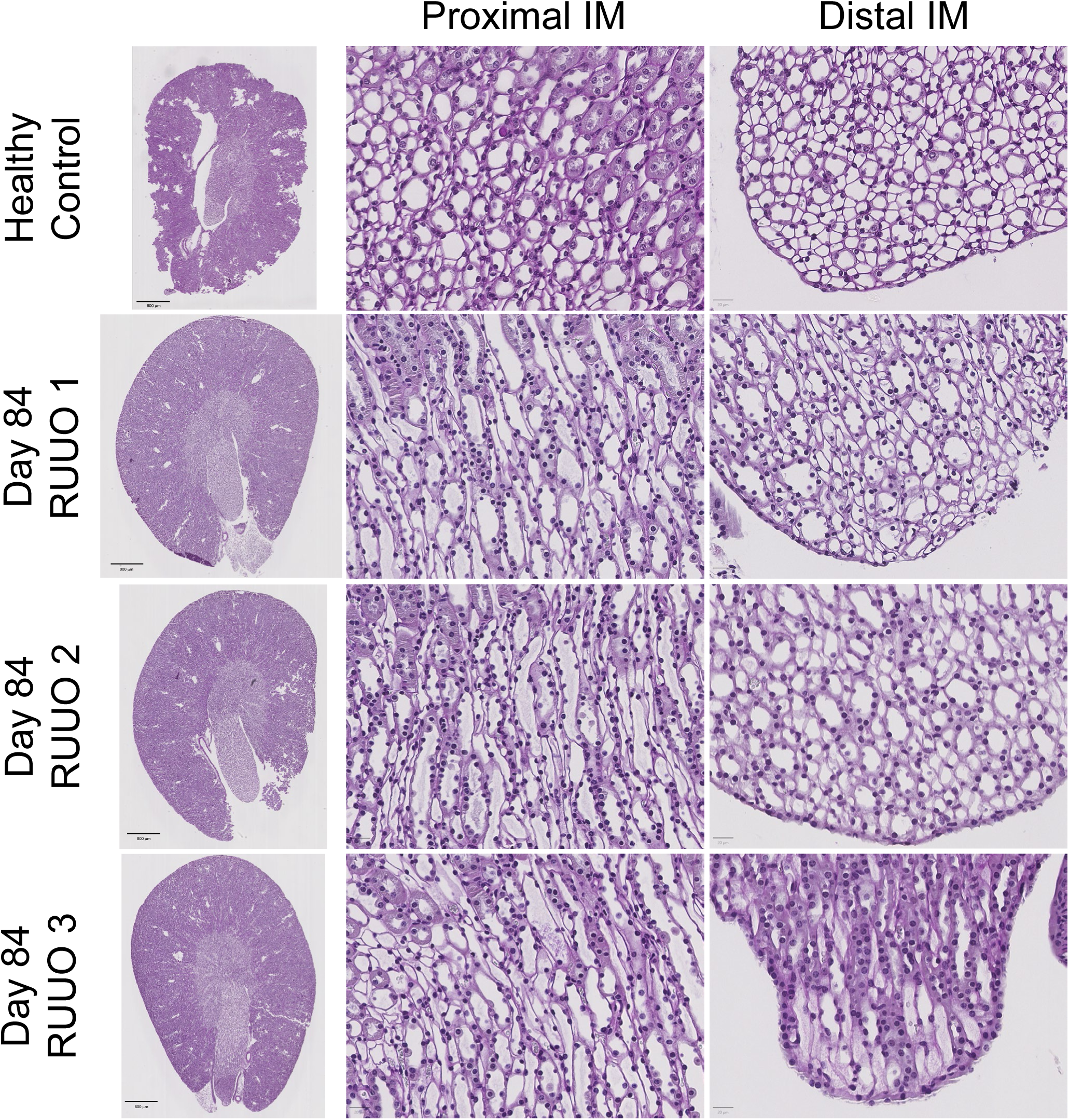
Normal histological appearance of renal papillas 84 days after R-UUO. PAS staining of a healthy control, and 3 Day 84 R-UUO kidneys scale bars=800um, proximal and distal inner medulla (IM), scale bars=20uM, as indicated.

**S. Figure 3.**
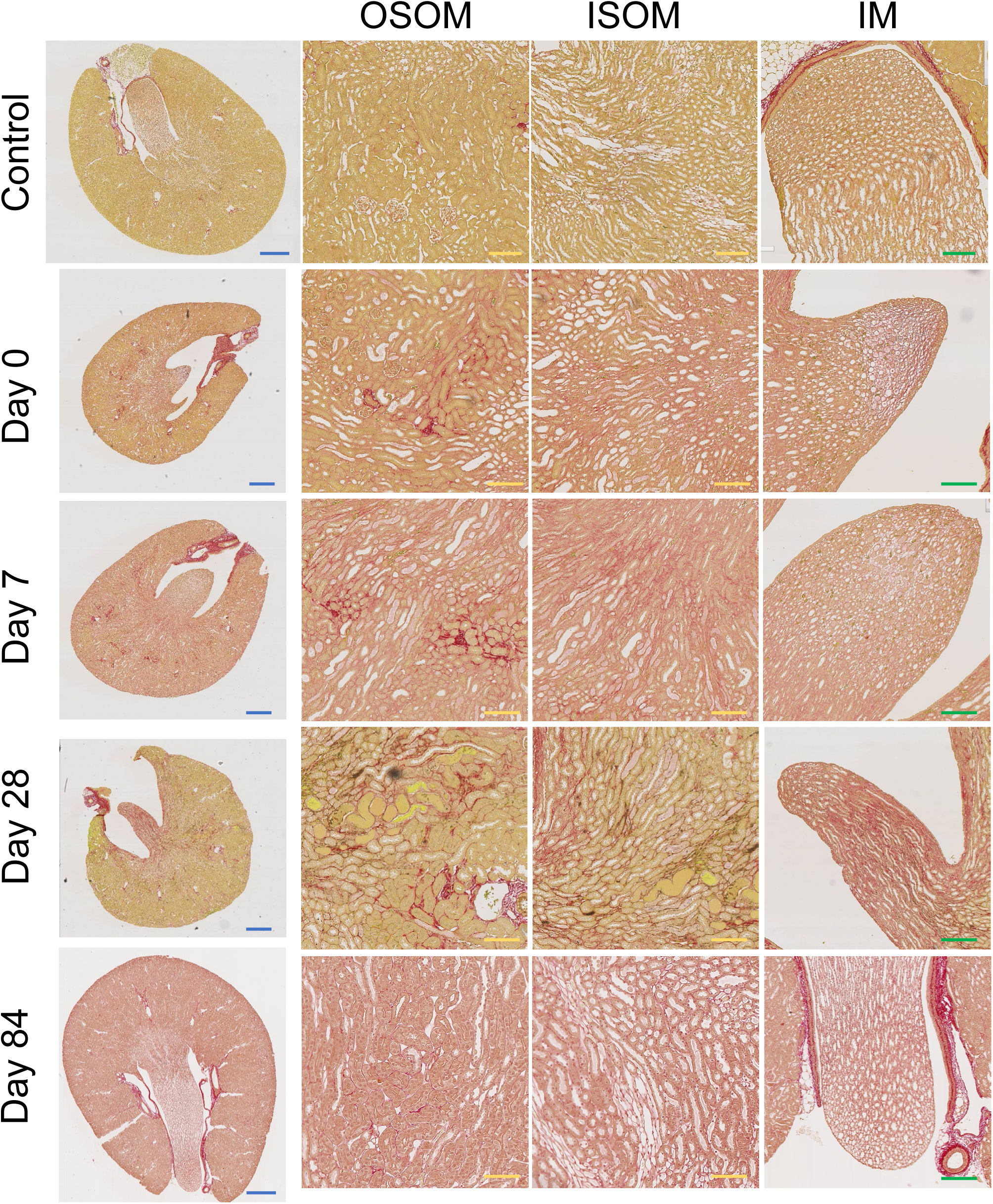
Time course of collagen 1 staining in the inner and outer medulla after R-UUO. Sirius red staining of type 1 collagen at different time points after R-UUO showing whole kidneys scale bars=800um (blue), ISOM and OSOM, scale bars=50um (yellow), inner medulla, scale bars=100um (green).

**S. Figure 4.**
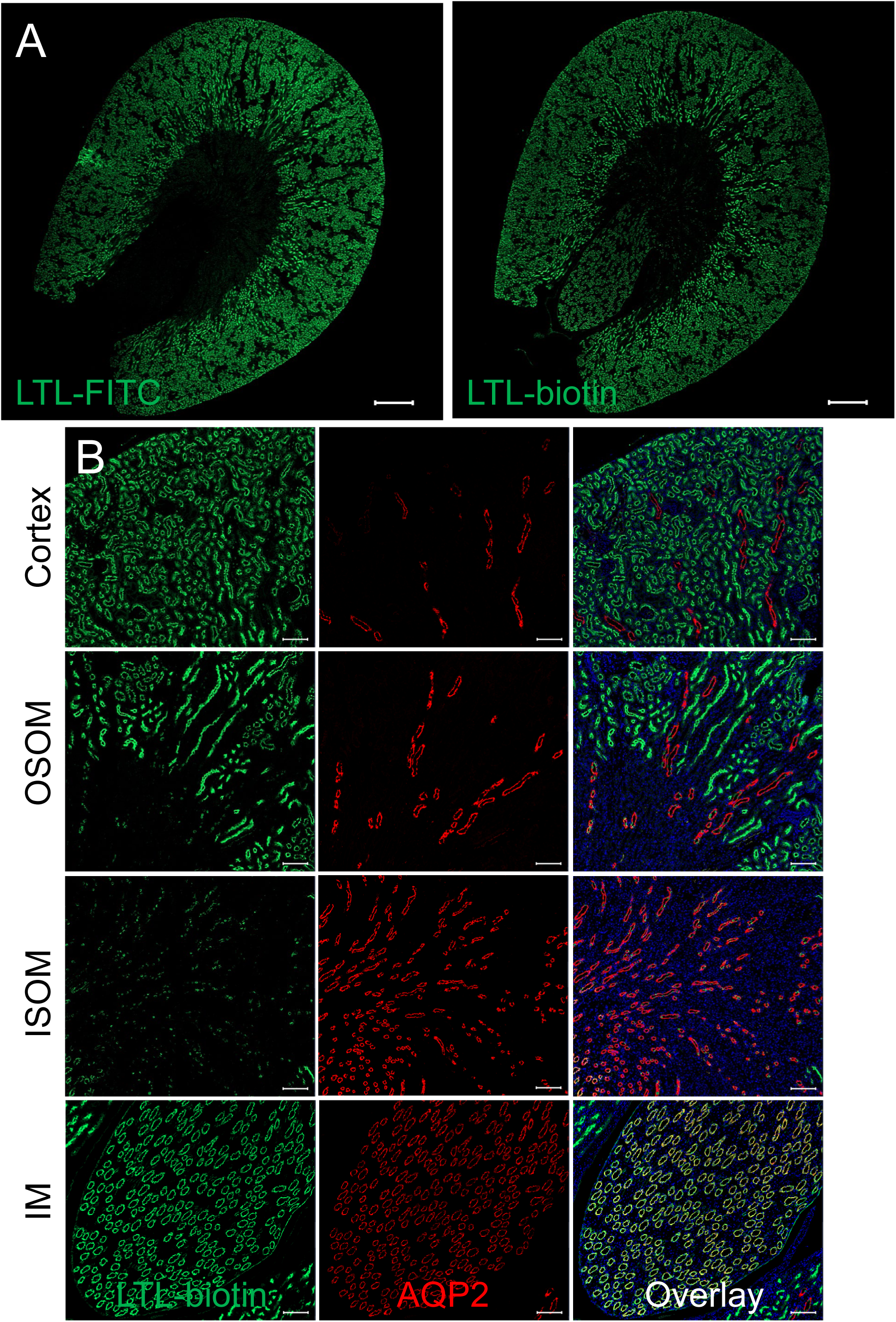
LTL stains inner medullary collecting ducts. A, Low power images comparing FITC-conjugated LTL and biotin conjugated LTL detected using neutravidin FITC amplification. In additional to widespread staining of proximal tubular epithelial cells throughout the cortex and OSOM, LTL-biotin amplification detects tubules in the inner medulla (IM). Scale bars=500uM ; B, Co-labeling LTL biotin with AQP2 showing co-localization of LTL staining with AQP2 in the IM. Additional punctate LTL staining of AQP2 negative collecting duct cells in the outer medulla and cortex. Scale bars=100uM.

**S. Figure 5.**
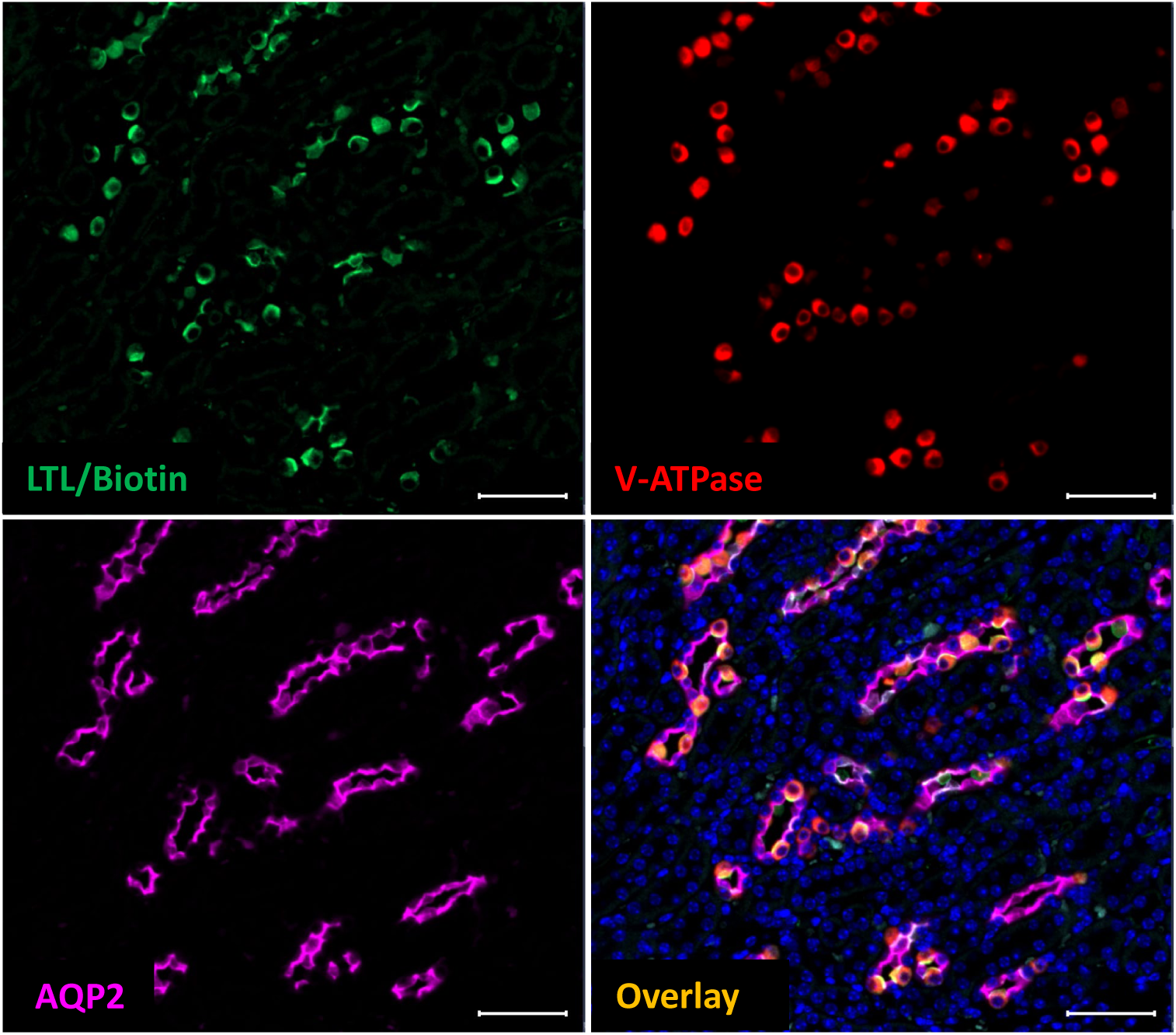
LTL stains intercalated cells in the inner stripe of the outer medulla. Co-labeling with LTL-biotin, AQP2 and V-ATPase B1/2 antibodies shows co-localization of LTL with V-ATPase staining in AQP2 negative V-ATPase positive intercalated cells in AQP2 positive collecting ducts seen in the ISOM. This is also seen in the outer stripe and cortical collecting duct intercalated cells. Scale bars=50uM.

**S. Figure 6.**
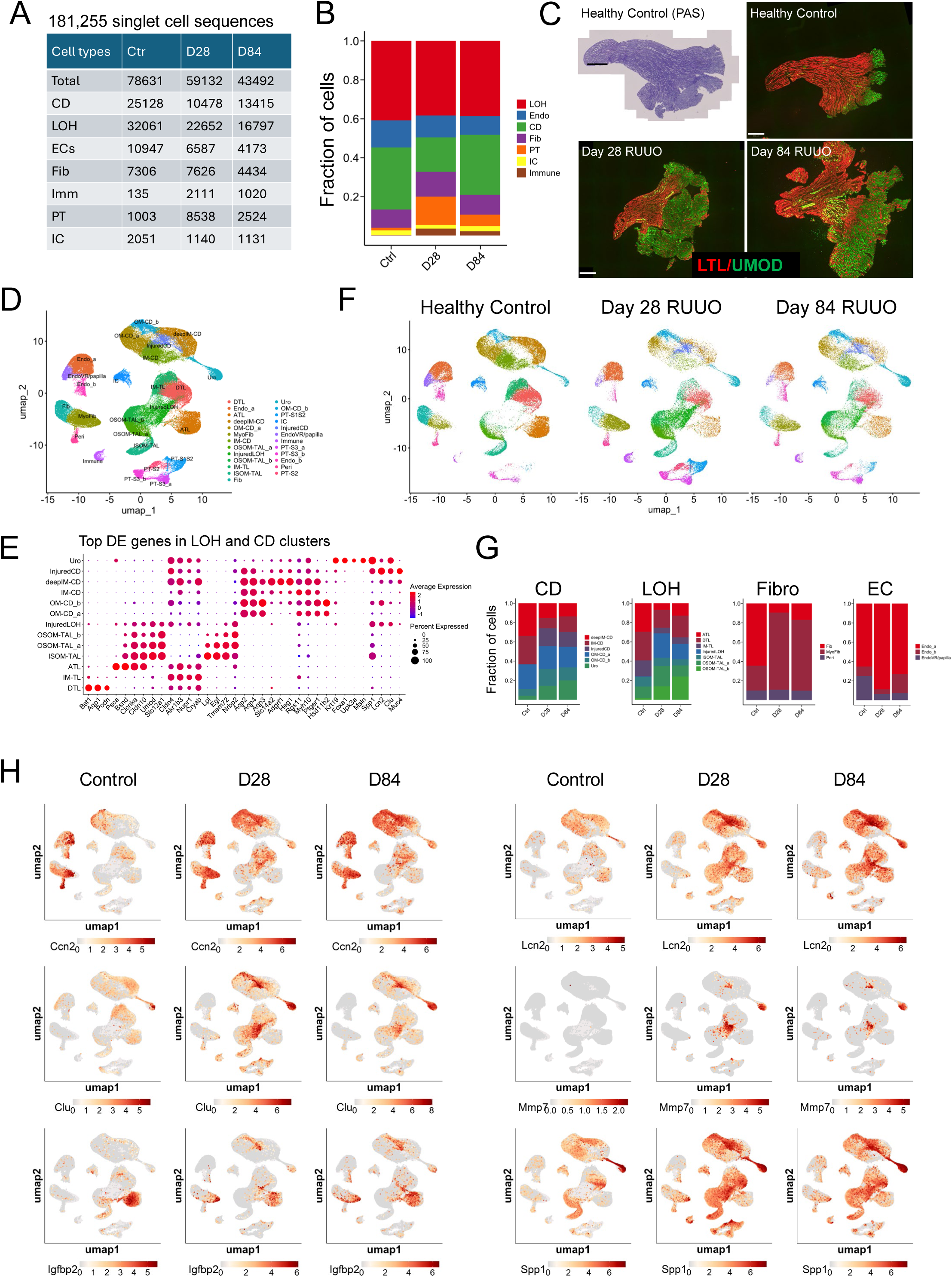
Single cell RNA sequencing atlas of mouse renal medullas after R-UUO. We performed single cell RNA sequencing (scRNA-Seq) on isolated renal medullas dissected kidneys from multiple healthy controls and mice 28 and 84 days after R-UUO. A, 181, 255 single cell sequences were obtained from 7 major renal medullary cell populations; B, Fraction of each cell type in the combined scRNA-Seq data set; C, Renal medullas dissected from healthy controls, and days 28 and 84 after R-UUO stained with PAS, or with LTL and uromodulin antibody. Scale bars=500um. D/F, “Big” UMAP and identification of 25 different cell populations using published anchor genes in the combined data (D), and at different time points after R-UUO (F); E, Top DEGs in LOH and CD sub-clusters; G, Fraction of CD, LOH, fibroblast and endothelial cell (EC) sub-clusters at different time points; H, “Big” UMAPs showing the expression distribution of selected injury markers in different renal medullary cell populations from controls, day 28 and 84 after R-UUO .

**S. Figure 7.**
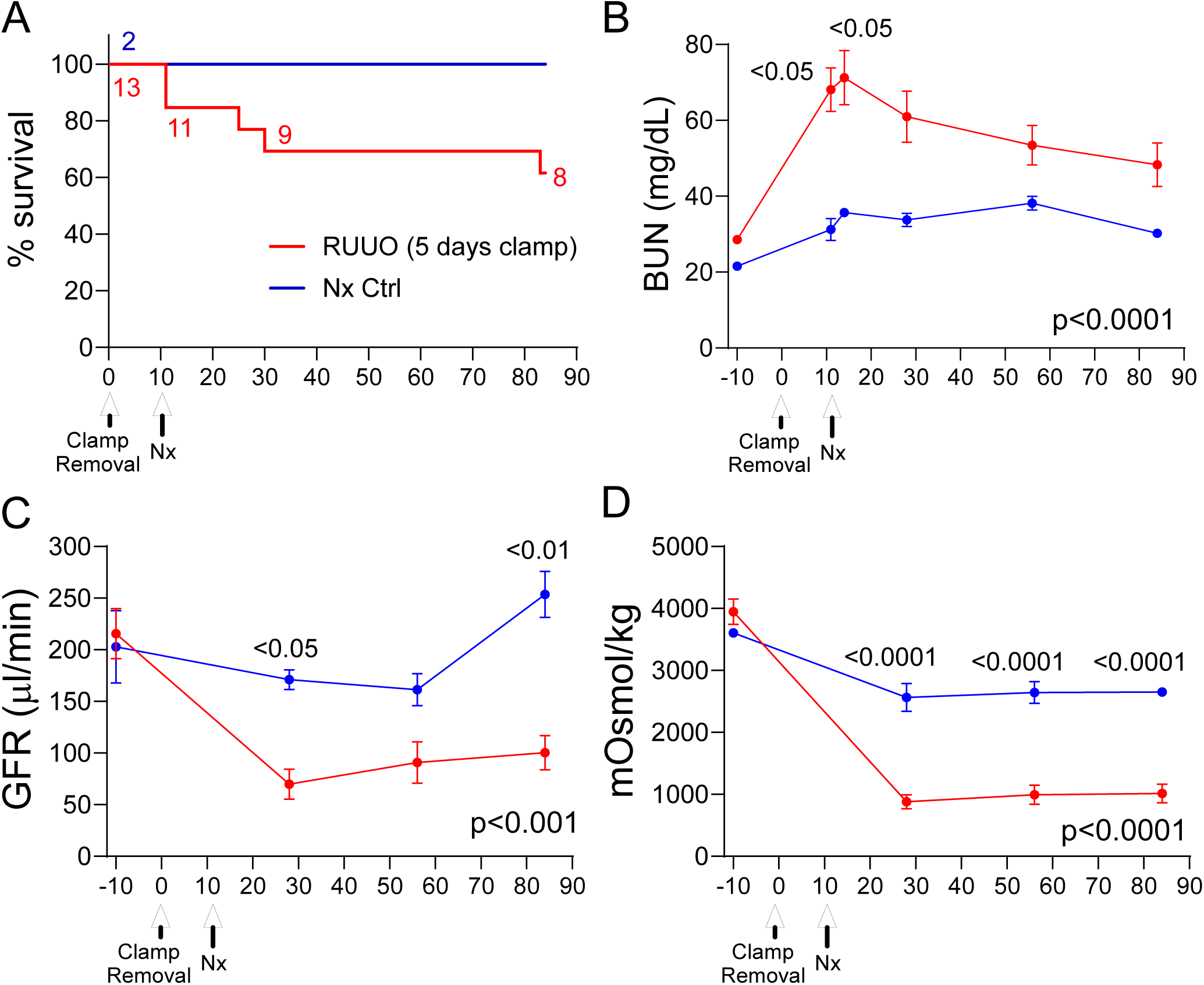
Persistent defect in renal function and urinary concentrating capacity in Six2 Cre; tdTomato mice after R-UUO. Male Six2 Cre; tdTomato mice (mixed background) underwent a 5-day R-UUO followed by contralateral nephrectomy, or nephrectomy alone (Nx). A, Survival, numbers of mice indicated; B, BUN time course after R-UUO; C, tGFR time course; D, Urinary osmolality after 18hr water restriction. Mouse numbers shown in A. Data shown as means +/-SEM. 2-way ANOVA p values indicated. If p<0.05, q values shown for between group comparisons corrected for repeat testing.

**S. Figure 8.**
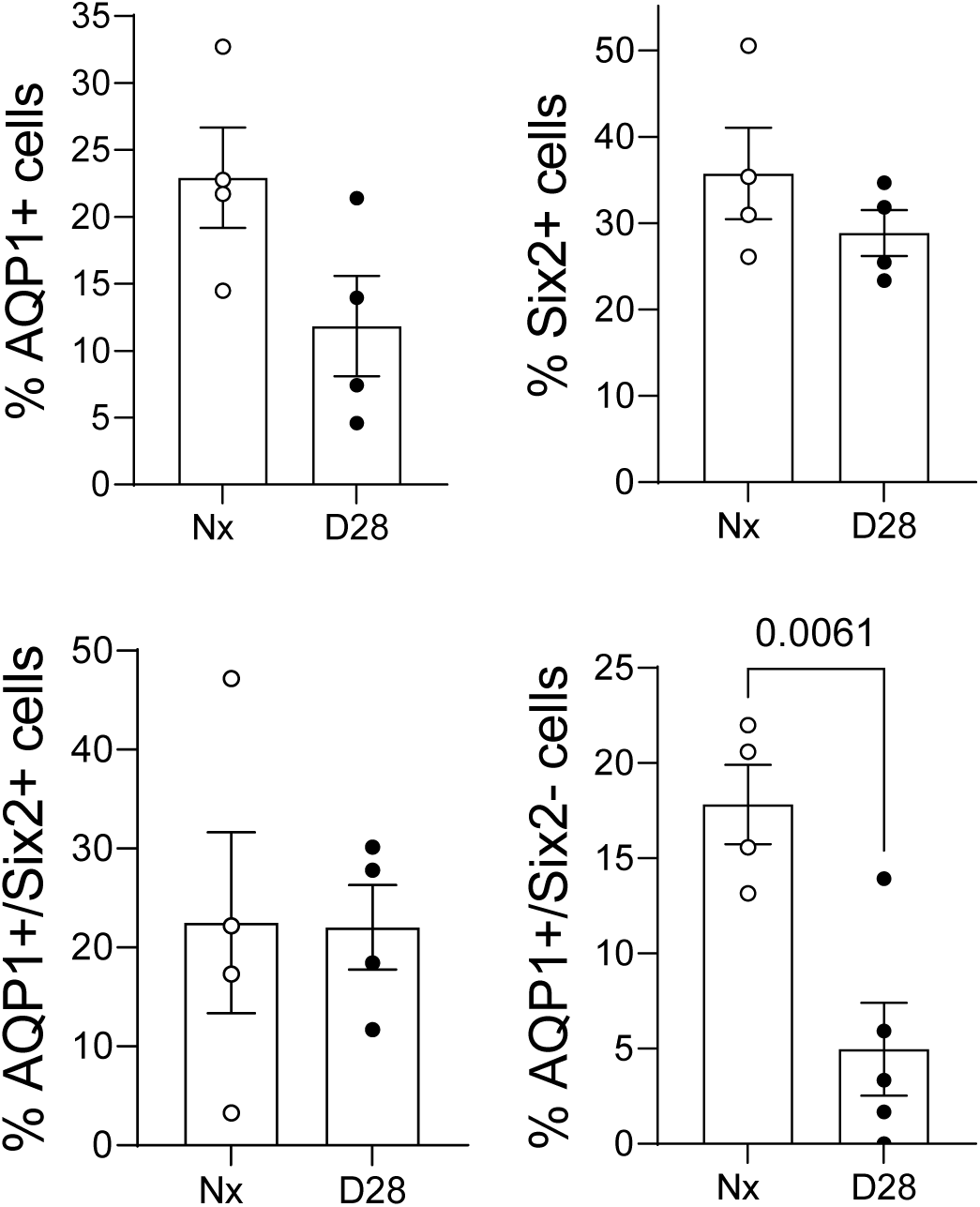
Reduced numbers of Six2 lineage negative, AQP1 positive descending vasa recta cells in the proximal IM after R-UUO. Quantification of Six2 lineage and AQP1 positive and negative cell populations in the proximal IM. Quantification as the % of total cells in each area. Individual data points, means +/- SEM. T-test, p values shown.

**S. Figure 9.**
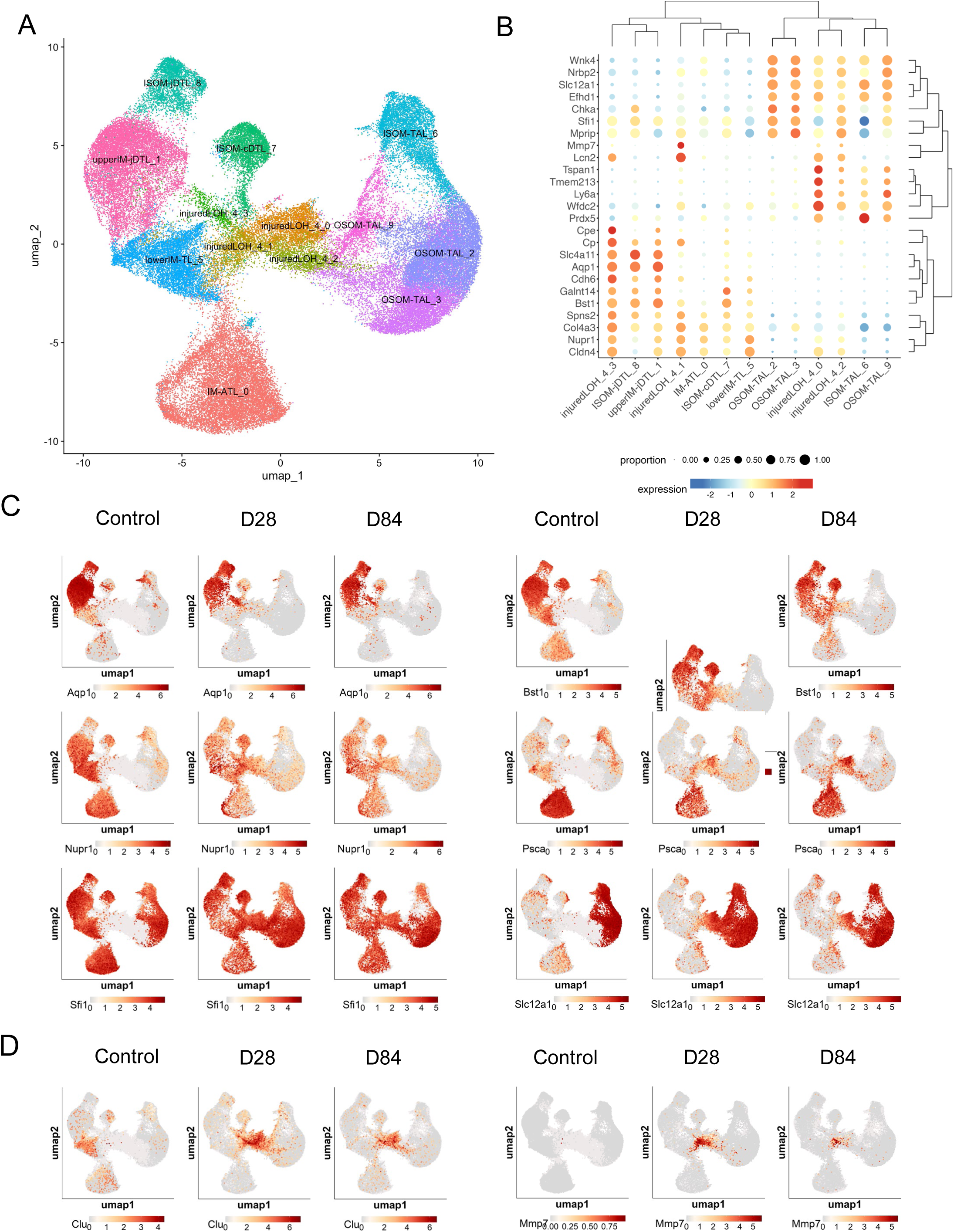
Injured cluster 4 cells are derived from different inner and outer medullary loop of Henle cell populations. A, Clustering of LOH cells identified 4 sub-clusters of injured LOH cluster 4 cells; B, Clustering of the top 25 LOH injured cluster 4 DEGs in other LOH cell populations in controls, day 28 and 84 after R-UUO; C/D, UMAPs showing changes in the distribution of LOH cell segment specific markers (C), and injury markers (D) in controls vs. day 28 and 84 after R-UUO.

**S. Figure 10.**
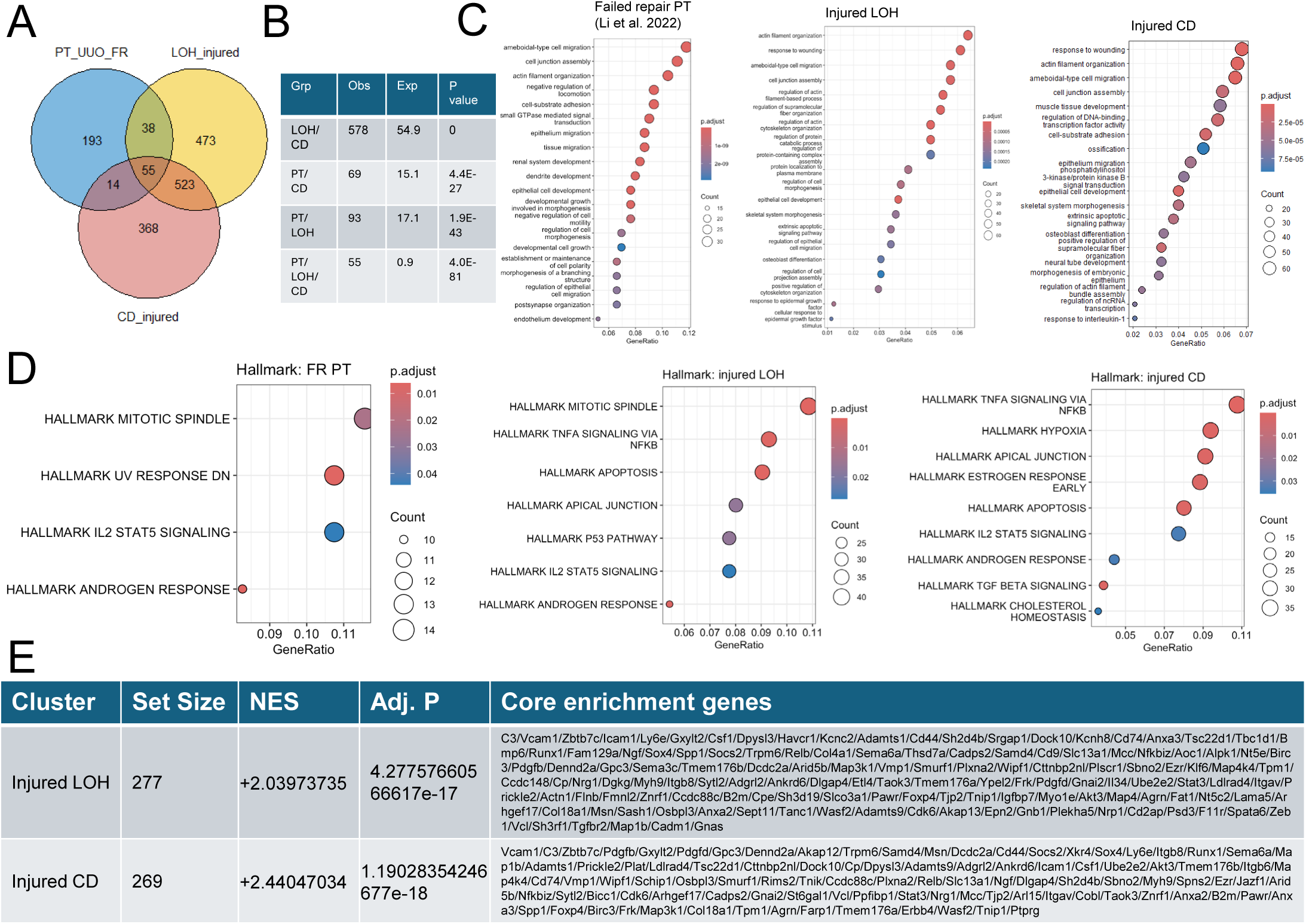
Overlap between failed repair PTECs after I-UUO, and injured LOH and CD clusters after R- UUO. A, Venn diagram illustrating the overlap between published FR-PTECs DEGs, and injured LOH cluster 4 and injured CD cluster 5 population DEGs from these studies; B, Statistical significance between multi-set intersections from the Venn Diagrams determined using the SuperExactTest function in R; C/D, GSEA of DEGs in FR-PTECs, injured LOH and CD populations showing upregulated GO terms (C), and Hallmark terms (D); E, GSEA of DEGs in injured LOH and CD populations with the DEG gene set from FR-PTECs showing set size, normalized enrichment scores (NES). Core enrichment genes indicate genes within the injured LOH and CD cluster DEGs that are driving the NES.

**S. Figure 11.**
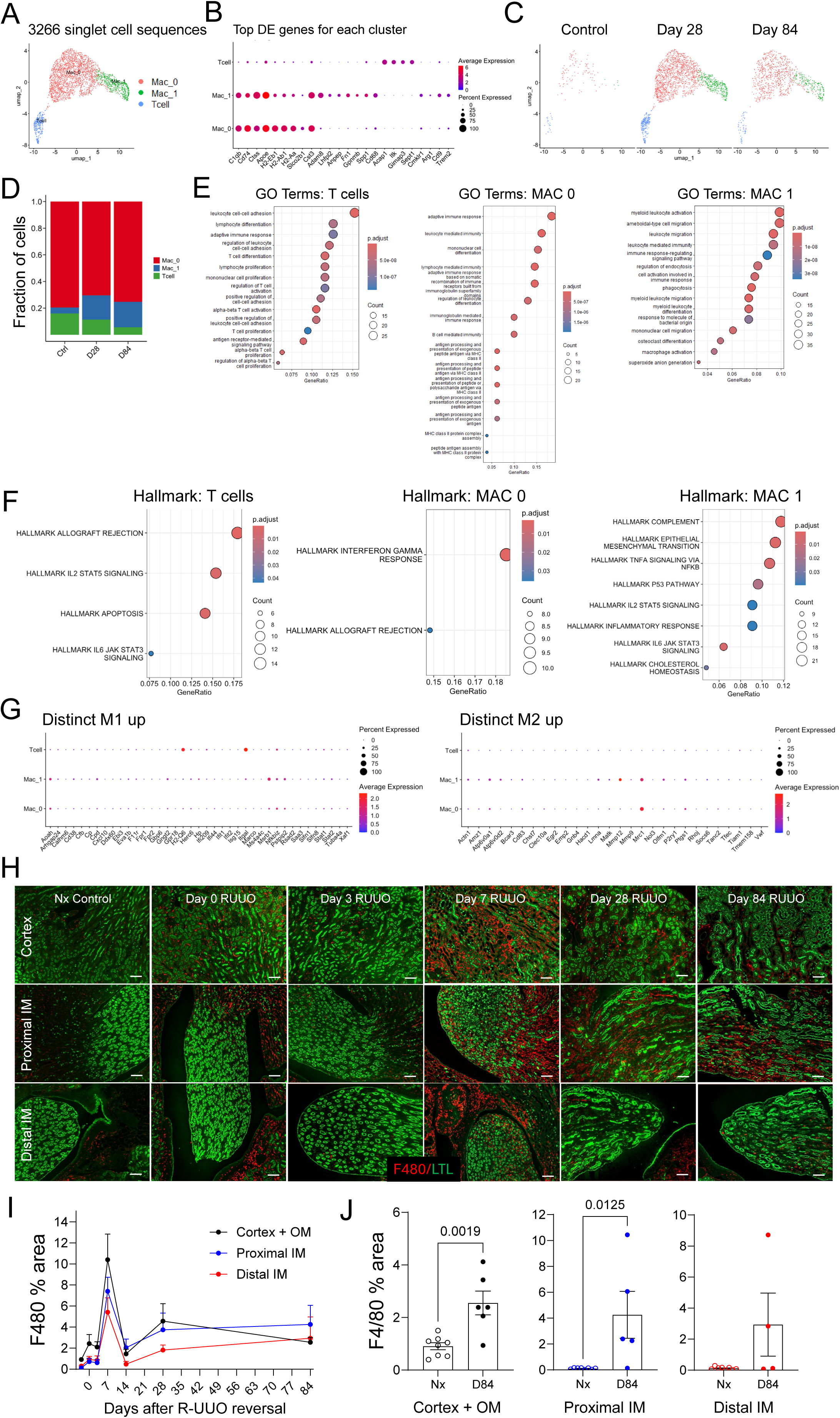
Only a minor long-term increase in inflammatory macrophages in the renal medulla after R-UUO. A, Re-clustering of inflammatory cells identified T cells, MAC0 and MAC1 population in the combined data (A), and at different time points after R-UUO (C); B, Top DEGs in inflammatory cells; D, Fraction of inflammatory cell clusters at different time points; E/F, Gene set enrichment analysis of the top T cell, MAC0 and MAC1 DEGs showing upregulated GO terms (E), and Hallmark terms (F). H, F4/80 antibody and LTL staining in the cortex, proximal and distal IM at different time points after R-UUO, and in Day 84 Nx controls. Scale bars=100um. I/J, Quantification of F4/80 staining (% of the selected area) in the cortex and OSOM, proximal and distal IM lineage cells in the distal IM over time (I), and day 84 after R-UUO (J). Means +/- SEM, individual data points. T tests, p values shown.

**S. Figure 12.**
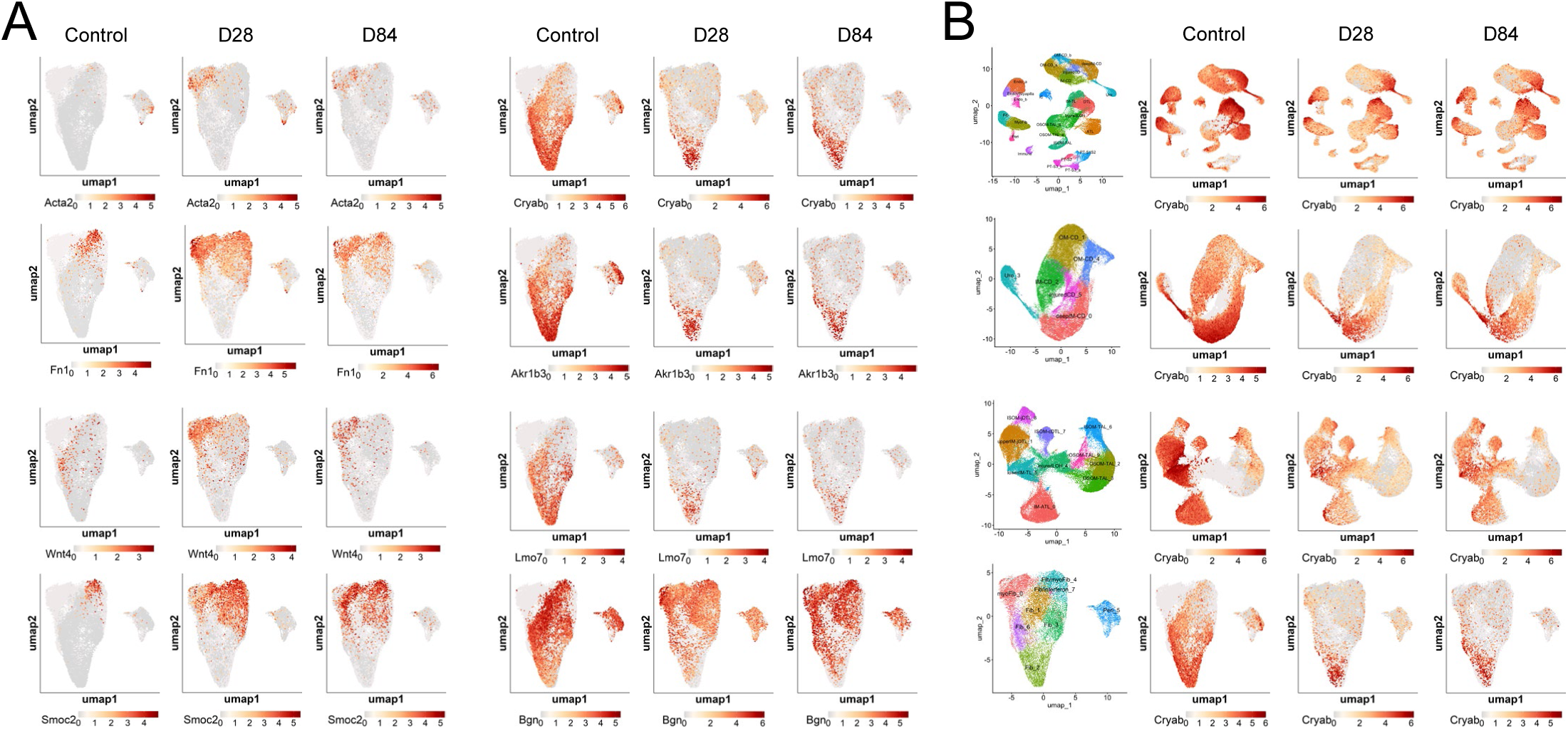
Distribution of fibroblast and inner medullary cell marker genes. UMAPs showing changes in the distribution of fibroblast and IM cell markers in fibroblast clusters (A), and of the IM cell marker, CryAB, in LOH, CD, and fibroblast clusters at different time points after R-UUO (B).

**S. Figure 13.**
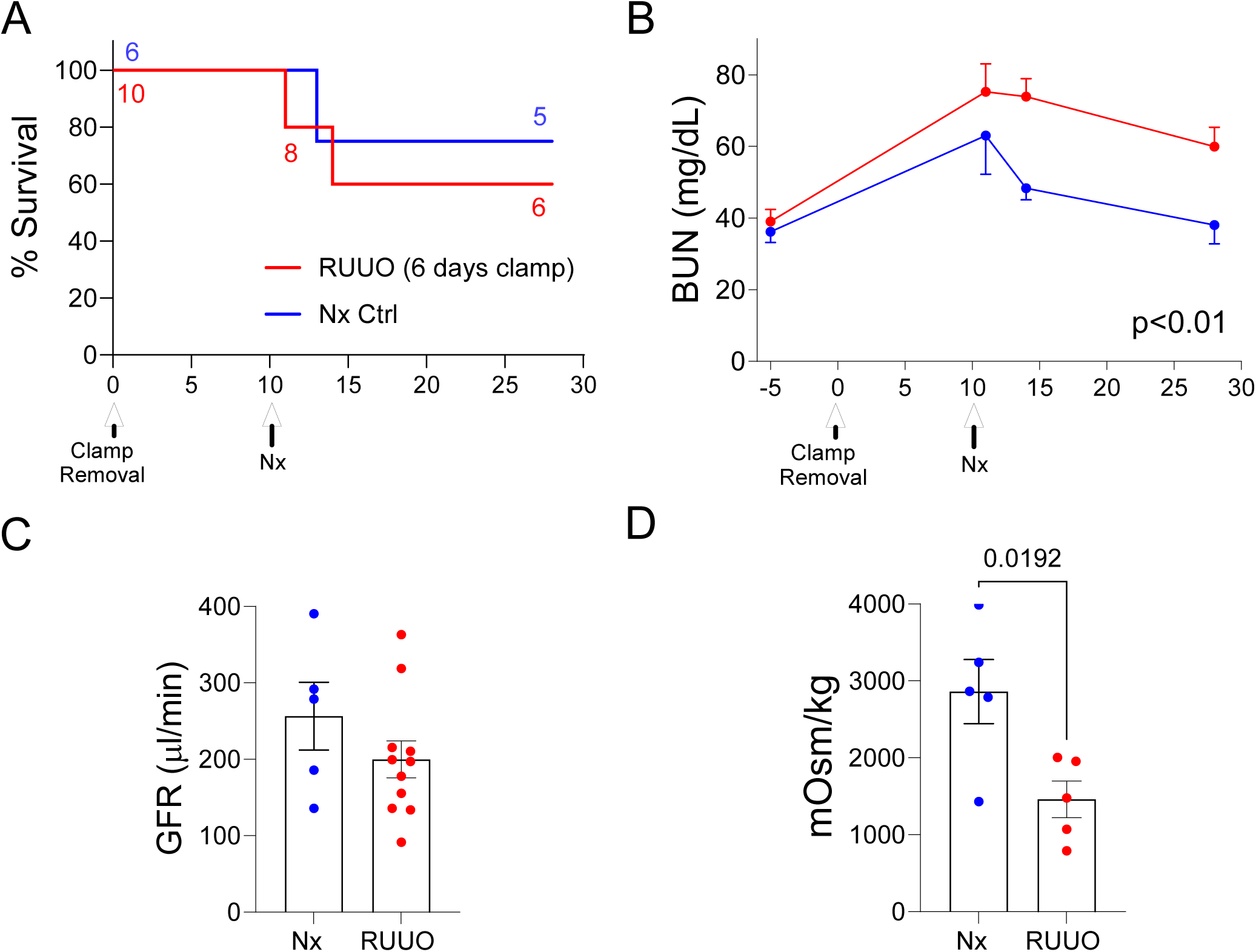
Persistent defect in urinary concentrating capacity in TNC CreERT2; tdTomato mice 28 days after R-UUO. Male TNC CreERT2; tdTomato mice (C57Bl/6 background) underwent a 5-day R-UUO followed by contralateral nephrectomy, or nephrectomy alone (Nx). A, Survival, numbers of mice indicated; B, BUN time course after R-UUO. Data shown means +/-SEM. 2-way ANOVA p values indicated; C/D, tGFR (C), and urinary osmolality after 18hr water restriction (D) 28 days after R-UUO, and in Nx controls. Means +/- SEM, individual data points shown. T test, p value shown.

**S. Figure 14.**
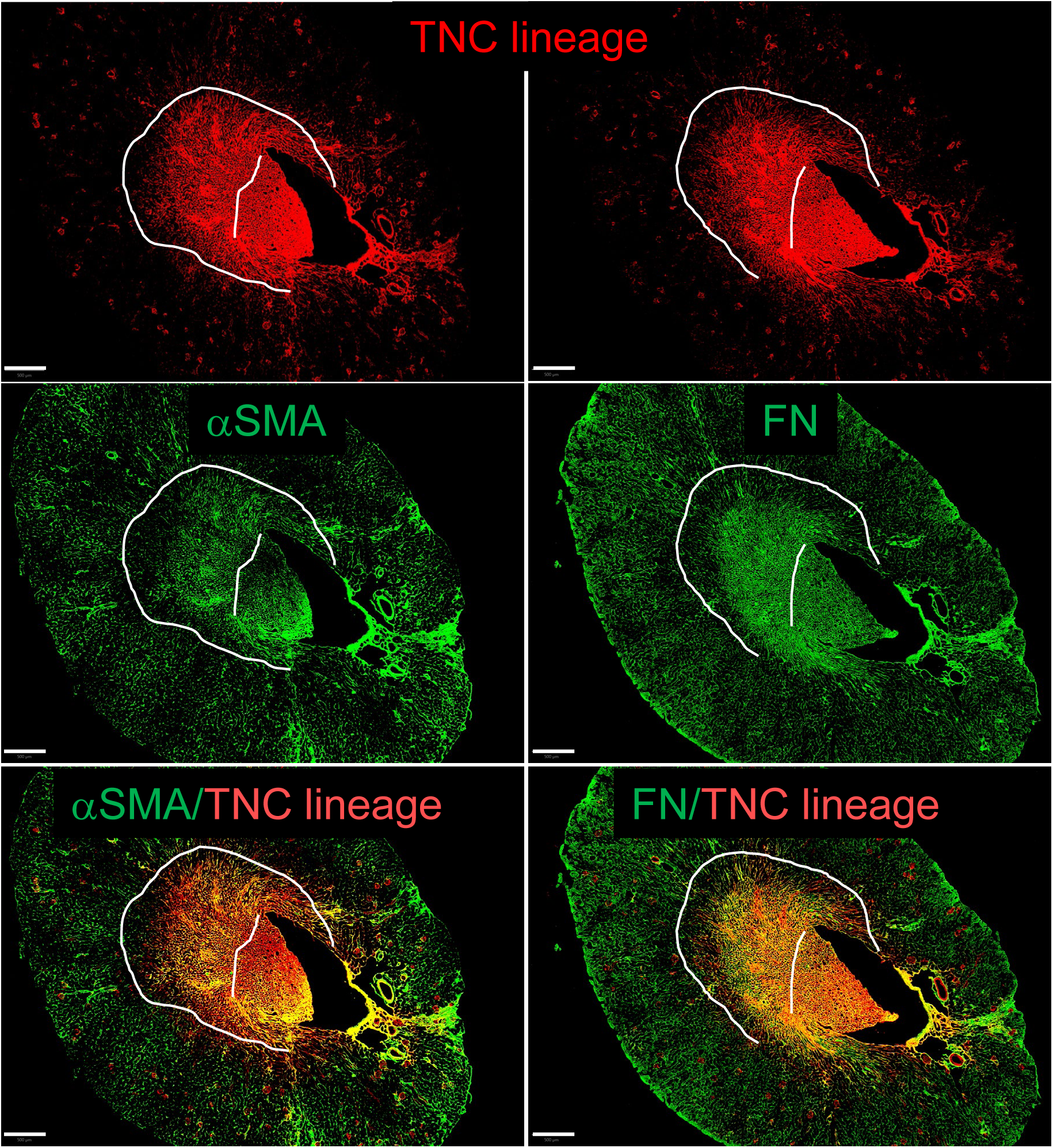
Immunofluorescence images showing overlap between TNC lineage cells and myofibroblasts in the renal medulla. Images showing TNC lineage (tdTomato), and α-SMA or Fibronectin (FN) staining on sequential sections of kidneys from a mouse 28 days after R-UUO. Scale bars=500um, dotted lines indicate the IM/ISOM boundaries.

**S. Figure 15.**
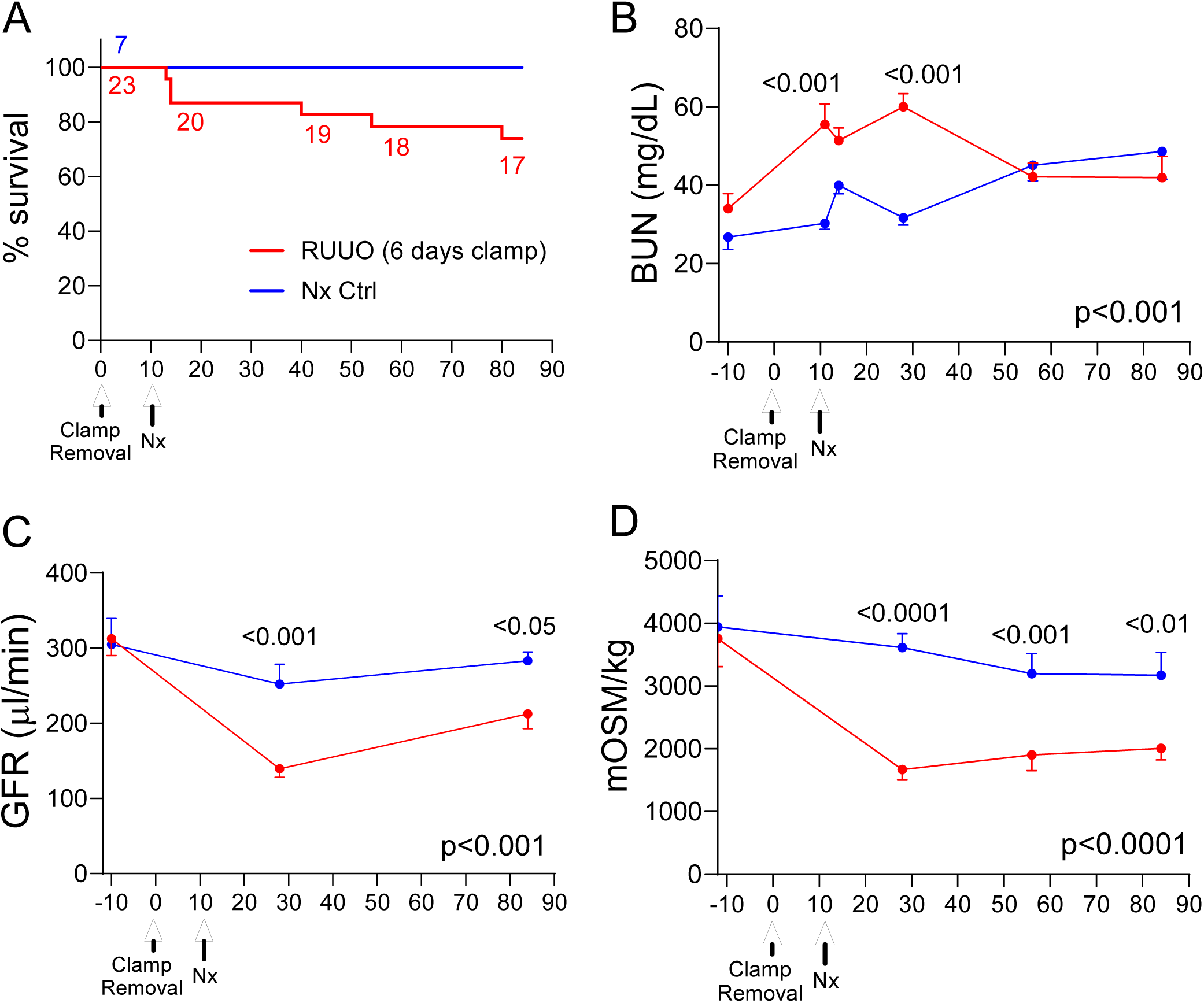
Persistent defect in renal function and urinary concentrating capacity in HoxB7; tdTomato mice after R-UUO. Male HoxB7 Cre; tdTomato mice (mixed background) underwent a 6-day R-UUO followed by contralateral nephrectomy, or nephrectomy alone (Nx). A, Survival, numbers of mice indicated; B, BUN time course after R-UUO; C, tGFR time course; D, Urinary osmolality after 18hr water restriction. Mouse numbers indicated in A. Data shown as means +/-SEM. 2-way ANOVA p values indicated. If p<0.05, q values shown for between group comparisons corrected for repeat testing.

**S. Figure 16.**
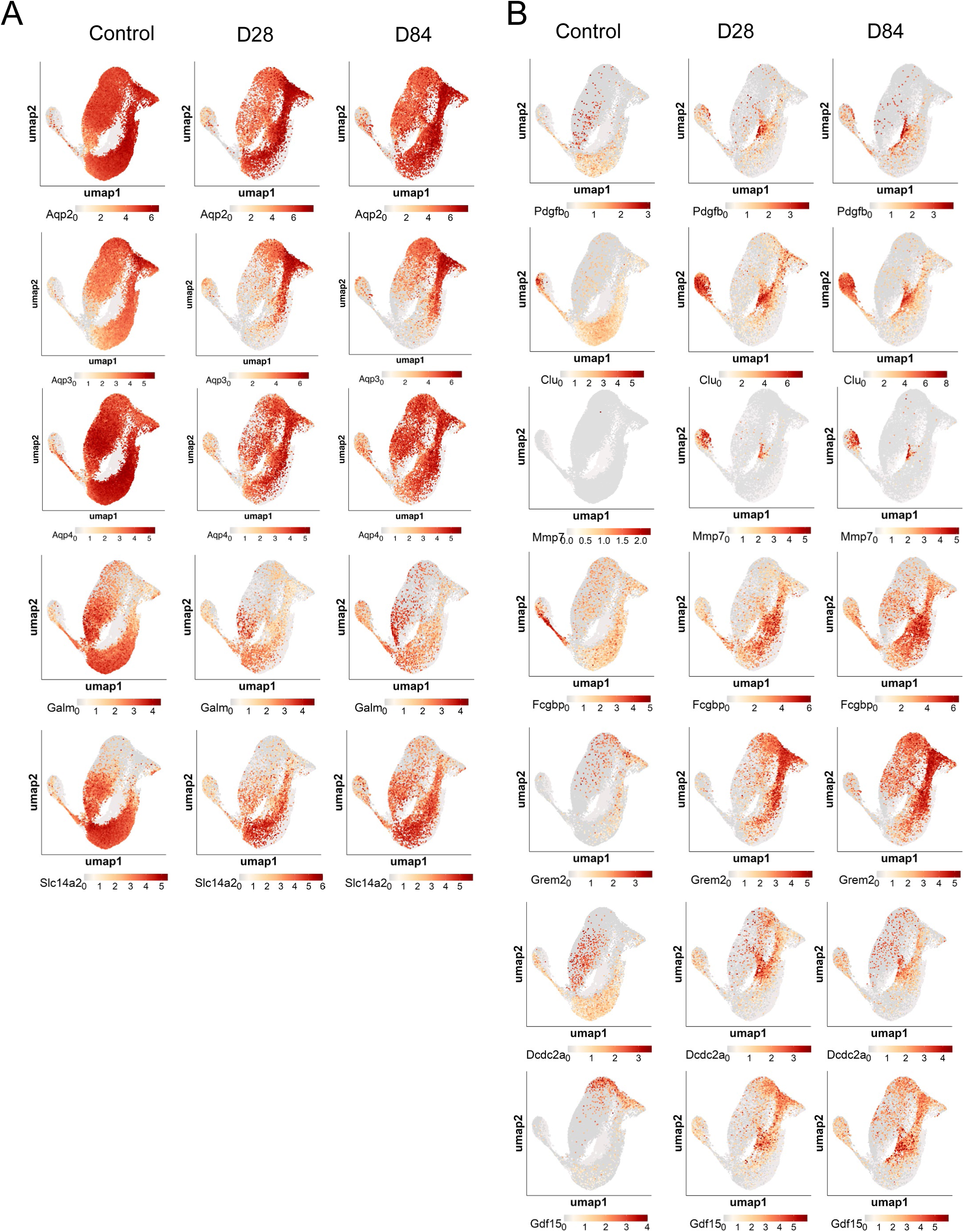
Distribution of CD differentiation and injury markers after R-UUO. UMAPs showing changes in the distribution of CD cluster differentiation (A), and injury (B) markers at different time points after R-UUO.

**S. Figure 17.**
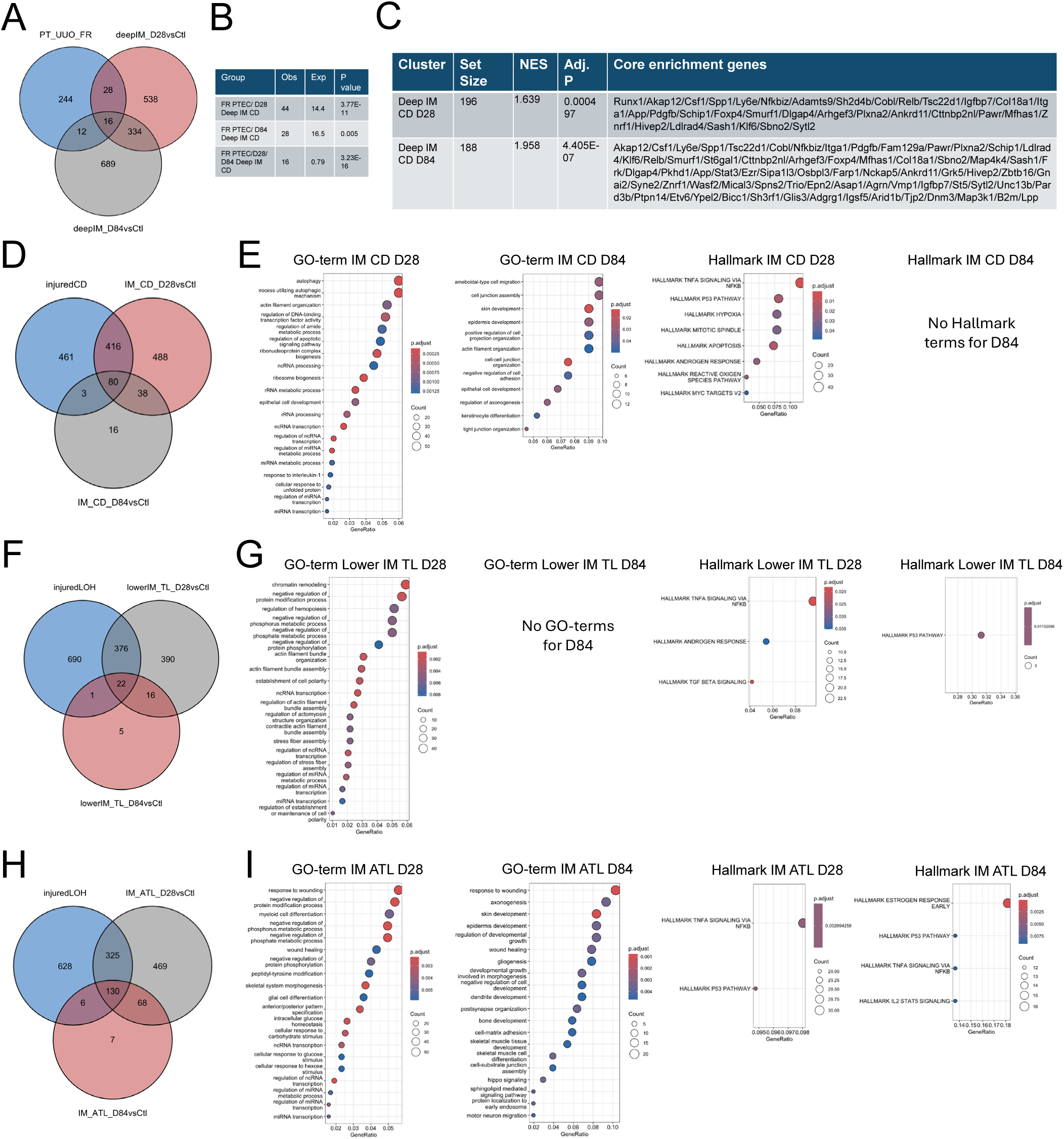
Similarities and differences different CD and LOH populations after R-UUO. A-C, Venn diagram showing the overlap between DEGs from FR-PTECs after I-UUO and deep IM CDs after R-UUO (A), statistical significance between multi-set intersections from the Venn Diagram (B), and GSEA of DEGs in Deep IM CD cells after R-UUO with the DEG gene set from FR-PTECs after I-UUO showing set size, normalized enrichment scores (NES). Core enrichment genes indicate genes within Deep IM CD day 28 and 84 cluster DEGs that are driving the NES; D/F/H, Venn diagrams showing the overlap between injured CD cluster 5 cells and IM CD cluster 2 cells after R-UUO (D), injured LOH cluster 4 and lower IM TL cells after R-UUO (F), and between injured LOH and IM ATL cells after R-UUO (H); E/G/I, statistical significance between multi-set intersections from the respective Venn Diagrams; L/G/I, GSEA showing upregulated GO and Hallmark terms in DEGs in IM CD cells (E), lower IM TL cells (G), and IM ATL cells (I) after R-UUO.

**S. Figure 18.**
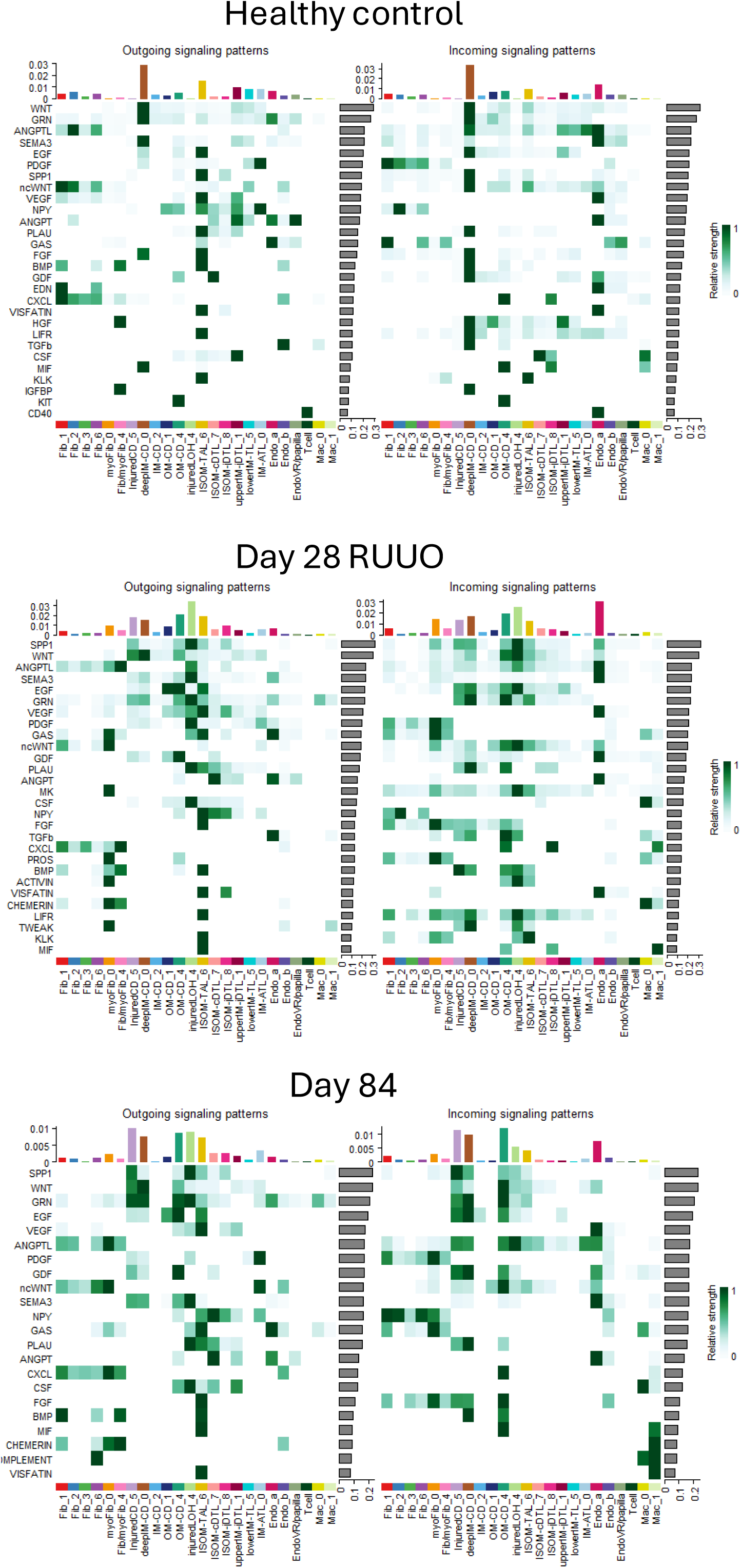
Long-term effects of R-UUO on cell-cell communications in the renal medulla. Cell-cell communication analysis was performed to identify ligand-receptor pairing of the major fibroblast, LOH, CD, endothelial and inflammatory cell clusters in renal medullas from controls, day 28 and day 84 after R-UUO. Outgoing ligand secretion is shown in the left-hand panels, incoming cognate receptors for the same pathway is shown in the right-hand panels. Contribution of individual cell clusters to the signaling patterns is indicated by the horizontal bar charts, relative strength of the response indicated with color coding and from the vertical bar charts.

