## Supplemental methods and figures for "Partial repair causes permanent defects in papillary structure and function after reversal of urinary obstruction"

#### Supplemental Methods 1

##### Mouse lines and strains

Male BALB/c mice were purchased from Charkes River. HoxB7 Cre transgenic mice (Tg(HoxB7-Cre)13Amc),[1] and Six2 eGFP-Cre BAC transgenic mice (Tg(Six2-EGFP/Cre)1Amc/J), [2] were kindly provided by Andy McMahon from USC Keck School of Medicine, but are also available as frozen stock at Jackson labs. tdTomato Cre reporter mice, Tenascin C Cre-ERT2-IRES-GFP knock in mice,[3] were kindly provided by Dr. Agnes Fogo from Vanderbilt University Medical Center with permission from Chuan-Ming Hao from Fudan University in Shanghai, China. B6.Cg-*Gt(ROSA)26Sor*$^{tm14(CAG-tdTomato)Hze}$/J, [4] were obtained from Jackson labs. Six2 and HoxB7 Cre mice are on mixed C57Bl/6, CD-1/Swiss Webester backgrounds, Tenascin C Cre-ERT2-IRES-GFP knock in and tdTomato Cre reporter mouse lines were on a C57Bl/6 background. Mice were genotyped and identified from each punch biopsies using allele-specific primers performed by Transnetyx. Genotyping primers are listed in S. Table 1.

Reversible unilateral ureteral obstruction.

Mice were maintained on a 12:12-h light-dark cycle with free access to food and water. Euthanasia was performed by cervical dislocation after anesthesia with inhaled isoflurane at the end of each experiment, or at humane end points. For all injury models, body weight was monitored throughout the study. R-UUO was performed in 12–13-wk male mice. For this, mice were anesthetized with ketamine/xylazine mixture (120–150 mg/kg ketamine and 12–15 mg/kg xylazine), and mice placed in a prone position. The left kidney was exteriorized through a dorsal incision, and after dissecting away lower pole fat and connective tissues, a small vascular clamp was applied to the left ureter directly below the pole (Fine Science Tools 00396-01 5-15g 7mm clamp), using applying forceps (Fine Science Tools 00071-14). The kidney and clamped ureter were then gently pushed back into the retroperitoneal space. After a variable interval of time depending on the mouse strain (between 5 and 7 days), ureteric obstruction was reversed. For this, mice were anesthetized again, the left kidney exteriorized, and the vascular clamp carefully removed. The connective tissues surrounding the clamp tissue has to be peeled away from the clamp carefully, and the clamp is opened slowly using the clamp applying forceps. For long-term studies, a contralateral nephrectomy was performed 10 days after reversal of the obstruction. For the nephrectomy surgery, mice were anesthetized using inhaled isoflurane, the right kidney exteriorized, the renal pedicle was tied off with silk suture, and the kidney removed. Nephrectomy only controls were performed at the same age/time point as the long-term R-UUO studies (e.g.: day 84 Nx control, Nx performed age ~14-16wks, and mice euthanized at 26-28wks). All mice are also given 0.5ml of 0.9% sodium chloride IP post operatively, and analgesia provided with SC buprenorphine for 48 h after each surgery. Surgeries were started in the morning and completed by noon.

Assessment of renal function. Blood was collected by submandibular vein or cardiac puncture into lithium-heparin-coated microcuvette tubes (Braintree Scientific # MV-CB300 16443-BX), for long term studies, which may include Days 11 (the day after nephrectomy), 14, 28, 56 and 84 after R-UUO, depending on the study. Plasma was collected for BUN and measured in 5 microliters in duplicate, according to the manufacturer's instructions (Bioassays Systems DIUR100). Serum creatinine was measured in 5microliters of plasma in duplicate by LC-MS/MS, [5] at the O'Brien Core Center for AKI Research of the University of Alabama. Transdermal glomerular filtration rates (tGFR) were measured in conscious mice, as described [6]. The FITC-sinistrin half-life was calculated using a three-compartment model with linear fit using MPD Studio software (MediBeacon, Mannheim, Germany). The FITC sinistrin half-life was converted to tGFR (in ml/min) with correction for mouse body weight, as described [6]. For urine osmolality studies, mice were water deprived for 18hrs, and placed on a 96 well plate inside a customized Perspex box. Urine is collected from the wells with a pipette and placed in a microcentrifuge tube and centrifuged for 10 minutes at 10000 rpm to separate debris. Clean urine is then placed in clean microcentrifuge tubes and stored at -20C. Urine osmolality is measured in batches using a Precision Systems 6002 Touch Micro OSMETTE 30 uL Osmometer.

Tissue harvesting. Mice were anesthetized using inhaled isoflurane, and a midline incision made from the abdomen to the sternum to expose the heart, a 27g needle butterfly catheter inserted into the left ventricle at the apex of the heart, and the lower abdominal aorta cut. The cardiovascular system was flushed with cold phosphate buffered saline (PBS) until the organs were pale. The left kidney was initially checked for obstruction, and if uncertain, methylene blue injected into the renal pelvis to determine whether the left ureter was patent. Since external evidence of hydronephrosis resolves within ~24 hrs of reversal of obstruction, if there was evidence of persistent obstruction at any time point after reversal, data for that mouse was discarded. If collecting kidney samples for RNA, a clamp was placed on the renal pedicle, the renal capsule removed, and a small transverse section from the center of the kidney cut and snap frozen. If RNA was not being collected, the PBS was replaced with 10% normal buffered formalin (NBF) (Fisher Scientific StatLab™ # 28600-1), until the body becomes stiff, the renal capsule was removed, and two small transverse sections from the center of the kidney cut and soaked in 10% NBF at room temperature for another 1hr and 4hrs for formalin fixed frozen (FFF), and formalin fixed paraffin embedded (FFPE) tissues, respectively. For FFF samples were washed in PBS, placed in 30% sucrose in PBS overnight, and mounted in Tissue-Tek OCT (Sakura), and stored at -80C for subsequent staining. For FFPE samples, blocks were washed in PBS, stored in 70% ethanol, and processed in paraffin for staining.

Immunofluorescence staining and analysis. FFF and FFPE sections were prepared, washed, blocked, and primary and secondary antibodies applied, as described (see S. Methods for more details on staining, and S. Table 1 for the list of antibodies, fluorophores, and conditions).[7] The genetically encoded tdTomato fluorophore was quenched in FFPE samples, but detected directly in FFF sections in studies indicated by Figs. 3F/H and 6F, and with anti-tdTomato antibodies in all other studies. GFP was quenched in FFPE samples and detected by amplification using anti-GFP antibodies in FFF samples using AF488 conjugated secondary antibodies. To quantify immunofluorescence (IF) images, digital images were scanned using Zeiss AxioScan Z1 slide scanner with DAPI, AF488, AF555, AF 647 and Cy7 filters on (Carl Zeiss Microscopy GmbH, Oberkochen, Germany, 10X), and digital images downloaded into QuPath (version 0.5.0) for analysis. For quantification, kidney regions of interest (ROIs, cortex, outer stripe of the outer medulla (OSOM), inner stripe of the outer medulla (ISOM), proximal approximately 1/3 and distal approximately 1/3 of the inner medulla (IM) were demarcated using the QuPath annotation tool on whole kidney scanned images using established landmarks depending on the experiment to create areas of interest including using juxta medullary glomeruli to demarcate the cortex/OSOM junction, LTL staining the OSOM/ISOM and the ISOM/IM junctions; and THP-1 staining to demarcate the ISOM/IM junction. If sections through the IM were less than 1mm in length because of the angle of the section, we would only quantify the area of interest in the proximal IM (which is why there are fewer datapoints for most of the distal IM analyses). Using QuPath pixel classifier settings, we created individual pixel classifiers for each of our stains. Cell numbers and cell types in different kidney areas were identified based on average values of fluorescence staining using machine learning and thresholding detection using QuPath. The pixel classifier was trained with a minimum of 30 training regions for the cell (DAPI stained nuclei) and cell type (e.g.: HoxB7 lineage tubules) and identified as areas of fluorescence above background levels (nucleus threshold, segmentation parameters, cell expansion, smoothed features). Once the machine learning classifier was created, the marker was quantified in the region of interest. These object classifiers were either run by themselves or in combination (e.g.: tdTomato and AQP1 staining overlays). By running classifiers in combination, we quantified the ratio of cells carrying one or both of these markers (e.g.: Six2 and AQP1 in Figure 3H/I). As indicated in the figures, images were either quantified as surface area stained in the ROI, or as the ratio of cells staining with the indicated markers as a fraction of the total cells in that ROI using Hoechst 33342 staining to identify nuclei.

To quantify collecting duct surface areas, digital images of HoxB7 lineage-labelled kidney sections after IF staining with LTL, tdTomato, and AQP2 antibodies with IM lengths of $\geq$1mm were used. Distal and proximal IM were defined as outlined above, as well as high and low LTL labeling, and CD areas were defined by staining with the HoxB7 lineage marker, tdTomato and AQP2. To quantify tubular surface areas in our mice, we only evaluated circular-shaped CDs defined as having major/minor axis ratios of >0.8, as previously described.[8] We quantified the surface area of each tubule and its lumen using the freehand selection tool in Fiji, 15 and 30 tubules in each area of interest, and the number of Hoechst 33342 stained nuclei counted in each tubule cross section. To quantify tubular numbers, CDs were quantified in 3 145X145mm ROIs with at least 5 tubules per ROI, in distal and proximal IM areas of interest, as defined above.

Histological staining and scoring. FFPE sections were staining with Periodic Acid Schiff (PAS) to evaluate morphology, and fibrosis/collagen deposition determined after staining with Sirius red (SR), by a blinded observer (R.D.). This was performed using an Olympus BX-41 microscope equipped with a polarized light filter, using ImageJ to quantify birefringent SR-stained collagen fibrils areas/total surface areas from digitally captured images, as described [9].

Single cell RNA sequencing (scRNA seq). Papillae were dissected at two times points of injury (d28, d84 after R-UUO) and from healthy controls. Three to four papillae from the same time point were placed in a cryogenic vial and flash frozen with liquid nitrogen, then stored at -80C. Each frozen tube was fixed and dissociated into single cells using the 10x Genomics Chromium Next GEM Single Cell Fixed RNA Sample Preparation Kit #1000414 (for details see S. Methods 3). Cell concentration was determined using the Countess III FL Automated Cell Counter (VANTAGE core) and propidium iodide staining solution, Invitrogen #P3566. For RNA sequencing, dissociated cells from each of the pooled samples were individually hybridized with bar coded genome-wide probe sets from 10X Genomics Flex kits according to the manufacturer's instructions. After probe hybridization and construction of libraries, samples were sequenced using an Illumina Novoseq 6000 PE150 sequencer, and bar codes deconvoluted to provide individual sample datasets.

Bioinformatics Pipeline and Analysis of transcriptome data. Gene expression matrices were generated using CellRanger software (10x Genomics), and raw data reads were processed in R. Cells then underwent quality control steps before downstream analyses. No differences in cell cycle scores were observed (data not shown). Cells with fewer than 300 detected genes or more than 10% mitochondria reads were excluded. Subsequently, we applied SoupX to remove ambient RNA.[10] Briefly, ambient RNA fraction was estimated from the empty droplets using the autoEstCont method, followed by the adjustCounts function to remove ambient RNA counts. In parallel, doublets were removed using scDblFinder.[11] Cells that had gene signature of mixed cell type were also removed as doublets. Count data were normalized using the NormalizedData function in the Seurat package,[12] and highly variable genes were identified with FindVariableFeatures function. Data were then scaled using the ScaleData function. Dimensionality reduction was performed using Principal Component Analysis (PCA) and visualized with uniform manifold projection (UMAP). Cell population identities were identified using published anchor genes for renal medullary loop of Henle (LOH), collecting duct (CD), fibroblast, endothelial cell (EC), and immune cell subclusters. [13-15] Unsupervised clustering was performed using the FindNeighbors and FindClusters functions. The number of principal components (PCs) was determined using the ElbowPlot function. Differential gene expression analysis between clusters and timepoints was carried out using the FindAllMarkers function and the FindMarkers function, respectively. GO and Hallmark pathway gene set enrichment analysis (GSEA) was performed and visualized by ClusterProlifer, as well as published FR-PTEC DEG gene sets from published data, [16] as described in the text. Enrichment scores were calculated by comparing the log fold change in each gene in the gene set with all other expressed genes. This represented as the normalized to the gene set size for each gene set to generate normalized enrichment scores (NES). All samples were combined and integrated using the IntegrateLayers function followed by UMAP reduction, clustering and differential gene expression analysis as above. The subset function was used to extract clusters of each cell type (LOH, CD, fibroblast, EC, and immune cells), on which PC analysis, re-clustering, and differential gene expression analysis were carried out. The injured LOH cluster 4 was subclustered using the FindSubCluster function. Cell-cell communication analysis was performed using CellChat. [17]The probabilities of secreted signaling pathway were obtained for different cell clusters based on expression levels of ligands or receptors within these clusters. The potential Ligand-receptor interactions was inferred and visualized using the netAnalysis_signalingRole_heatmap function. Data were visualized using Seurat, dittoSeq, and ShinyCell R packages.[18]

Statistical analysis. were performed using GraphPad Prism 10.1.2 software (San Diego, CA), and results expressed as means +/- SEM, with individual datapoints shown where indicated. Two-tailed, unpaired T-Tests were used to compare two groups, 1-way ANOVA to compare between 3 or more groups, and 2-way ANOVA to compare groups over time. If p values were <0.05 for between group analyses, multiple between group comparisons were performed controlling for false discovery rates (FDR) of p<0.05 using the two-step method of Benjamini, Krieger and Yekutieli. Q values are indicated after correction for multiple between group comparisons. Statistically significance between multi-set intersections from the scRNA-Seq subcluster Venn Diagrams was performed using the SuperExactTest function in R.[19]

#### Supplemental Methods 2

##### Immunofluorescence Staining Protocol on Mouse Kidney Tissue in the de Caestecker Lab

##### Reagents
Citrisolv
100% Ethanol
dd$H_2O$
10X IHC Antigen Retrieval Solution 10X low pH (Invitrogen, 00-4955-58)
1X Phosphate Buffered Saline (PBS)
10% buffered formalin (Cardinal Health, C4305-12)
Hydrophobic Barrier Pen (Vector Labs, H-4000)
Glycine (Sigma-Aldrich, 410225)
Avidin/Biotin Blocking Kit (Vector Labs, SP-2001)
10X Power Block Universal Blocking Reagent (BioGenex, HK085-5K)
Hoechst 33342 (20mM, Thermo Fisher, 62249)
100% Glycerol (Fisher Scientific, G33)
Mouse on Mouse Kit (Vector Labs, BMK-2202
Mouse Serum (Millipore, S25)
Rabbit Serum (Sigma, R9133)

##### Equipment
Steamer for antigen retrieval
Microwave or pressure cooker can be used
Moisture Chamber
100-Slide Storage Box (Fisher Scientific, 03-448-1)
Kimwipes, 8.4 in x 4.4 in (Fisher Scientific, 06-666)
dd$H_2O$ in a wash bottle

##### Procedure
###### De-Paraffinization and antigen retrieval for FFPE sample
1) De-wax and hydrate paraffin sections.
2) Transfer slides to 1X PBS.
3) Dilute 10X citrate buffer to 1X using dd$H_2O$.
4) Place slides in buffer and cap the container holding the slides. Steam for 26 minutes.
5) Remove slides from steamer. Remove the cap and allow slides to equilibrate to room temperature.
6) Pour citrate buffer into appropriate waste container and wash slides in 1X PBS for 3 minutes three times.

###### Frozen sections
1) Fill Coplin jar with 10% buffered formalin
2) Place slides directly from the-80C into the formalin for 5 minutes
3) Remove slides and transfer to 1X PBS
4) Wash sections in 1X PBS 4 times for 3 minutes

###### Immunofluorescence staining
1) Using a hydrophobic pen, draw a large barrier around each section. Do not allow pen to touch section.

2) Place slides in humidified chamber and cover section in PBS by using a pipettor so that they do not dry.
3) To remove solutions from sections that are in the humidified chamber, tip the solution off of the slide into the chamber or aspirate them off.
4) Optional: If using Biotin Amplification:
   Block sections in avidin solution for 15 minutes.
   a) Wash sections in PBS twice.
   b) Block sections in biotin solution for 15 minutes.
   c) Wash sections in PBS twice.
5) Block sections for 1 hour with 1X Universal Blocking Reagent (UBR) at room temperature.
6) Dilute 10X blocking reagent to 1X using 9-parts 1X PBS to 1-part UBR.
7) Optional: If using mouse antibodies on mouse tissue
   a) Use M.O.M Kit to make mouse block
   b) Block for one hour
   c) Dilute primary antibodies in M.O.M Protein Concentrate (7.5ml 1X PBS with 600uL concentrate)
8) Dilute primary antibody to desired working concentration in UBR during blocking step.
9) Add diluted antibody to section and incubate overnight at 4℃.
10) Remove solution, and wash sections with PBS for 3 minutes three times.
11) If primary antibody is directly conjugated with a fluorophore, skip to #14.
12) Optional: If using two sequential antibodies of the same species that are not directly conjugated (ex. Rabbit on rabbit)
    a) After sections are washed, dilute rabbit serum at a concentration of 1:200 in 1X PBS and allow to incubate for 1 hour.
    b) Remove solution and wash sections 3 times for 5 minutes
13) If using indirect immunofluorescence, dilute fluorophore-conjugated secondary antibody using Antibody diluent buffer.
14) Add antibody solution to sections and incubate for 60 minutes at room temperature.
15) Remove solution, and wash sections with PBS for 3 minutes three times.
16) If a biotin-conjugated primary or secondary antibody was used:
    a) Dilute fluorophore-conjugated Neutravidin in blocking reagent at 1:500 for 1 hour.
    b) Wash sections in PBS for 3 minutes three times.
17) Incubate sections in Hoechst 33342 (1:5,000 dilution of 20mM solution in 1X PBS) for 5 minutes.
18) Remove Hoechst 33342, and wash sections with PBS for 3 minutes twice.
19) Mount slides in 50% glycerol in PBS. Do not seal the coverslips as they will need to be removed later (note that because they are not sealed, slides have to be kept horizontal, to prevent the coverslip from falling off, and in the humidified chamber)
20) Scan Image
21) Store slides at 4℃ in a moisture chamber.

Supplemental Methods 3

#### Tissue Fixation & Dissociation for Chromium Fixed RNA Profiling in the de Caestecker Lab

##### Reagents and consumables

10x Genomics Chromium Next GEM Single Cell Fixed RNA Sample Preparation Kit (includes Conc. Fix & Perm Buffer, Conc. Quench Buffer, and Enhancer), 10x Genomics #100414
Formaldehyde 37%, Fisher #BP531-25
Liberase TL, Sigma #5401020001
Gibco RPMI 1640, Thermo #11875093 or Storeroom
Gibco PBS 1x without calcium and magnesium, Storeroom #10010-023
Nuclease-free water, Storeroom #P1195
Sterile water, VU Research Pharmacy iLab
Pyrex Reusable Petri Dishes, Fisher #08-747B
Single edge blades, Fisher #12-640
Micro jewelers dissecting forceps, Fisher #13-820-078
High precision micro dissecting scissors, Fisher #08-953-1B
Pipette tips 1000 ul, Fisher #13-889-157
Pipette tips 200 ul, Fisher #13-889-155
2 ml low protein binding tubes, Thermo #88379
5 ml centrifuge tube, Fisher #50-969-831
30 um mesh cell strainers, Fisher #NC0922459
1 ml syringe with plunger, Fisher #14-555-462
Propidium iodide staining solution, Invitrogen #P3566
Countess III

##### Preparations of buffers

All buffer preparations should be fresh (1 ml)

| *Fixation buffer- maintain at room temperature* | Stock | Final | Per 25 mg tissue (ul) |
|---|---|---|---|
| Nuclease-free water | - | - | 791.9 |
| Conc. Fix & Perm Buffer | 10X | 1X | 100 |
| Formaldehyde | 37% | 4% | 108.1 |

| *Quenching buffer- maintain at 4 °C* | Stock | Final | Per 25 mg tissue (ul) |
|---|---|---|---|
| Nuclease-free water | - | - | 875 |
| Conc. Quench Buffer | 8x | 1x | 125 |

*Dissociation Solution prepare 2 ml*

- Reconstitute 5 mg Liberase TL by adding 1 ml sterile water. Agitate at 2-8 °C until dissolved. Store stock solution in single-use aliquots at-20 °C.
- Prepare RPMI + 0.2 mg/ml Liberase TL (Add 80 ul Liberase TL stock solution into 1,920 ul of RPMI, mix, maintain at 4 °C)
- Warm dissociation solution for 10 minutes at 37 °C before use.

#### Protocol Overview

##### 1. Fix Tissue

Maintain frozen tissue on dry ice before and after weighing. Use a pre-chilled petri dish placed on ice while mincing the tissue.

a. 3 papillae are ~20-25mg.
b. Place the tissue on a pre-chilled glass petri dish maintained on ice and using a blade, mince tissue finely (enables passing through a 1 ml wide-bore pipette tip) in extra fixation buffer ~200 ul.
c. Aspirate the solution containing the minced tissue and transfer to a 2 ml low protein binding centrifuge tube. Using a wide-bore 1 ml pipette add 1 ml Fixation Buffer. Pass the tissue up and down. Maintain on ice.
d. Incubate for 16-24 h at 4 °C. DO NOT agitate or mix the sample during incubation. Fixation time and temperature should be consistent across all samples in an experiment. Our lab used 20 hours.
e. Centrifuge at 850 rcf (2900 rpm on lab centrifuge) for 5 min at room temperature.
f. Remove the supernatant without disturbing the tissue pellet.
g. Add 2 ml chilled PBS and resuspend the tissue pellet.
h. Centrifuge at 850 rcf (2900 rpm on lab centrifuge) for 5 min at room temperature.
i. Remove the supernatant without disturbing the tissue pellet.
j. Add 1 ml Quenching Buffer, resuspend the tissue pellet, and maintain on ice. Fixed tissue pieces can be stored at 4 °C for up to 1 week or at-80 °C for up to 6 months.
k. Centrifuge at 850 rcf / 2900 rpm (on lab centrifuge) for 5 minutes at room temperature.
l. Remove the supernatant without disturbing the fixed tissue pellet.
m. Proceed to tissue dissociation.

##### 2. Dissociate Fixed Tissue

a. Warm Dissociation Solution (Liberase TL used) for 10 minutes at 37 °C water bath before use.
b. Add 2 ml pre-warmed dissociation solution to the sample. Incubate for 20 minutes at 37 °C warm water bath. Shake the tube every 5 minutes. After 20 minutes, use a P1000 pipette tip and triturate the tissue pieces 20X (solution should turn slightly cloudy).
c. Pass the dissociated tissue through a 30 um strainer into a 5 ml conical to remove debris and undissociated tissue. Use the back of a 1 ml syringe plunger to push the pieces through the strainer.
d. Perform an additional wash of the 30 um filter by adding 2 ml PBS to the filter. Collect in the same tube.
    a. Centrifuge using swinging bucket rotor at 1800 rpm / 800 rcf (in tissue culture room) for 5 minutes.
e. Remove the supernatant without disturbing the pellet.
f. Resuspend pellet in 1 ml chilled quenching buffer.
g. Determine the cell concentration using a Countess III FL Automated Cell Counter (in Vantage core) using propidium iodide staining solution.
h. Ensure “PI stain” protocol selected
i. Select RFP under BF, then select Count
j. Expected value to be ~500,000 cells
k. Proceed immediately to Chromium Fixed RNA Profiling protocols or store the sample right after resuspending in appropriate reagents.

#### 3. Fixed Sample Storage Guidelines

*Short-term Storage at 4 °C (for fixed tissue pieces & cells)*

a. Thaw Enhancer for 10 minutes at 65 °C. Vortex and centrifuge briefly. Keep warm and verify no precipitate before use.
    i. DO NOT keep the thawed reagent on ice, or the solution will precipitate. Once thawed, Enhancer can be kept at 42 °C for up to 10 minutes.
b. Add 0.1 volume pre-warmed Enhancer to fixed cells in Quenching Buffer. For example, add 100 µl Enhancer to 1,000 µl fixed cells in Quenching Buffer. Pipette mix.
c. Store sample at 4 °C for up to 1 week.

*Long-term Storage at −80 °C (for fixed tissue pieces & cells)*

a. aThaw Enhancer for 10 minutes at 65 °C. Vortex and centrifuge briefly. Keep warm and verify no precipitate before use.
    i. DO NOT keep the thawed reagent on ice, or the solution will precipitate. Once thawed, Enhancer can be kept at 42 °C for up to 10 minutes.
b. Add 0.1 volume of pre-warmed Enhancer to fixed sample in Quenching Buffer. For example, add 100 µl Enhancer to 1,000 µl fixed sample in Quenching Buffer. Pipette mix.
c. Add 50% Glycerol for a final concentration of 10%. For example: add 275 µl 50% Glycerol to 1,100 µl fixed sample in Quenching Buffer and Enhancer. Pipette mix.
d. Store at −80 °C for up to 6 months. Storing fixed cells at −80 °C is recommended for best results.

#### 4. Fixed Cells – Post-Storage Processing

Samples may undergo a color change during storage (e.g. black, light gray, or green), however this will not impact assay performance.

If samples were stored at −80 °C, thaw at room temperature until no ice is present.

a. Centrifuge sample at 850 rcf for 5 minutes at room temperature.
b. Remove the supernatant without disturbing the pellet.
c. Resuspend cell pellet in 1 ml Quenching Buffer and maintain on ice.
d. Determine cell concentration of the fixed sample using an Automated Cell Counter (Countess III).
e. Proceed immediately to appropriate Chromium Fixed RNA Profiling protocols.

S. Figure 1

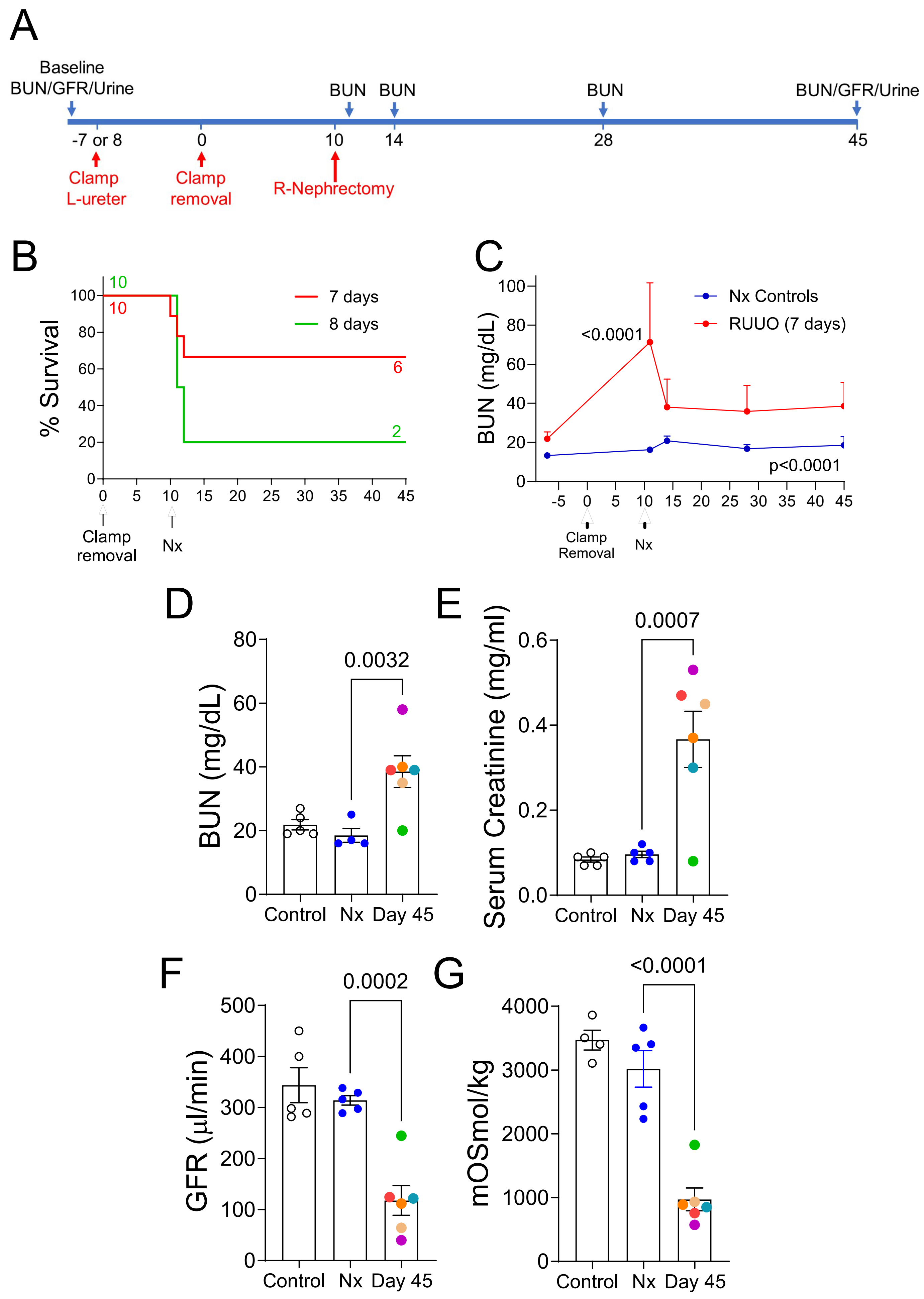


**S. Figure 1. Optimizing ureteric clamp times in male BALB/c mice to achieve long-term survival after R-UUO.** Male BALB/c mice underwent 7 or 8-day R-UUO and contralateral nephrectomy, or nephrectomy alone (Nx). **A,** Study design; **B,** Survival, numbers of mice indicated; **C,** BUN time course after 7-day R-UUO; **D-G,** BUN, serum creatinine, tGFR, and urine osmolality after 18hrs water restriction, in healthy controls (HC), Nx, and day 45 after 7-day R-UUO. **C**, BUN mean +/-SD. 2-way ANOVA. **D-G**, individual data points shown with mean +/-SEM. R-UUO datapoints are color coded to show the relationship between BUN, creatinine, tGFR, and urine osmolality in individual mice. 1-way ANOVA Nx vs. HC and R-UUO. If p<0.05, q values shown for between group comparisons corrected for repeat testing.

S. Figure 2

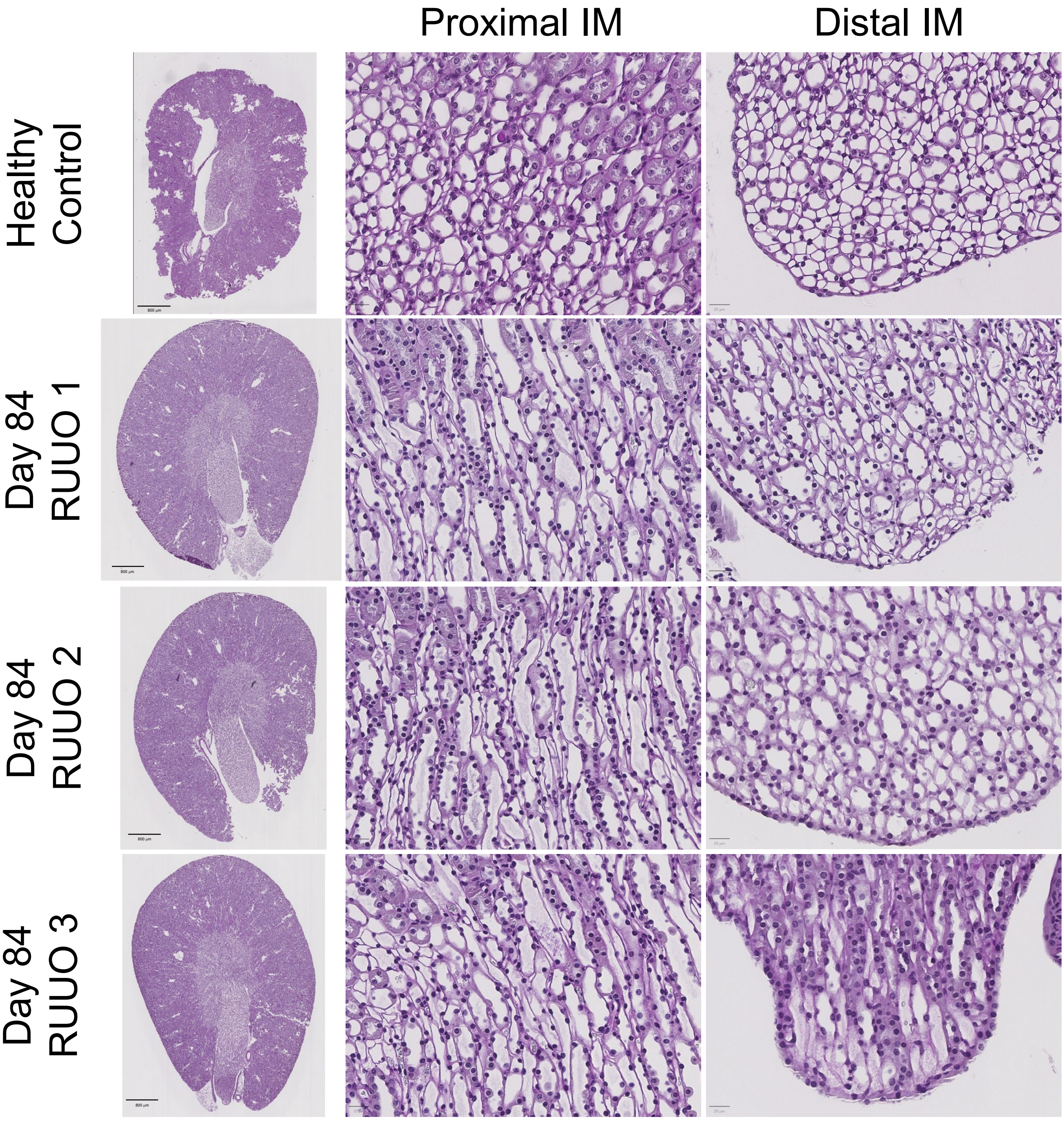


**S. Figure 2. Normal histological appearance of renal papillas 84 days after R-UUO.** PAS staining of a healthy control, and 3 Day 84 R-UUO kidneys scale bars=800um, proximal and distal inner medulla (IM), scale bars=20uM, as indicated.

S. Figure 3

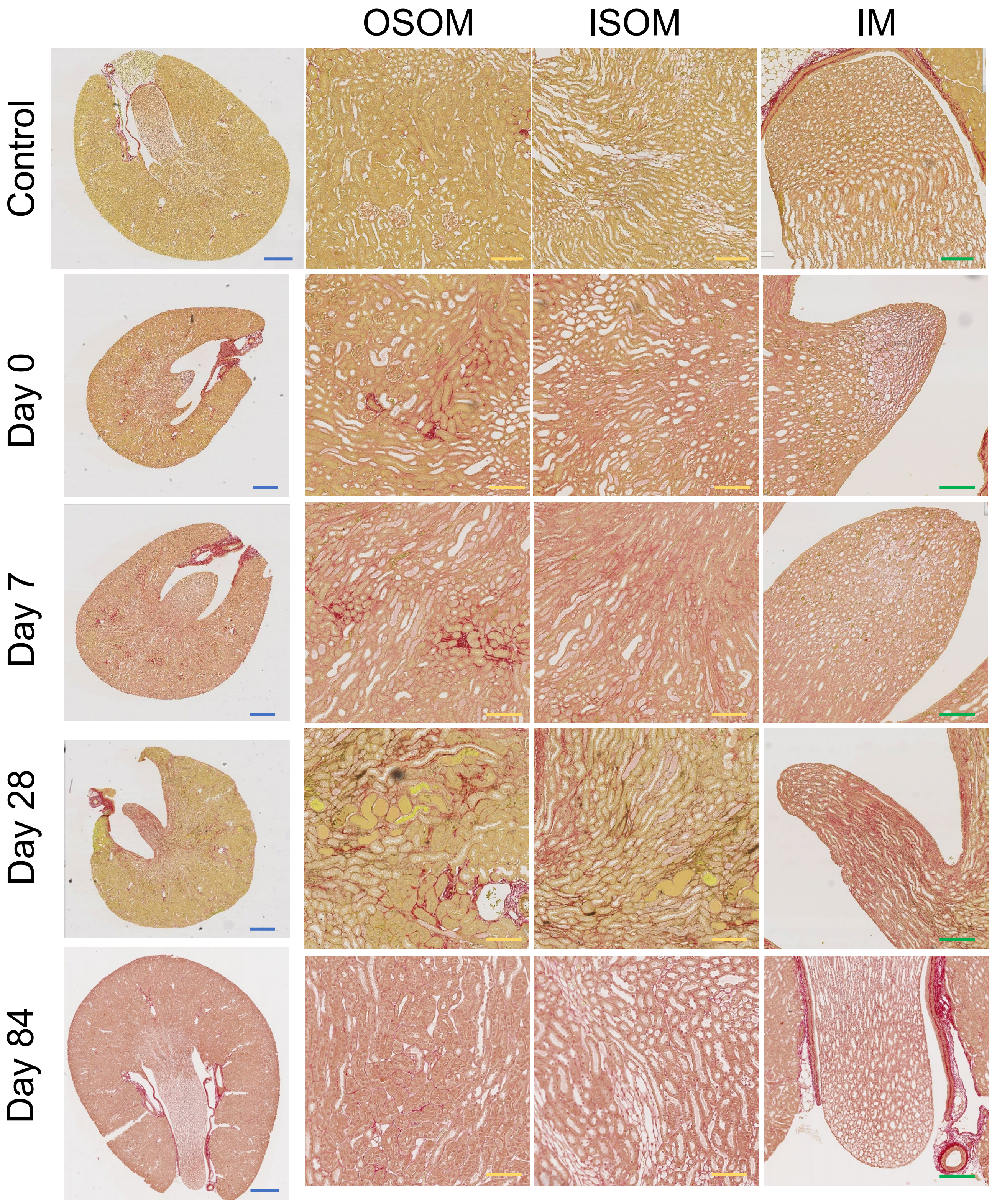


**S. Figure 3. Time course of collagen 1 staining in the inner and outer medulla after R-UUO.** Sirius red staining of type 1 collagen at different time points after R-UUO showing whole kidneys scale bars=800um (blue), ISOM and OSOM, scale bars=50um (yellow), inner medulla, scale bars=100um (green).

S. Figure 4

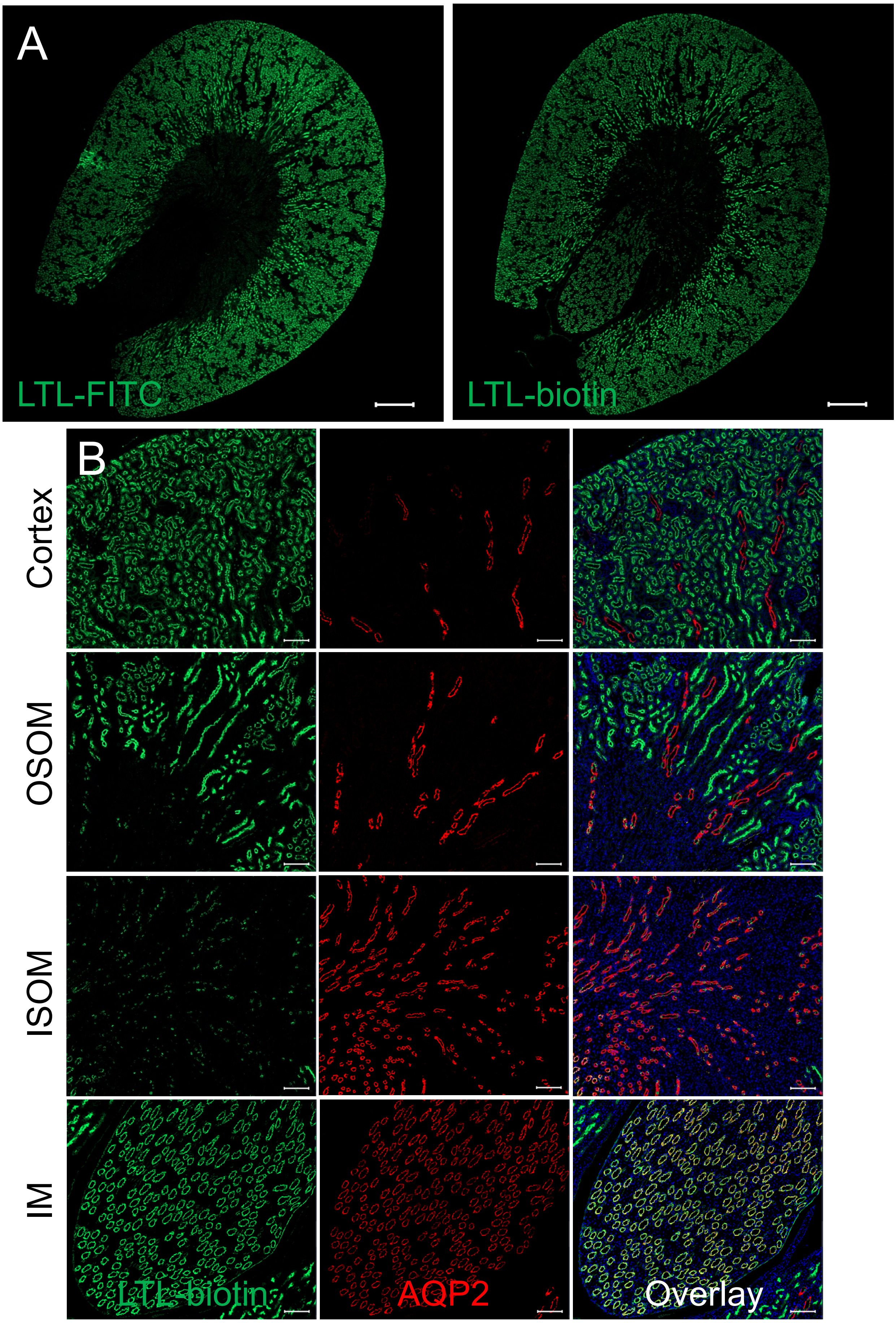


**S. Figure 4. LTL stains inner medullary collecting ducts. A,** Low power images comparing FITC-conjugated LTL and biotin conjugated LTL detected using neutravidin FITC amplification. In additional to widespread staining of proximal tubular epithelial cells throughout the cortex and OSOM, LTL-biotin amplification detects tubules in the inner medulla (IM). Scale bars=500uM ; **B,** Co-labeling LTL biotin with AQP2 showing co-localization of LTL staining with AQP2 in the IM. Additional punctate LTL staining of AQP2 negative collecting duct cells in the outer medulla and cortex. Scale bars=100uM.

S. Figure 5

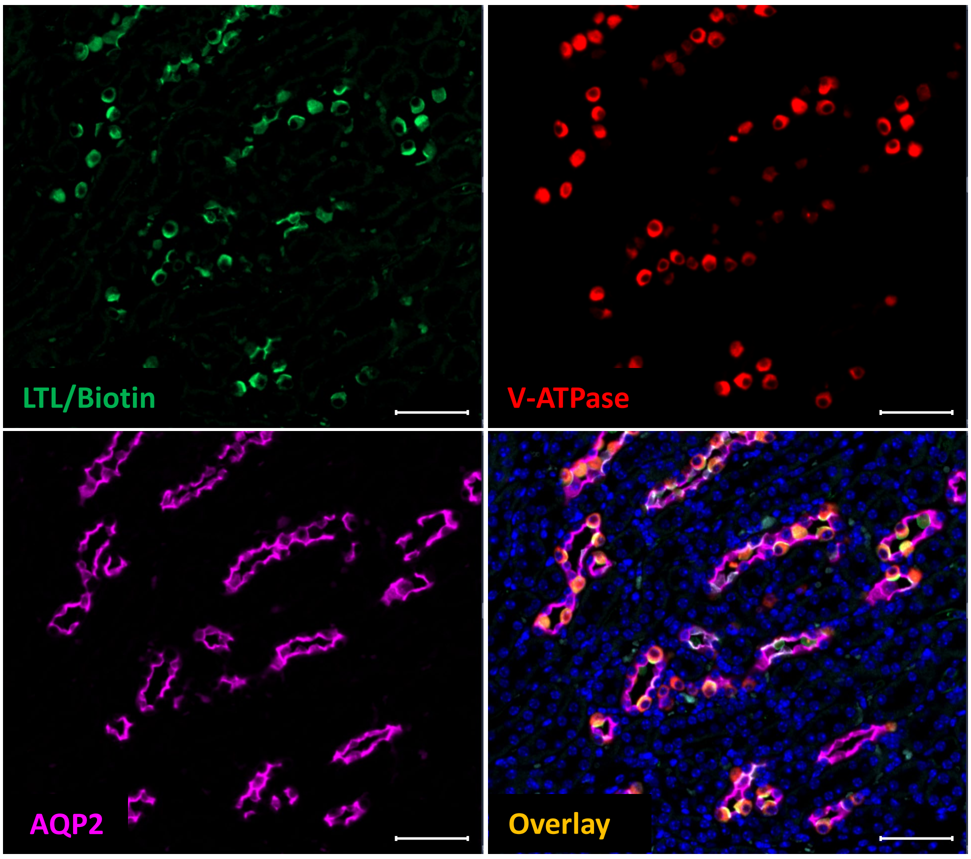


**S. Figure 5. LTL stains intercalated cells in the inner stripe of the outer medulla.** Co-labeling with LTL-biotin, AQP2 and V-ATPase B1/2 antibodies shows co-localization of LTL with V-ATPase staining in AQP2 negative V-ATPase positive intercalated cells in AQP2 positive collecting ducts seen in the ISOM. This is also seen in the outer stripe and cortical collecting duct intercalated cells. Scale bars=50uM.

### S. Figure 6

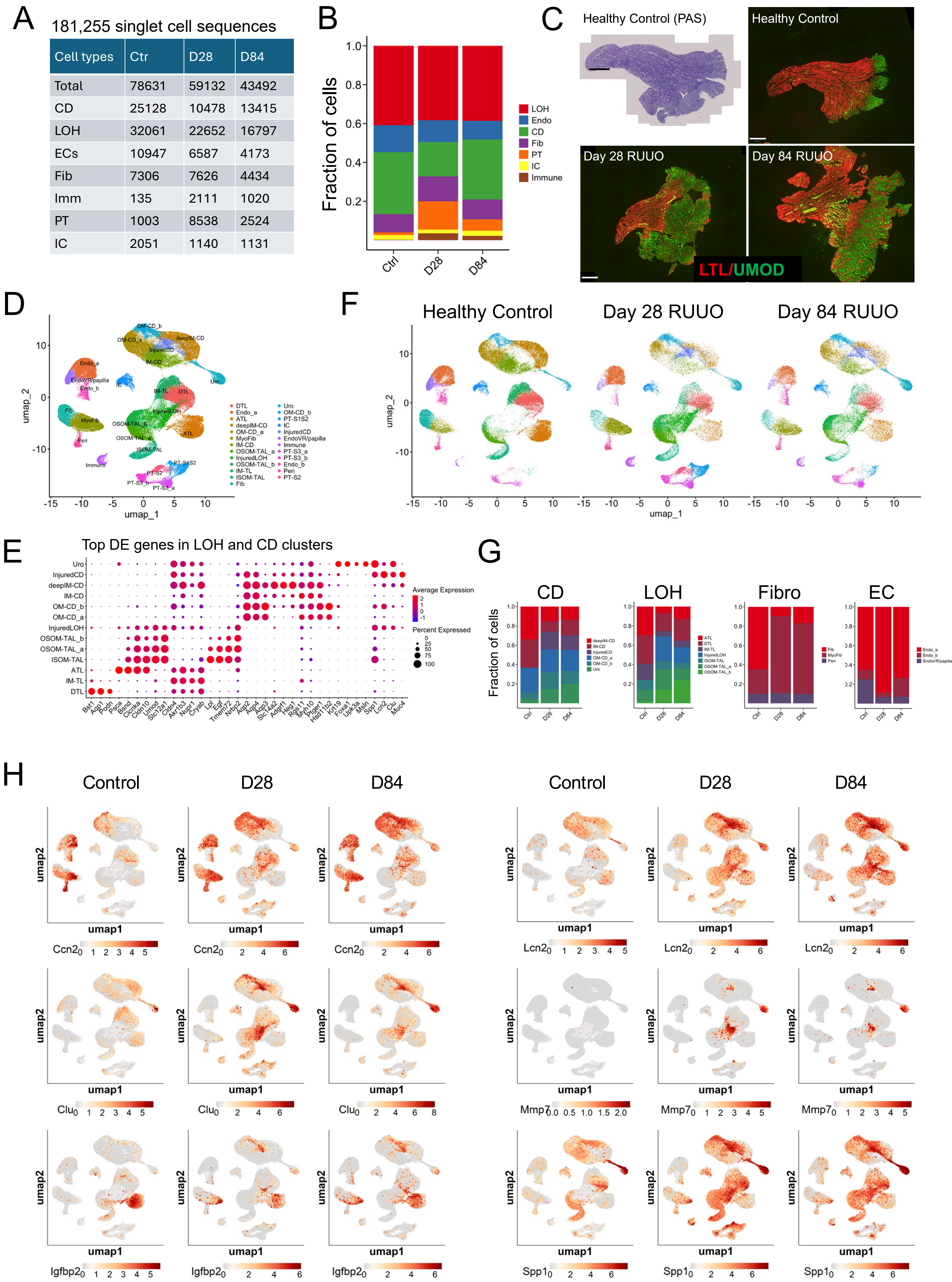


| Cell types | Ctr | D28 | D84 |
|---|---|---|---|
| Total | 78631 | 59132 | 43492 |
| CD | 25128 | 10478 | 13415 |
| LOH | 32061 | 22652 | 16797 |
| ECs | 10947 | 6587 | 4173 |
| Fib | 7306 | 7626 | 4434 |
| Imm | 135 | 2111 | 1020 |
| PT | 1003 | 8538 | 2524 |
| IC | 2051 | 1140 | 1131 |

**S. Figure 6. Single cell RNA sequencing atlas of mouse renal medullas after R-UUO.** We performed single cell RNA sequencing (scRNA-Seq) on isolated renal medullas dissected kidneys from multiple healthy controls and mice 28 and 84 days after R-UUO. **A,** 181, 255 single cell sequences were obtained from 7 major renal medullary cell populations; **B,** Fraction of each cell type in the combined scRNA-Seq data set; **C,** Renal medullas dissected from healthy controls, and days 28 and 84 after R-UUO stained with PAS, or with LTL and uromodulin antibody. Scale bars=500um. **D/F,** "Big" UMAP and identification of 25 different cell populations using published anchor genes in the combined data (D), and at different time points after R-UUO (F); **E,** Top DEGs in LOH and CD sub-clusters; **G,** Fraction of CD, LOH, fibroblast and endothelial cell (EC) sub-clusters at different time points; **H,** "Big" UMAPs showing the expression distribution of selected injury markers in different renal medullary cell populations from controls, day 28 and 84 after R-UUO .

### S. Figure 7

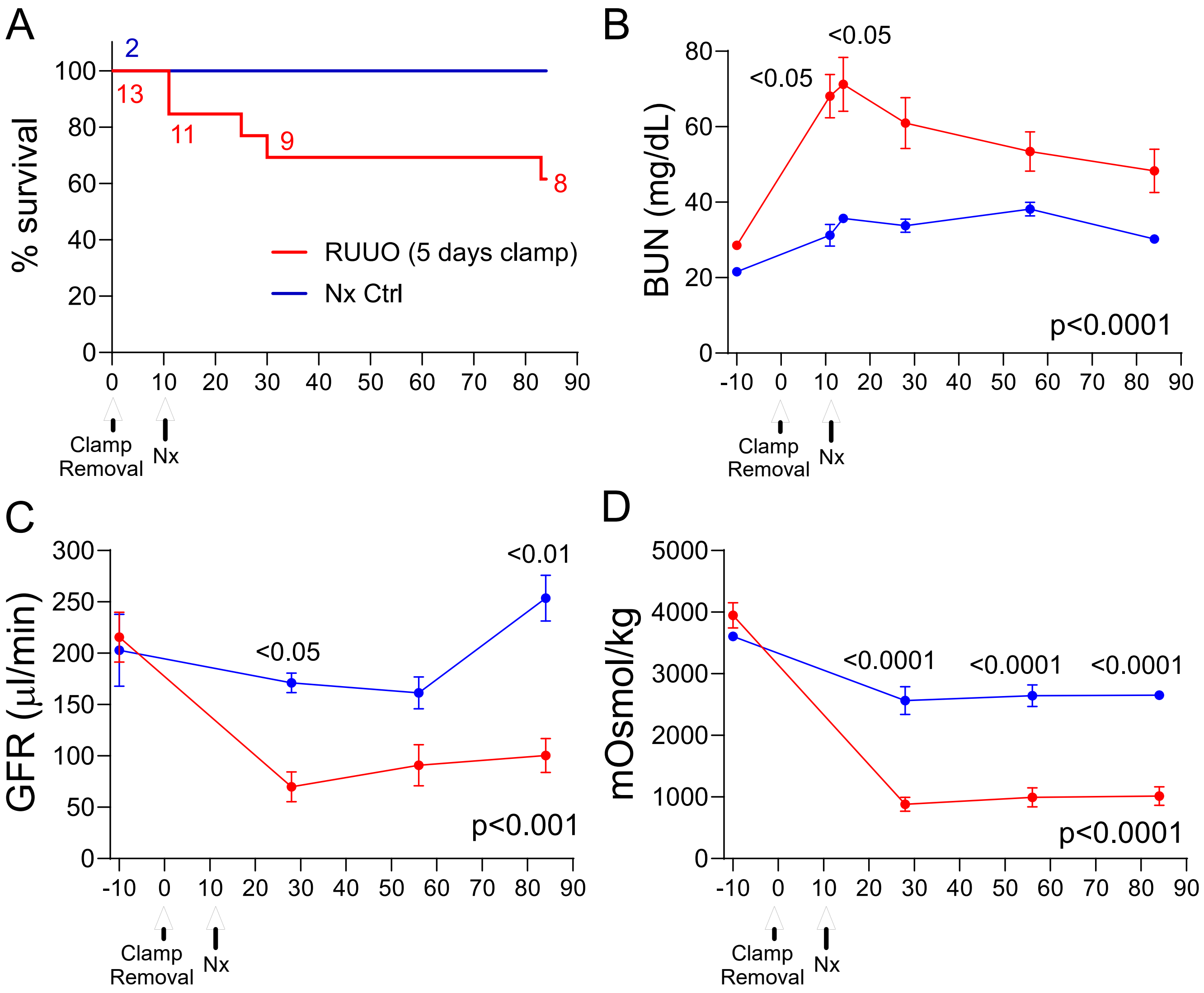


**S. Figure 7. Persistent defect in renal function and urinary concentrating capacity in Six2 Cre; tdTomato mice after R-UUO .** Male Six2 Cre; tdTomato mice (mixed background) underwent a 5-day R-UUO followed by contralateral nephrectomy, or nephrectomy alone (Nx). **A,** Survival, numbers of mice indicated; **B,** BUN time course after R-UUO; **C,** tGFR time course; **D,** Urinary osmolality after 18hr water restriction. Mouse numbers shown in A. Data shown as means +/-SEM. 2-way ANOVA p values indicated. If p<0.05, q values shown for between group comparisons corrected for repeat testing.

S. Figure 8

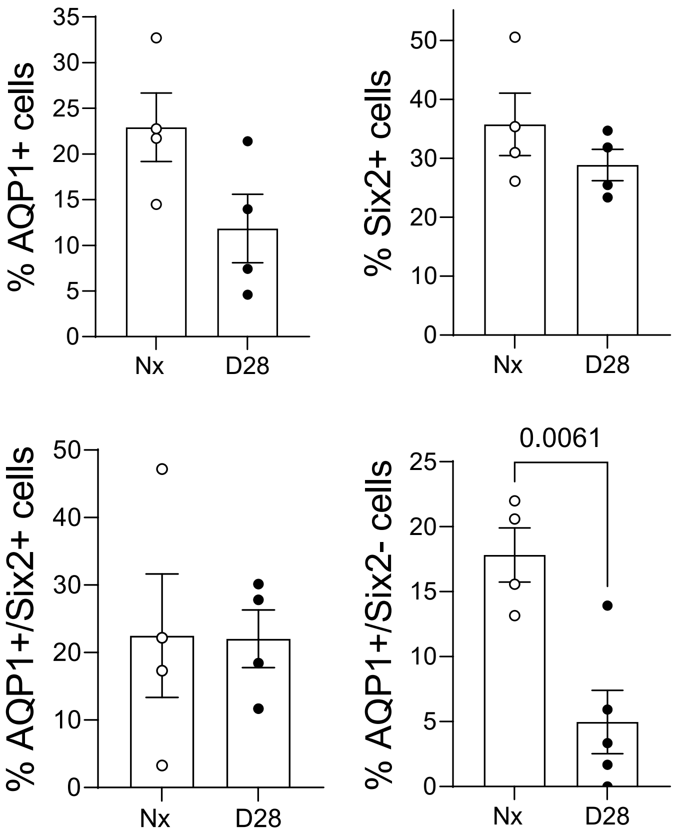


**S. Figure 8. Reduced numbers of Six2 lineage negative, AQP1 positive descending vasa recta cells in the proximal IM after R-UUO.** Quantification of Six2 lineage and AQP1 positive and negative cell populations in the proximal IM. Quantification as the % of total cells in each area. Individual data points, means +/- SEM. T-test, p values shown.

### S. Figure 9

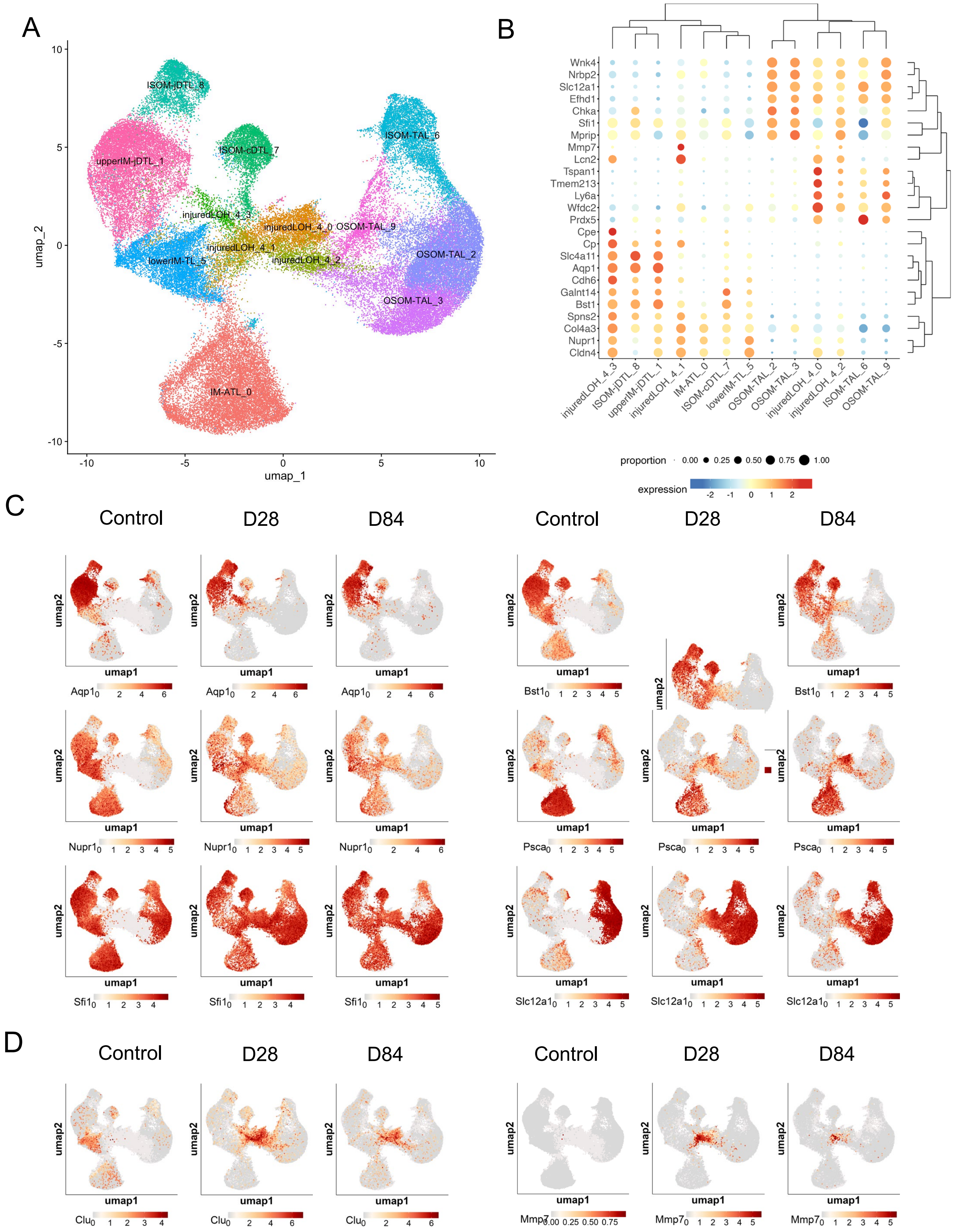


**S. Figure 9. Injured cluster 4 cells are derived from different inner and outer medullary loop of Henle cell populations. A,** Clustering of LOH cells identified 4 sub-clusters of injured LOH cluster 4 cells; **B,** Clustering of the top 25 LOH injured cluster 4 DEGs in other LOH cell populations in controls, day 28 and 84 after R-UUO; **C/D,** UMAPs showing changes in the distribution of LOH cell segment specific markers (C), and injury markers (D) in controls vs. day 28 and 84 after R-UUO.

### S. Figure 10

A

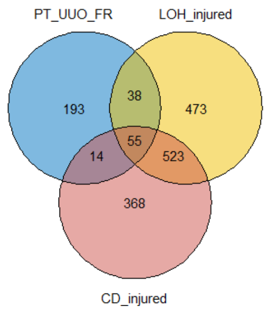


B

| Grp | Obs | Exp | P value |
|---|---|---|---|
| LOH/ CD | 578 | 54.9 | 0 |
| PT/ CD | 69 | 15.1 | 4.4E-27 |
| PT/ LOH | 93 | 17.1 | 1.9E-43 |
| PT/ LOH/ CD | 55 | 0.9 | 4.0E-81 |

C

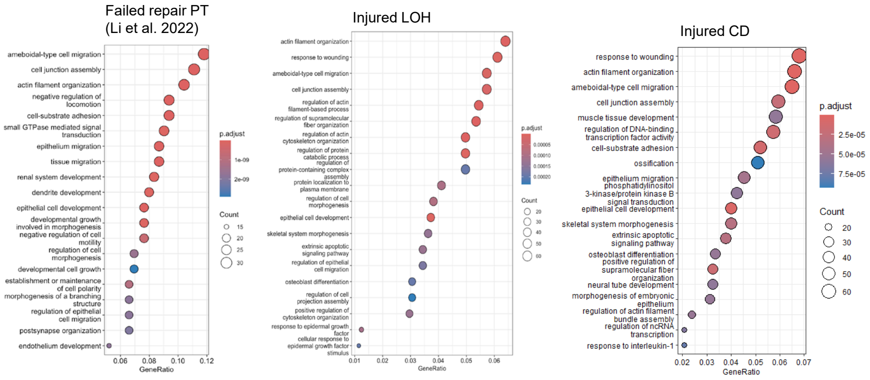


D

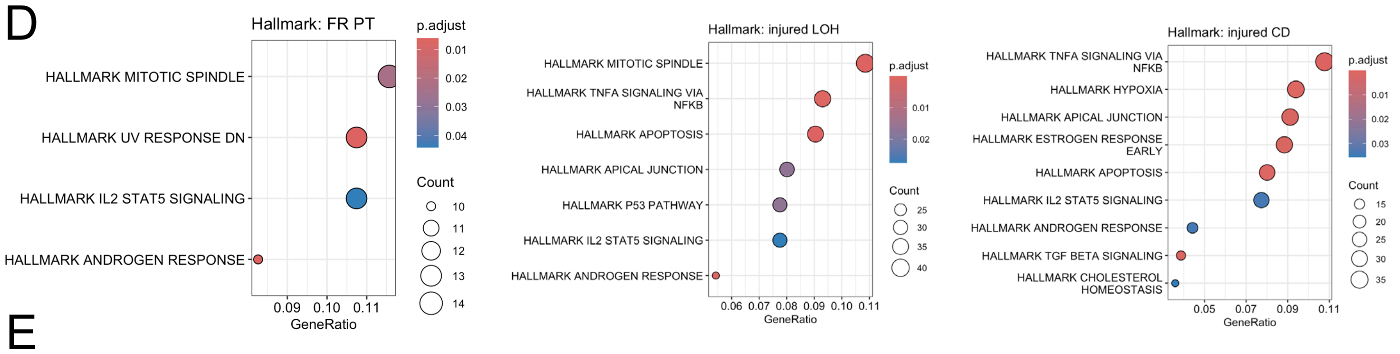


E

| Cluster | Set Size | NES | Adj. P | Core enrichment genes |
|---|---|---|---|---|
| Injured LOH | 277 | +2.03973735 | 4.27757660566617e-17 | C3/Vcam1/Zbtb7c/Icam1/Ly6e/Gxylt2/Csf1/Dpysl3/Havcr1/Kcnc2/Adamts1/Cd44/Sh2d4b/Srgap1/Dock10/Kcnh8/Cd74/Anxa3/Tsc22d1/Tbc1d1/Bmp6/Runx1/Fam129a/Ngf/Sox4/Spp1/Socs2/Trpm6/Relb/Col4a1/Sema6a/Thsd7a/Cadps2/Samd4/Cd9/Slc13a1/Mcc/Nfkbiz/Aoc1/Alpk1/Nt5e/Birc3/Pdgfb/Dennd2a/Gpc3/Sema3c/Tmem176b/Dcdc2a/Arid5b/Map3k1/Vmp1/Smurf1/Plxna2/Wipf1/Cttnbp2nl/Plscr1/Sbno2/Ezr/Klf6/Map4k4/Tpm1/Ccdc148/Cp/Nrg1/Dgkg/Myh9/Itgb8/Sytl2/Adgrl2/Ankrd6/Dlgap4/Etl4/Taok3/Tmem176a/Ypel2/Frk/Pdgfd/Gnai2/Il34/Ube2e2/Stat3/Ldlrad4/Itgav/Prickle2/Actn1/Flnb/Fmnl2/Znrf1/Ccdc88c/B2m/Cpe/Sh3d19/Slco3a1/Pawr/Foxp4/Tjp2/Tnip1/Igfbp7/Myo1e/Akt3/Map4/Agrn/Fat1/Nt5c2/Lama5/Arhgef17/Col18a1/Msn/Sash1/Osbpl3/Anxa2/Sept11/Tanc1/Wasf2/Adamts9/Cdk6/Akap13/Epn2/Gnb1/Plekha5/Nrp1/Cd2ap/Psd3/F11r/Spata6/Zeb1/Vcl/Sh3rf1/Tgfbr2/Map1b/Cadm1/Gnas |
| Injured CD | 269 | +2.44047034 | 1.19028354246677e-18 | Vcam1/C3/Zbtb7c/Pdgfb/Gxylt2/Pdgfd/Gpc3/Dennd2a/Akap12/Trpm6/Samd4/Msn/Dcdc2a/Cd44/Socs2/Xkr4/Sox4/Ly6e/Itgb8/Runx1/Sema6a/Map1b/Adamts1/Prickle2/Plat/Ldlrad4/Tsc22d1/Cttnbp2nl/Dock10/Cp/Dpysl3/Adamts9/Adgrl2/Ankrd6/Icam1/Csf1/Ube2e2/Akt3/Tmem176b/Itgb6/Map4k4/Cd74/Vmp1/Wipf1/Schip1/Osbpl3/Smurf1/Rims2/Tnik/Ccdc88c/Plxna2/Relb/Slc13a1/Ngf/Dlgap4/Sh2d4b/Sbno2/Myh9/Spns2/Ezr/Jazf1/Arid5b/Nfkbiz/Sytl2/Bicc1/Cdk6/Arhgef17/Cadps2/Gnai2/St6gal1/Vcl/Ppfibp1/Stat3/Nrg1/Mcc/Tjp2/Arl15/Itgav/Cobl/Taok3/Znrf1/Anxa2/B2m/Pawr/Anxa3/Spp1/Foxp4/Birc3/Frk/Map3k1/Col18a1/Tpm1/Agrn/Farp1/Tmem176a/Erbb4/Wasf2/Tnip1/Ptprg |

**S. Figure 10. Overlap between failed repair PTECs after I-UUO, and injured LOH and CD clusters after R-UUO. A,** Venn diagram illustrating the overlap between published FR-PTECs DEGs, and injured LOH cluster 4 and injured CD cluster 5 population DEGs from these studies; **B,** Statistical significance between multi-set intersections from the Venn Diagrams determined using the SuperExactTest function in R; **C/D,** GSEA of DEGs in FR-PTECs, injured LOH and CD populations showing upregulated GO terms (C), and Hallmark terms (D); **E,** GSEA of DEGs in injured LOH and CD populations with the DEG gene set from FR-PTECs showing set size, normalized enrichment scores (NES). Core enrichment genes indicate genes within the injured LOH and CD cluster DEGs that are driving the NES.

### S. Figure 11

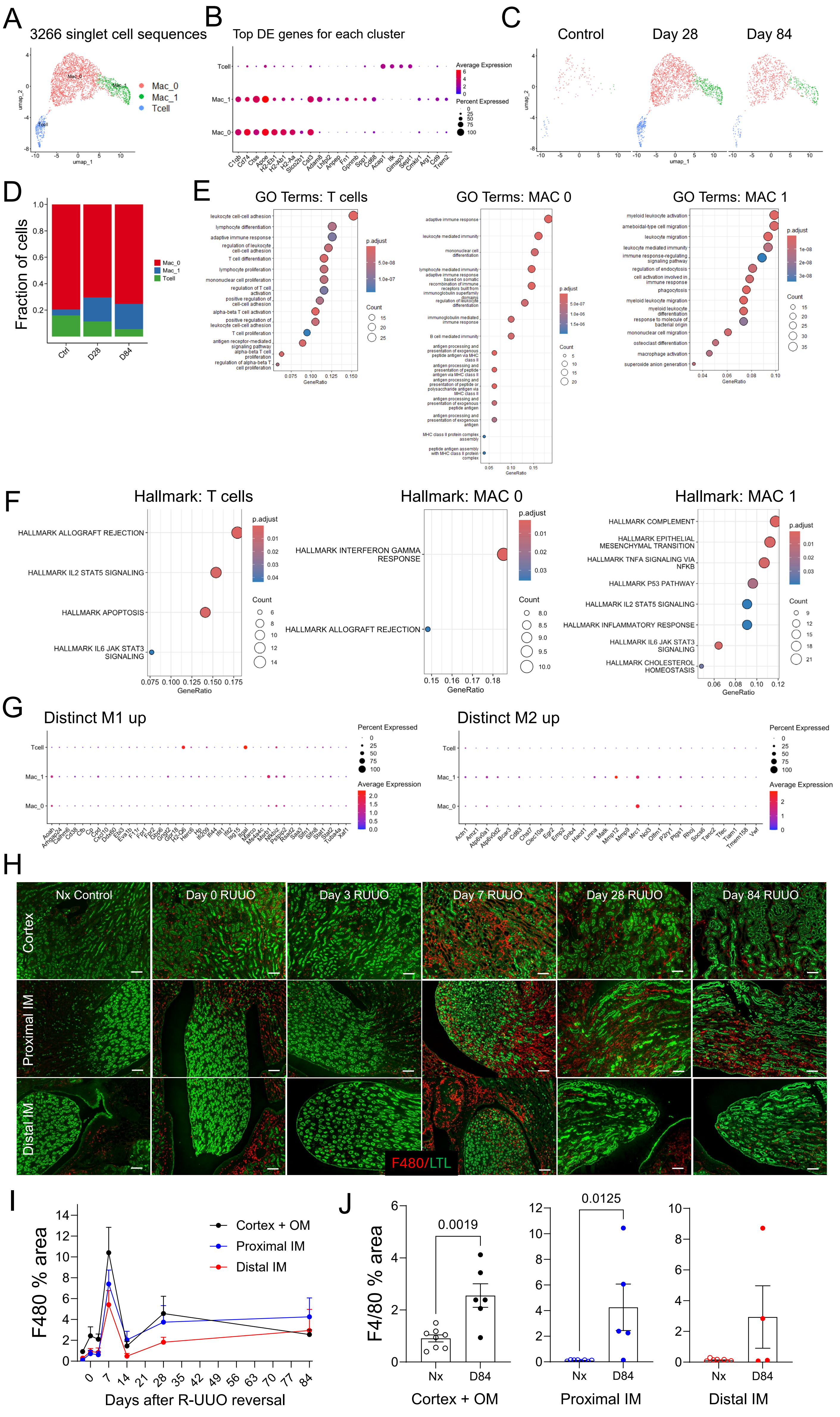
A
3266 singlet cell sequences
Mac_0
Mac_1
Tcell
umap_1
umap_2
B
Top DE genes for each cluster
Tcell
Mac_1
Mac_0
Average Expression
Percent Expressed
C1qb Cd74 Ctss Apoe H2-Eb1 H2-Ab1 H2-Aa Slco2b1 Csf3 Adam8 Lhfpl2 Anpep Fn1 Gpnmb Spp1 Cd68 Acap1 Itk Gimap3 Sept1 Cmklr1 Arg1 Cd9 Trem2
C
Control
Day 28
Day 84
D
Fraction of cells
Ctrl
D28
D84
E
GO Terms: T cells
GO Terms: MAC 0
GO Terms: MAC 1
GeneRatio
p.adjust
Count
F
Hallmark: T cells
HALLMARK ALLOGRAFT REJECTION
HALLMARK IL2 STAT5 SIGNALING
HALLMARK APOPTOSIS
HALLMARK IL6 JAK STAT3 SIGNALING
Hallmark: MAC 0
HALLMARK INTERFERON GAMMA RESPONSE
HALLMARK ALLOGRAFT REJECTION
Hallmark: MAC 1
HALLMARK COMPLEMENT
HALLMARK EPITHELIAL MESENCHYMAL TRANSITION
HALLMARK TNFA SIGNALING VIA NFKB
HALLMARK P53 PATHWAY
HALLMARK IL2 STAT5 SIGNALING
HALLMARK INFLAMMATORY RESPONSE
HALLMARK IL6 JAK STAT3 SIGNALING
HALLMARK CHOLESTEROL HOMEOSTASIS
GeneRatio
G
Distinct M1 up
Distinct M2 up
Percent Expressed
Average Expression
H
Nx Control
Day 0 RUUO
Day 3 RUUO
Day 7 RUUO
Day 28 RUUO
Day 84 RUUO
Cortex
Proximal IM
Distal IM
F480/LTL
I
F480 % area
Cortex + OM
Proximal IM
Distal IM
Days after R-UUO reversal
J
F4/80 % area
0.0019
0.0125
Nx
D84
Cortex + OM
Proximal IM
Distal IM

### S. Figure 12

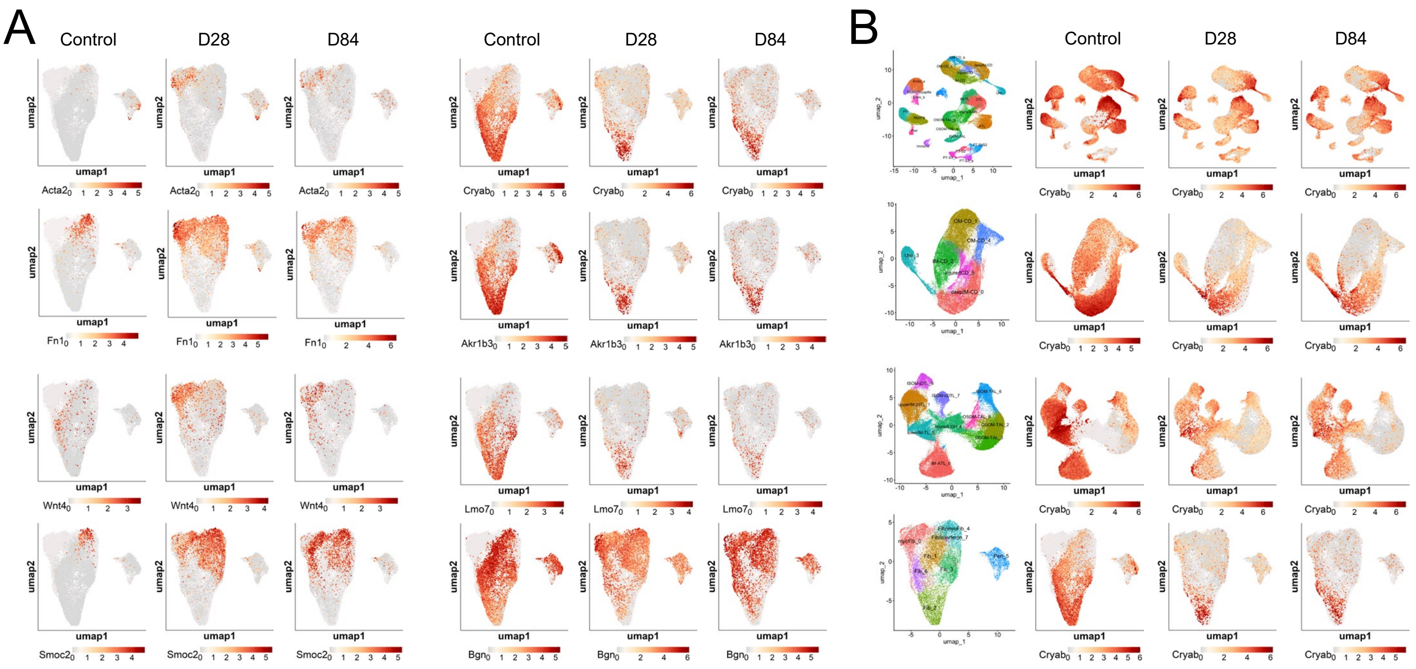


**S. Figure 12. Distribution of fibroblast and inner medullary cell marker genes.** UMAPs showing changes in the distribution of fibroblast and IM cell markers in fibroblast clusters (A), and of the IM cell marker, CryAB, in LOH, CD, and fibroblast clusters at different time points after R-UUO (B).

### S. Figure 13

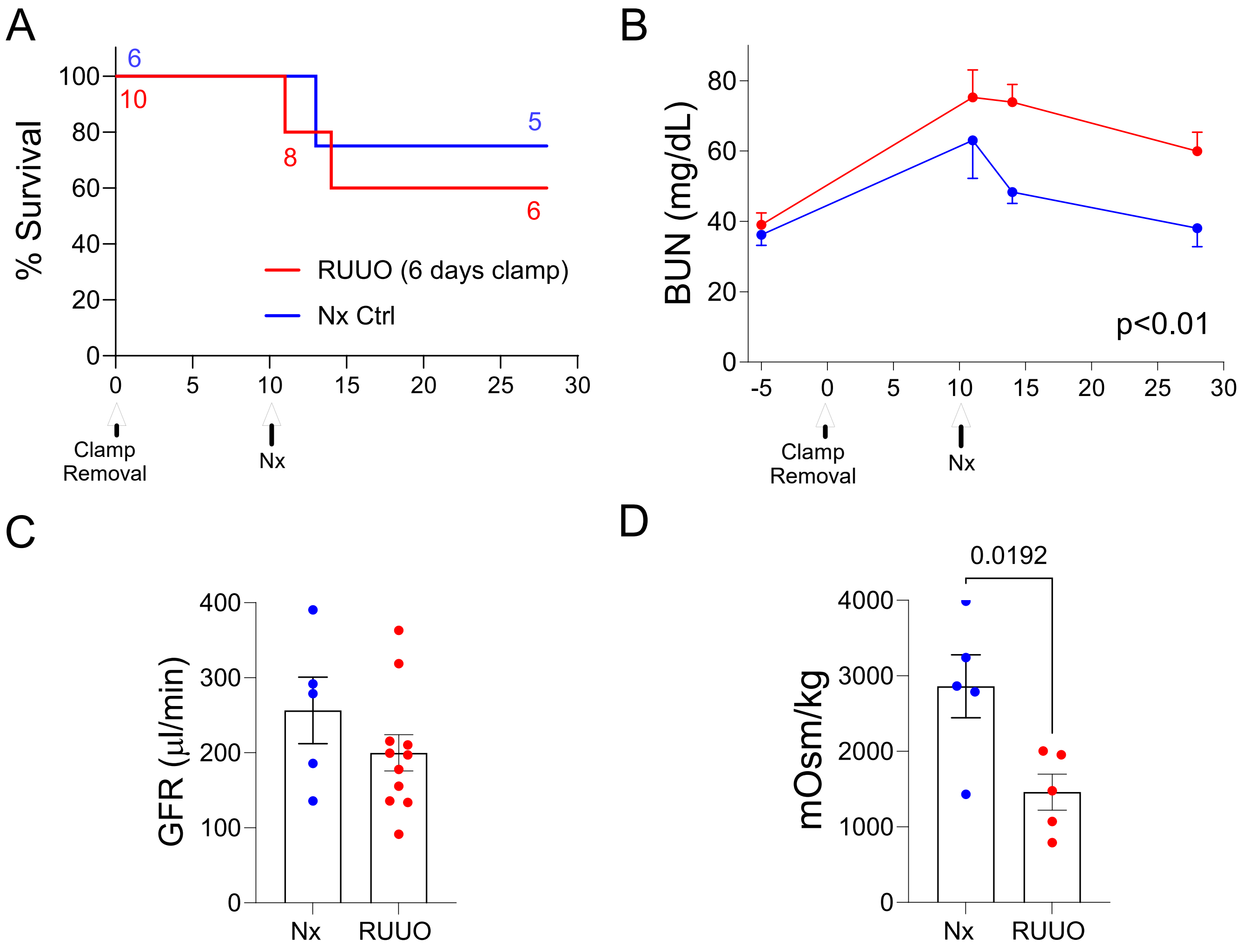


**S. Figure 13. Persistent defect in urinary concentrating capacity in TNC CreERT2; tdTomato mice 28 days after R-UUO.** Male TNC CreERT2; tdTomato mice (C57Bl/6 background) underwent a 5-day R-UUO followed by contralateral nephrectomy, or nephrectomy alone (Nx). **A,** Survival, numbers of mice indicated; **B,** BUN time course after R-UUO. Data shown means +/-SEM. 2-way ANOVA p values indicated; **C/D,** tGFR (C), and urinary osmolality after 18hr water restriction (D) 28 days after R-UUO, and in Nx controls. Means +/- SEM, individual data points shown. T test, p value shown.

S. Figure 14

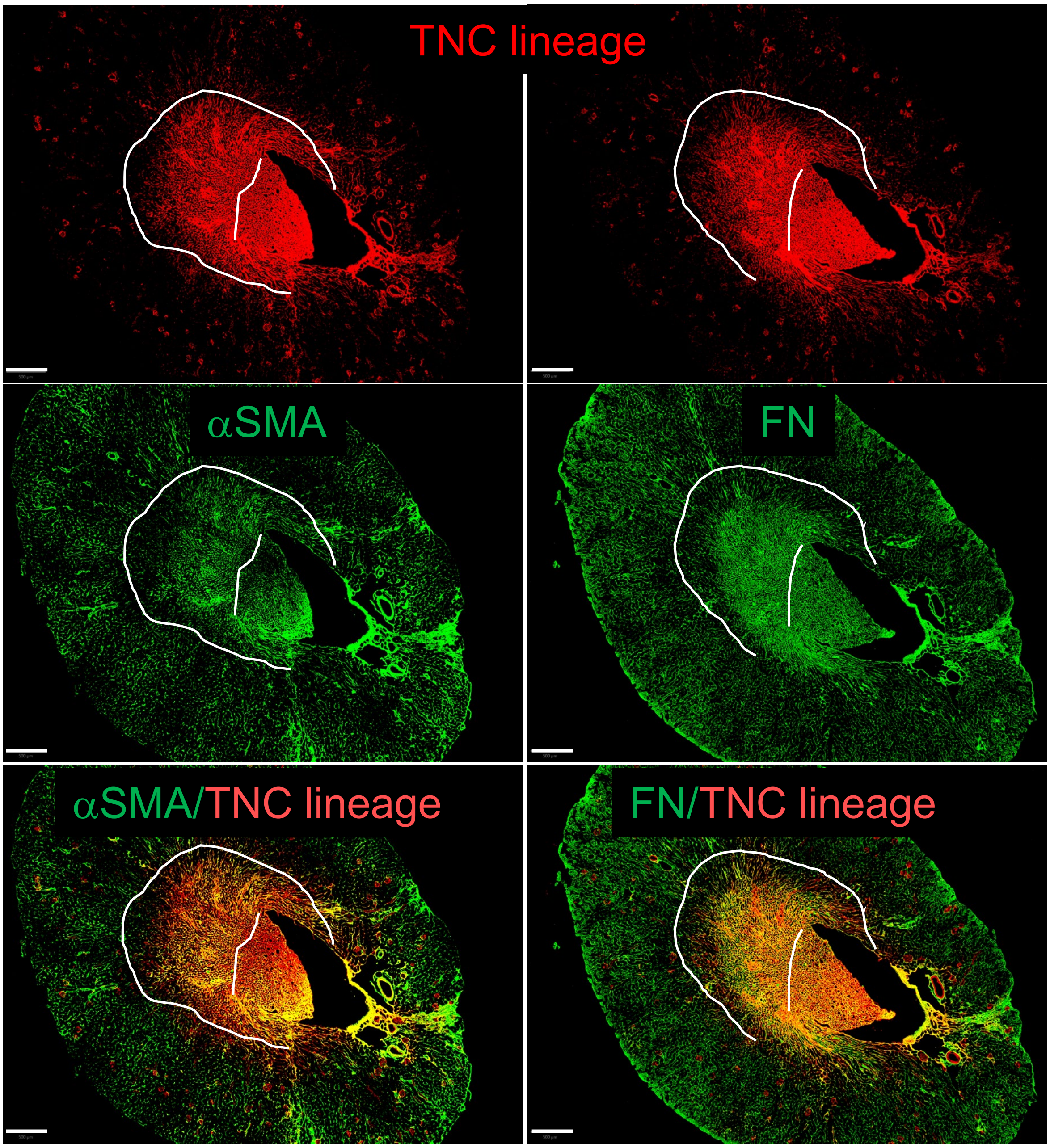


**S. Figure 14. Immunofluorescence images showing overlap between TNC lineage cells and myofibroblasts in the renal medulla.** Images showing TNC lineage (tdTomato), and α-SMA or Fibronectin (FN) staining on sequential sections of kidneys from a mouse 28 days after R-UUO. Scale bars=500um, dotted lines indicate the IM/ISOM boundaries.

S. Figure 15

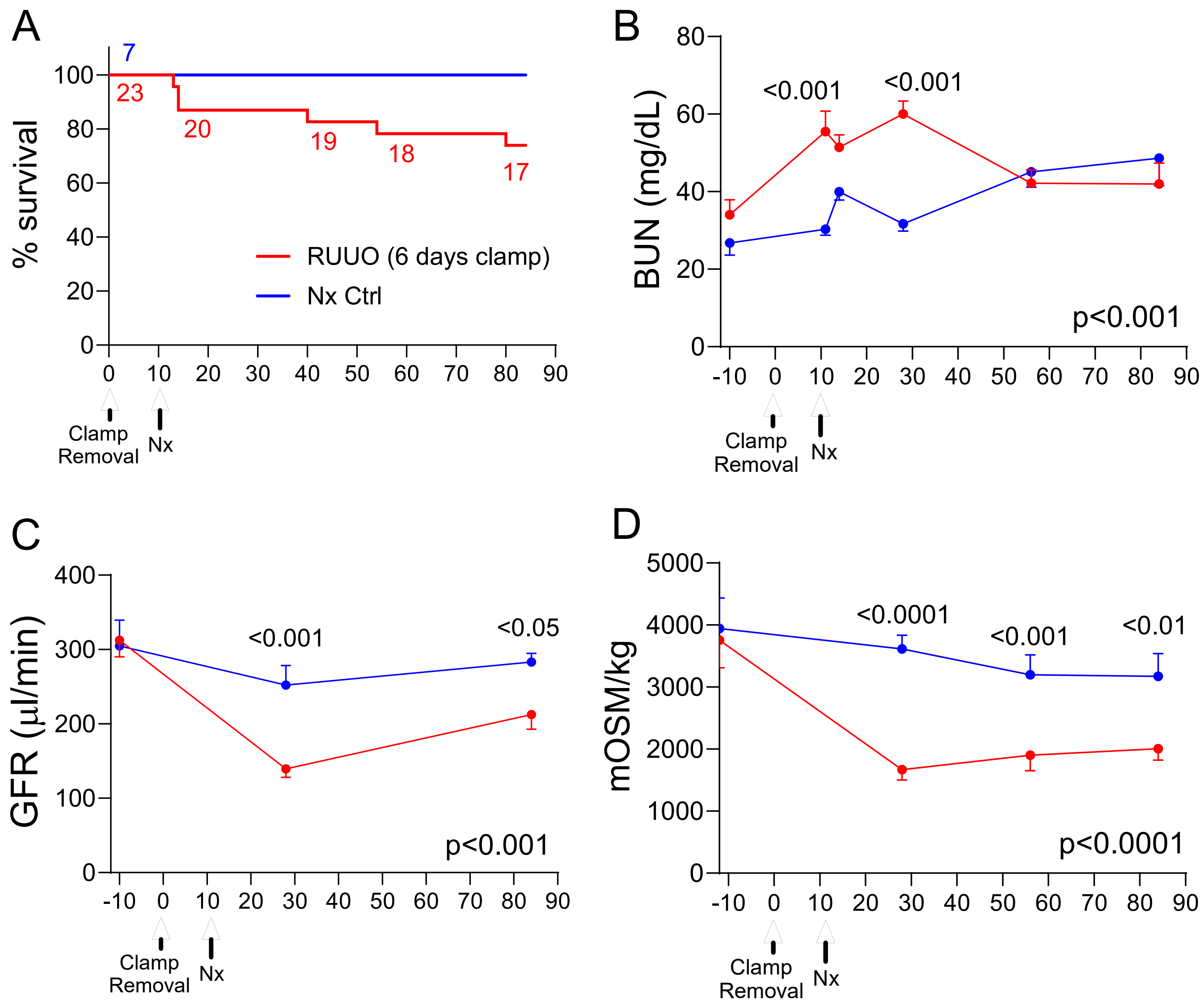


**S. Figure 15. Persistent defect in renal function and urinary concentrating capacity in HoxB7; tdTomato mice after R-UUO.** Male HoxB7 Cre; tdTomato mice (mixed background) underwent a 6-day R-UUO followed by contralateral nephrectomy, or nephrectomy alone (Nx). **A,** Survival, numbers of mice indicated; **B,** BUN time course after R-UUO; **C,** tGFR time course; **D,** Urinary osmolality after 18hr water restriction. Mouse numbers indicated in A. Data shown as means +/-SEM. 2-way ANOVA p values indicated. If $p<0.05$, q values shown for between group comparisons corrected for repeat testing.

### S. Figure 16

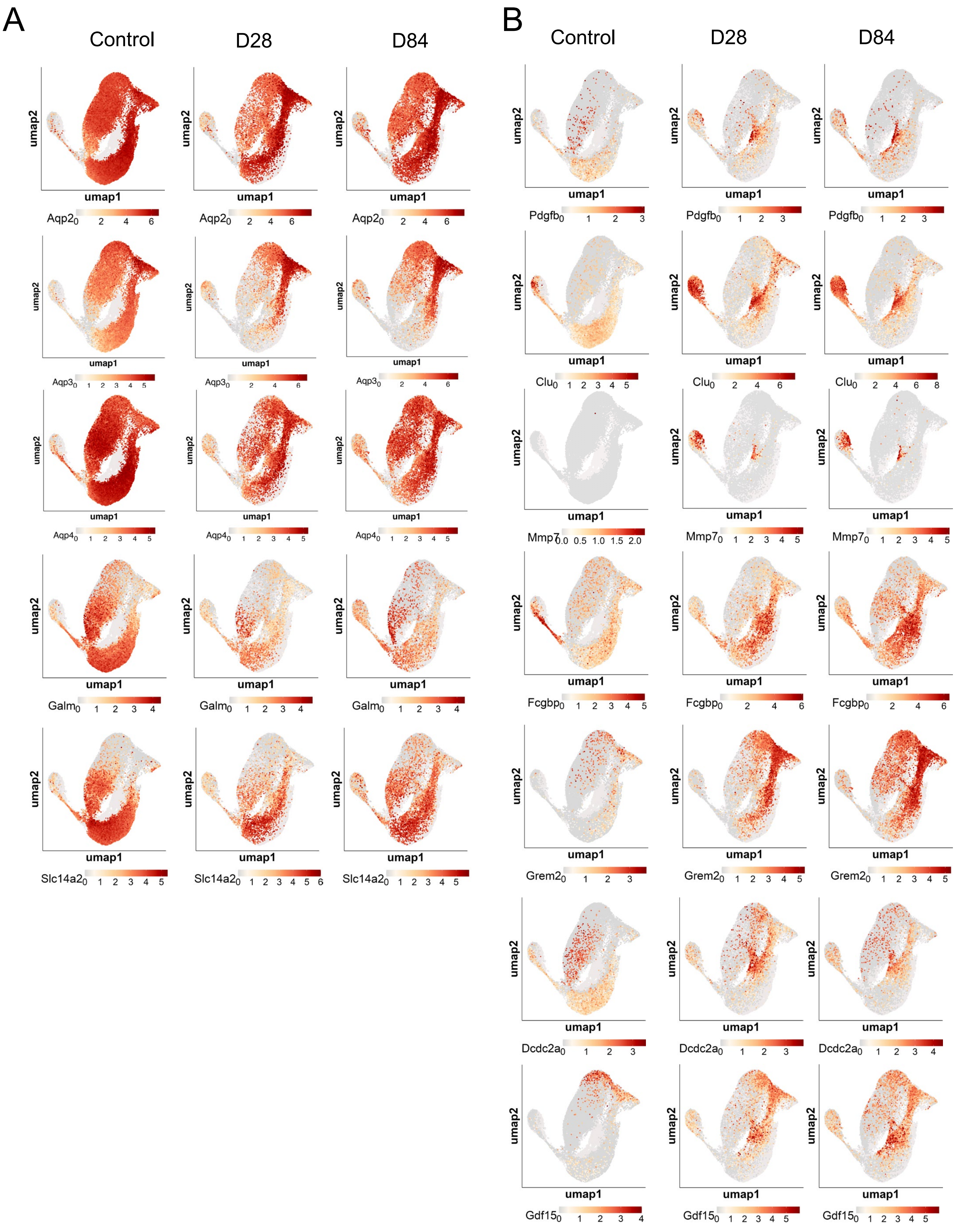


**S. Figure 16. Distribution of CD differentiation and injury markers after R-UUO.** UMAPs showing changes in the distribution of CD cluster differentiation (A), and injury (B) markers at different time points after R-UUO.

### S. Figure 17

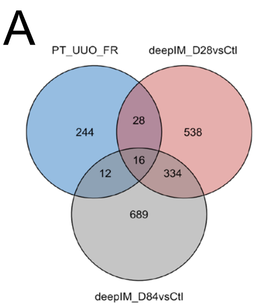


B

| Group | Obs | Exp | P value |
|---|---|---|---|
| FR PTEC/ D28 Deep IM CD | 44 | 14.4 | 3.77E-11 |
| FR PTEC/ D84 Deep IM CD | 28 | 16.5 | 0.005 |
| FR PTEC/D28/ D84 Deep IM CD | 16 | 0.79 | 3.23E-16 |

C

| Cluster | Set Size | NES | Adj. P | Core enrichment genes |
|---|---|---|---|---|
| Deep IM CD D28 | 196 | 1.639 | 0.000497 | Runx1/Akap12/Csf1/Spp1/Ly6e/Nfkbiz/Adamts9/Sh2d4b/Cobl/Relb/Tsc22d1/Igfbp7/Col18a1/Itga1/App/Pdgfb/Schip1/Foxp4/Smurf1/Dlgap4/Arhgef3/Plxna2/Ankrd11/Cttnbp2nl/Pawr/Mfhas1/Znrf1/Hivep2/Ldlrad4/Sash1/Klf6/Sbno2/Sytl2 |
| Deep IM CD D84 | 188 | 1.958 | 4.405E-07 | Akap12/Csf1/Ly6e/Spp1/Tsc22d1/Cobl/Nfkbiz/Itga1/Pdgfb/Fam129a/Pawr/Plxna2/Schip1/Ldlrad4/Klf6/Relb/Smurf1/St6gal1/Cttnbp2nl/Arhgef3/Foxp4/Mfhas1/Col18a1/Sbno2/Map4k4/Sash1/Frk/Dlgap4/Pkhd1/App/Stat3/Ezr/Sipa1l3/Osbpl3/Farp1/Nckap5/Ankrd11/Grk5/Hivep2/Zbtb16/Gnai2/Syne2/Znrf1/Wasf2/Mical3/Spns2/Trio/Epn2/Asap1/Agrn/Vmp1/Igfbp7/St5/Sytl2/Unc13b/Pard3b/Ptpn14/Etv6/Ypel2/Bicc1/Sh3rf1/Glis3/Adgrg1/Igsf5/Arid1b/Tjp2/Dnm3/Map3k1/B2m/Lpp |

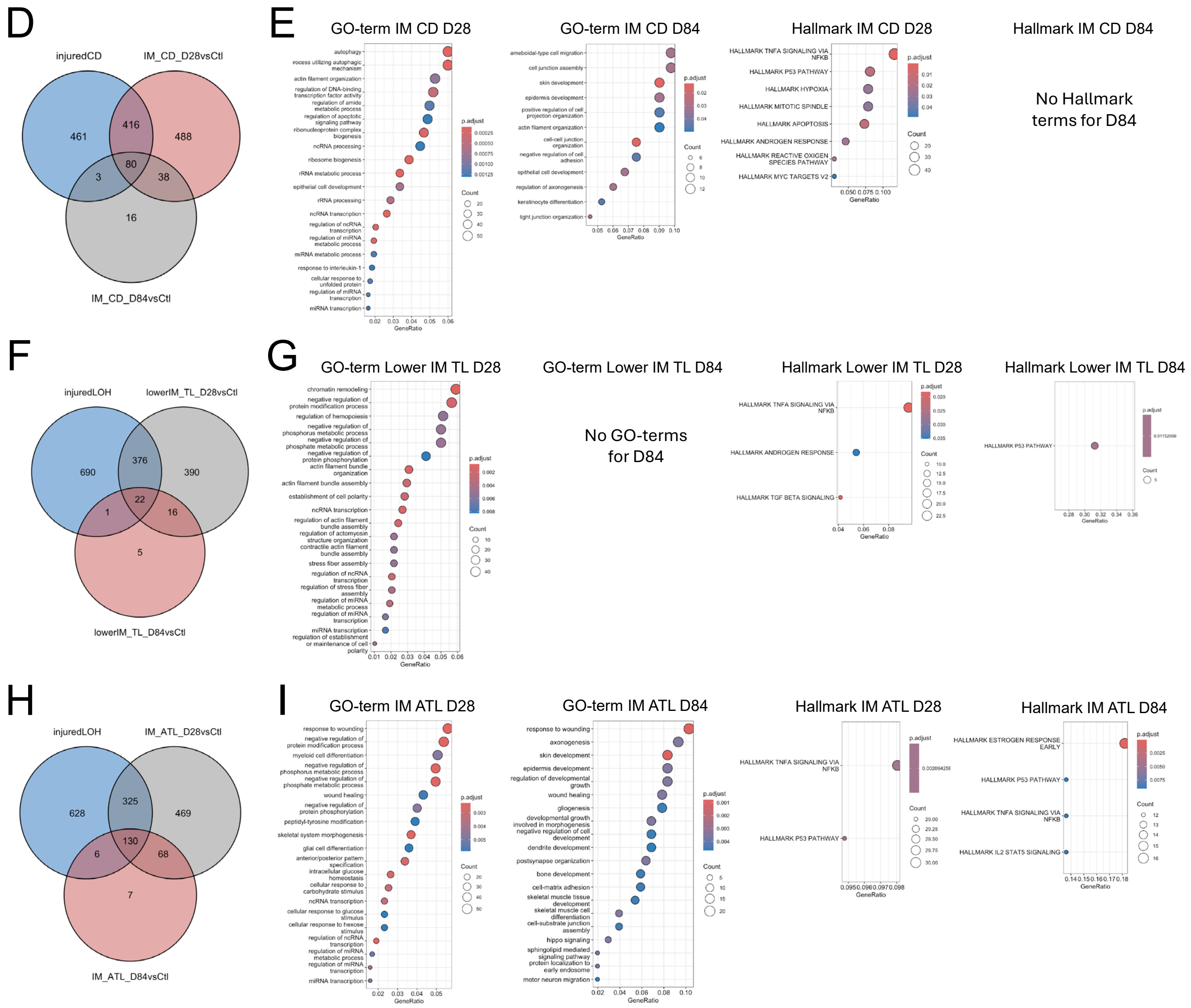


**S. Figure 17. Similarities and differences different CD and LOH populations after R-UUO. A-C,** Venn diagram showing the overlap between DEGs from FR-PTECs after I-UUO and deep IM CDs after R-UUO (A), statistical significance between multi-set intersections from the Venn Diagram (B), and GSEA of DEGs in Deep IM CD cells after R-UUO with the DEG gene set from FR-PTECs after I-UUO showing set size, normalized enrichment scores (NES). Core enrichment genes indicate genes within Deep IM CD day 28 and 84 cluster DEGs that are driving the NES; **D/F/H,** Venn diagrams showing the overlap between injured CD cluster 5 cells and IM CD cluster 2 cells after R-UUO (D), injured LOH cluster 4 and lower IM TL cells after R-UUO (F), and between injured LOH and IM ATL cells after R-UUO (H); **E/G/I,** statistical significance between multi-set intersections from the respective Venn Diagrams; **L/G/I,** GSEA showing upregulated GO and Hallmark terms in DEGs in IM CD cells (E), lower IM TL cells (G), and IM ATL cells (I) after R-UUO.

S. Figure 18

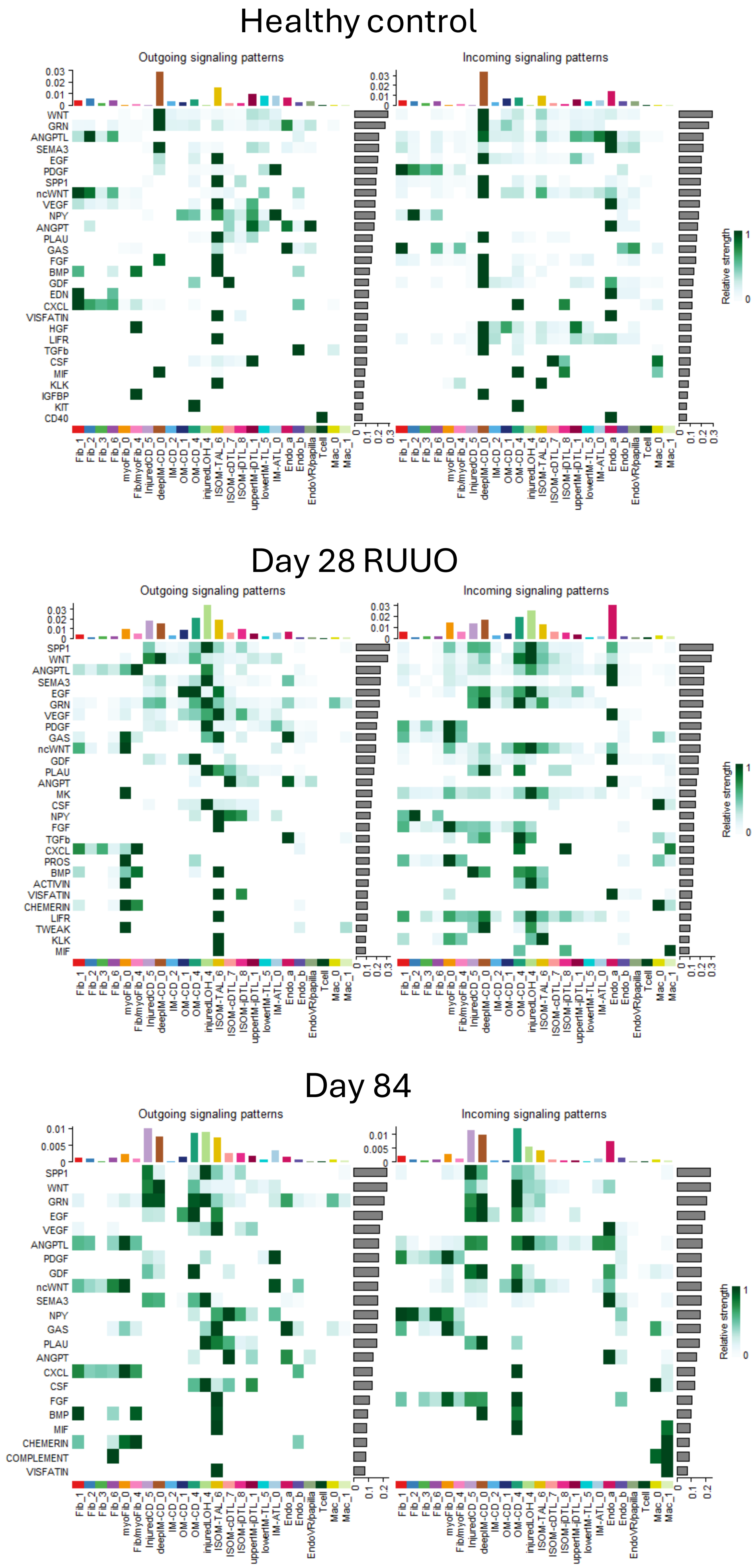


**S. Figure 18. Long-term effects of R-UUO on cell-cell communications in the renal medulla.** Cell-cell communication analysis was performed to identify ligand-receptor pairing of the major fibroblast, LOH, CD, endothelial and inflammatory cell clusters in renal medullas from controls, day 28 and day 84 after R-UUO. Outgoing ligand secretion is shown in the left-hand panels, incoming cognate receptors for the same pathway is shown in the right-hand panels. Contribution of individual cell clusters to the signaling patterns is indicated by the horizontal bar charts, relative strength of the response indicated with color coding and from the vertical bar charts.
